# Reuse of an insect wing venation gene-regulatory subnetwork in patterning the eyespot rings of butterflies

**DOI:** 10.1101/2021.05.22.445259

**Authors:** Tirtha Das Banerjee, Antónia Monteiro

**Affiliations:** Department of Biological Sciences, National University of Singapore, Singapore - 117557; Science Division, Yale-NUS College, Singapore - 138527

**Keywords:** Gene-subnetwork co-option, venation patterning, *Optix*, *decapentaplegic*, *spalt*, eyespots, *Bicyclus anynana*

## Abstract

Novel organismal traits might reuse ancestral gene-regulatory networks (GRNs) in their development, but data supporting this mechanism are still sparse. Here we show the reuse of an ancestral insect venation gene regulatory subnetwork patterning the sharp and distinct rings of color in butterfly eyespots. Using laser microdissection followed by RNA-Seq we first obtained transcriptional profiles of the anterior and posterior compartment of larval wings, and eyespot and adjacent control tissue in pupal wings of *Bicyclus anynana* butterflies. We identified key venation patterning genes such as *Mothers against dpp 6 (Mad6), thickveins, Optix, spalt, optomotor-blind (omb), aristaless, cubitus interruptus,* and *patched* differentially expressed (DE) across compartments, and a sub-set of these genes also DE across eyespot and non-eyespot samples. Fluorescent *in-situ* hybridization (HCR3.0) on the jointly DE genes *Mad6, Optix,* and *spalt*, as well as *dpp* showed clear eyespot-center, eyespot-rings, and compartment-specific expression. Knocking out *dpp* resulted in an individual with venation defects and loss of eyespots, whereas knockouts of *Optix* and *spalt* resulted in the loss of orange scales and black scales, respectively. Furthermore, using CRISPR-Cas9 followed by immunostainings, we showed that Spalt represses *Optix* in the central region of the eyespot, limiting *Optix* expression to a more peripheral ring, which parallels the regulatory interaction found in venation patterning in the anterior compartment of fly larval wings. These network similarities suggest that part of the venation GRN was co-opted to aid in the differentiation of the eyespot rings.

**One-sentence summary:** We showed the reuse of an ancestral insect wing venation GRN in patterning a novel complex trait in butterflies.

## Introduction

While it is possible that novel organismal traits emerge from the gradual construction of dedicated gene-regulatory networks (GRNs), one gene being added at a time, such networks may instead emerge from pre-existing clusters of pre-wired genes that are repurposed during development. So far, a few examples of this mechanism of GRN co-option have been proposed for the development of novel complex traits. These include the reuse of a GRN that differentiates breathing spiracles in *Drosophila* larvae in the development of the posterior lobe of adult genitalia (Glassford et al., 2015); the reuse of the leg GRN in patterning the wings of insects (Bruce and Patel, 2020; Clark-Hachtel and Tomoyasu, 2020) or the head horns of beetles (Hu et al., 2019; Zattara et al., 2016); the reuse of the wing GRN in the development of a treehopper’s helmet (Fisher et al., 2020); or the reuse of an appendage GRN to pattern butterfly eyespot centers (Murugeshan et al., 2022).

Eyespots, unlike the other examples of appendage co-option, where the novel trait protrudes from the body, do not grow out of the wing (Monteiro, 2015). This means that either only a small core network of genes shared between appendages and eyespots was co-opted in the first place, or that the similarities in the development of these traits break apart later in development, after the eyespot centers have differentiated in the late larval stage. One mechanism for this to happen is if another network gets rewired downstream of the shared core network to produce a novel output, i.e., to differentiate rings of color around the central signalling cells rather than tissue outgrowth. Here we explore whether the network of genes involved in early vein patterning in butterflies could have been repurposed to differentiate the color rings in eyespots at later stages of development.

The mechanism of wing vein patterning during larval development in *Bicyclus anynana* butterflies has many similarities to that of *Drosophila*, which is a classic example of positional information in developmental biology (Blair, 2007) (**Figure S1**). In both flies and butterflies, the morphogen Decapentaplegic (Dpp) is expressed in a stripe in the middle of the larval wing, along the Anterior-Posterior (AP) compartment boundary (Banerjee and Monteiro, 2020; Lawrence and Struhl, 1996). In flies, Dpp activates the transcription factors *spalt* and *Optix* (the latter expressed only in the anterior compartment), at different concentration thresholds and distances from the AP boundary (Martín et al., 2017). Subsequently, the cross-regulatory interaction between these two genes determines a sharp boundary that eventually positions longitudinal vein 2, L2 (Martín et al., 2017) (**Figure S1**). In butterflies, the expression of Spalt and Optix proteins is also found straddling a stripe of *dpp* expression, in homologous patterns to those in flies (Banerjee and Monteiro, 2020), suggesting a conserved vein patterning mechanism in both insects at early stages of wing development.

At later stages of wing development, and in butterflies alone, a similar mechanism of positional information has been hypothesized to specify the multiple rings of color in species with eyespots. Here, instead of an AP boundary line source, a central morphogen secreted by the eyespot centers would activate two or more transcription factors at decreasing concentration thresholds and distances from the center (Brunetti et al., 2001; Monteiro et al., 2006; Nijhout, 1978). The similarities between eyespot development and vein patterning, however, rest solely on the expression of Spalt protein in the central black scales of the eyespot (Brunetti et al., 2001), as *Optix, dpp* and other venation candidate genes have never been tested in and around the eyespots during the pupal stage.

Here we tested the hypothesis that a part of the vein patterning GRN, involving the ligand Dpp, the candidate transcription factors Spalt and Optix, as well as other vein-patterning genes, might have been co-opted in the differentiation of the eyespot color rings. We first identified differentially expressed (DE) genes between the anterior and the posterior compartments in mid larval wings, followed by DE genes between pupal eyespot and control tissue in 18-22 hrs pupal wings. We then examined the expression of several candidate Dpp pathway members as well as *spalt* and *Optix,* using enzyme and fluorescent based in-situ hybridization, and tested their function using CRISPR-Cas9. Finally, we targeted both *spalt* and *Optix*, in turn, and followed these disruptions with visualizations of the presence of both proteins in the eyespot field using immunofluorescence, to test for a potential functional interaction in color ring differentiation similar to that observed in L2 vein differentiation in *Drosophila* (Martín et al., 2017).

## Results

### Differential gene expression between anterior and posterior compartments of larval wings

To identify and confirm differentially expressed genes between the anterior and the posterior compartments of *B. anynana* mid-larval wings, we used RNA extracted from laser-based microdissections followed by RNA-Seq and DESeq2 analyses (**Figure 1A, Figure S2**). Character-tree analysis showed that the biological replicates from the anterior and the posterior compartment clustered together (**Figure 1B**). Genes that were significantly up-regulated in the anterior compartment included *aristaless, aristaless-like, cubitus-interruptus, patched, Optix,* and *dachshund* (**Figure 1D, Table S1**), whereas posterior compartment up-regulated genes included *engrailed-like, invected-like, hedgehog, hexamine-like, caupolican-like,* and *Toll-like receptor 6* (**Figure 1D, Table S1**). These data are consistent with studies on anterior and posterior compartment-specific genes in insects (Abouheif and Wray, 2002; Banerjee and Monteiro, 2020; Biehs et al., 1998; Bier, 2000; Blair, 2007; Martín et al., 2017).

**Figure 1.**
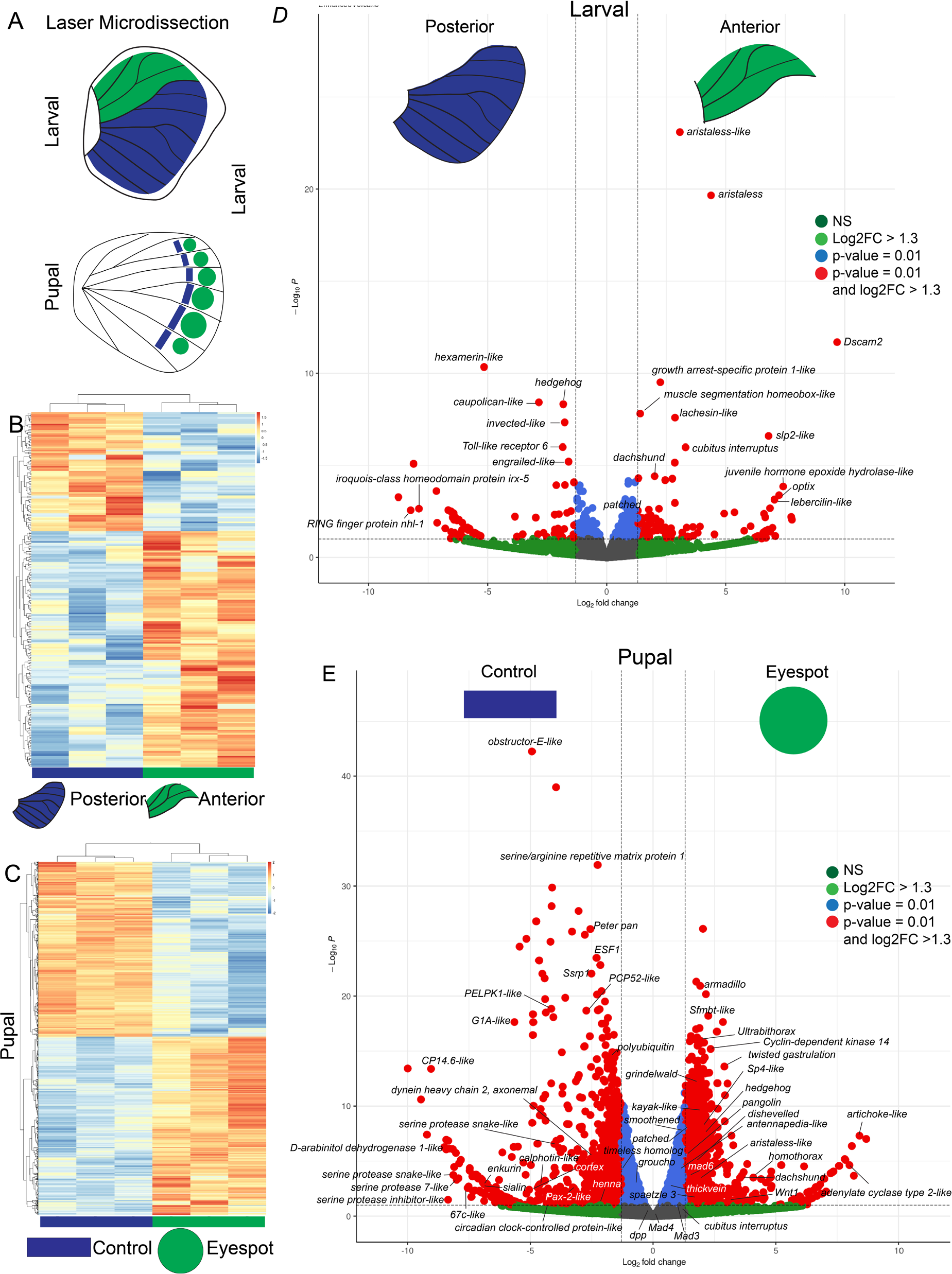
RNA-Seq of the larval anterior and posterior compartment and of pupal eyespot and control tissues dissected with a laser. **(A)** Laser microdissection was performed to isolate the anterior and posterior compartment of larval wing discs, and the eyespot and adjacent control tissue in 18-22 hrs pupal wings. **(B)** Heatmap clustering of the biological replicates of the anterior and the posterior compartments of larval wings and of **(C)** eyespot and adjacent control tissues of pupal wings. **(D)** Volcano plot showing differentially expressed genes between the anterior and the posterior compartments of larval wings and of **(E)** eyespot and control tissues of pupal wings. Threshold parameters used for the Volcano Plot: Log2FoldChange = 1.3 and p-value = 0.01.

### Differential gene expression between eyespot and control tissue of pupal wings

To identify upregulated genes in eyespots we performed the same laser microdissection experiment in eyespots and adjacent control tissue (**Figure 1, Figure S2**) in 18-22 hrs pupal wings, followed by RNA-Seq and DEseq2 analysis. Character-tree analysis showed that biological replicates from each tissue clustered together (**Figure 1C**). Some of the significantly upregulated genes in the eyespots included Dpp signaling member *Mad6*; Hh signaling members *hedgehog, patched,* and *smoothened*; Wnt signaling members *wnt1, armadillo, pangolin,* and *dishevelled*, and other interesting genes such as *homothorax, dachshund, Ultrabithorax, Sp4-like, twisted gastrulation, grindelwald, Cyclin dependent kinase-14, aristaless-like, spaetzle 3*, and *kayak-like* (**Figure 1E, Table S2**). In the control tissue, highly expressed genes included serine proteases, Wnt repressor *groucho*, *obstructor E-like, peter pan, timeless homolog*, and *henna* (**Figure 1E, Table S2**). Interestingly *dpp, Mad3, Mad4*, and *cubitus interruptus* were not DE across tissues.

### *dpp* and *Mothers against dpp 6 (Mad6)* are expressed in eyespot centers and *dpp* functions in eyespot development

Since we could not identify the presence of *dpp* in our DEseq eyespot data (performed at 18-22 hrs) we conducted hybridization chain reaction (HCR3.0) (Choi et al., 2018), as well as enzyme-based *in-situ* hybridizations at a few earlier and later stages of pupal development. We had inconsistent spatial expression results in 16-24 hrs pupal wings with some showing clear expression of *dpp* in the eyespot centers (**Figure 2F and G, Figure S4 Q and R**), while others not (**Figure S4 A and B**). The 16-24 hrs window overlaps the window of development when protein products of the known Dpp target gene *spalt* are observed in the black scale cells (Monteiro et al., 2006; Brunetti et al., 2001). In wings where *dpp* expression was observed, it localized to only five of the seven eyespot centers, from M1-Cu1 (**Figure 2F and G**). This suggests that the period investigated for our RNA-Seq experiment might be close to the end of Dpp signaling in eyespot centers.

**Figure 2:**
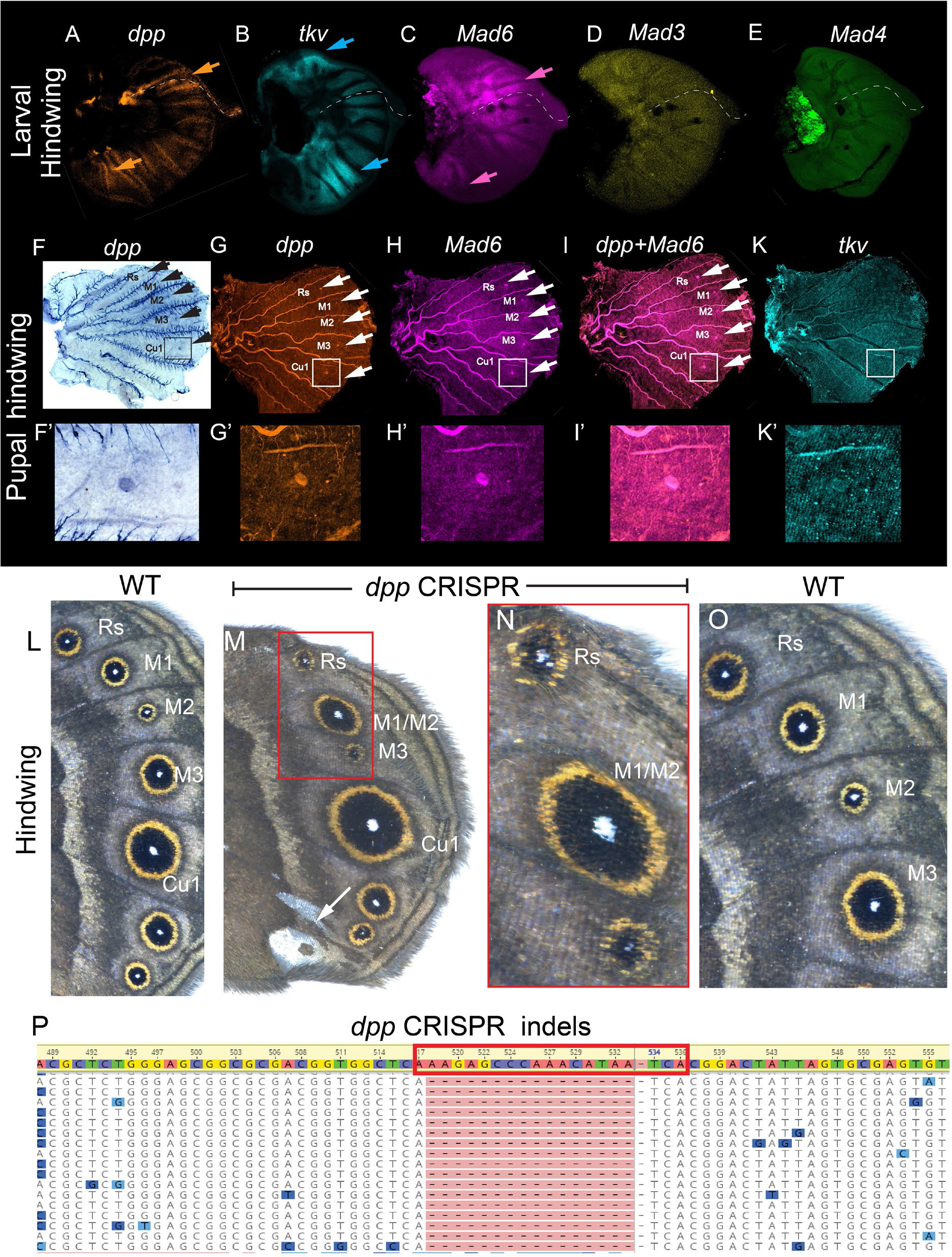
Expression of *decapentaplegic (dpp), thickvein (tkv), Mothers against dpp 6 (Mad6), Mad3,* and *Mad4* in larval and 16-24 hrs pupal hindwings of *B. anynana*. Expression of (**A**) *dpp* in the anterior compartment at the AP boundary (dotted line), (**B**) *tkv* in the anterior and middle posterior wing regions (blue arrows), and (**C**) *Mad6* in a broad domain spanning the AP boundary and in a small lower posterior domain (pink arrow) in the larval hindwings. No expression of (**D**) *Mad3* and (**E**) *Mad4* was observed in larval hindwings. (**F and G**) Expression of *dpp* in the eyespot center cells, and (**H**) of *Mad6* in the eyespot centers and in a broad domain spanning the AP boundary. (**I**) Co-expression of *dpp* and *Mad6*. (**K**) No specific domain of expression was observed for *tkv*. **(L)** Adult WT. **(M and N)** CRISPR-Cas9 knockout of *dpp* during embryonic development resulted in the loss of either the M1 or M2 eyespot, increases in size of two eyespots M1 (or M2) and Cu1, and ectopic appearance of silver scales (white arrow) in a single individual. (**O**) Adult WT hindwing. (**P**) Deletions (red shaded region) at 3 bps 5’ of the PAM sequence of the *dpp* guide RNA site (boxed area) obtained using DNA extracted from the wing shown in panels **M** and **N**.

To confirm the deployment of the Dpp signaling pathway in eyespots, we examined the spatial expression of other pathway members, including the receptor *tkv* and the transcription factors *Mad6*, *Mad3* and *Mad4*. During larval wing development, we observed two strong domains of *tkv* away from the strong domains of *dpp* expression (**Figure 2A and B, Figure S3J, N-P**), consistent with data from *Drosophila* where *tkv* is expressed away from the source of Dpp proteins (Zecca et al., 1996). *Mad6* was expressed strongly in a broad domain spanning the AP boundary where its likely transducing the Dpp signaling (**Figure 2C, Figure S3K**), and no expression domains for *Mad3* and *Mad4* were observed (**Figure 2D and E, Figure S3L and M**). In the pupal stages (at 16-24 hrs after pupation) *Mad6* was expressed in the center of the eyespots consistent with the RNA-seq data mentioned above, along with the broad domain spanning the AP boundary conserved from the larval stage (**Figure 2H, Figure S4R**). No specific domain of expression was observed for *tkv* during the pupal stage (**Figure 2K, Figure S4R**).

To test the function of *dpp* in eyespot development we used CRISPR-Cas9. Knocking out *dpp* by injecting a single guide RNA (sgRNA) and Cas9 during the embryonic stages proved to be extremely difficult and resulted in high embryonic mortality. This is likely due to *dpp* being involved in patterning the early embryos (Ashe et al., 2000). Late embryonic knockouts (around 6 hrs) of *dpp* resulted in one individual with the loss of an anterior hindwing eyespot, increases in the size of two other eyespots (one being the large Cu1 eyespot), loss of either M1 or M2 veins, incomplete development of M1/M2 and M3 veins (**Figure S5**), and the appearance of ectopic silver scales (**Figure 2M and N, Figure S5**). These phenotypes are partially like *Ultrabithorax (Ubx)* knockout phenotypes (Matsuoka and Monteiro, 2021), that essentially transform a hindwing into the morphology of a forewing, leading to the ectopic appearance of silver scales, and changes in eyespot number and size.

### *Optix* and *spalt* are expressed in larval wings and in eyespot rings of pupal wings

Next, we examined the expression of *Optix* and *spalt* mRNA using HCR. During larval wing development, *Optix* was expressed strongly in an anterior domain on the hindwing (**Figure 3A**), whereas it was expressed in an additional domain on the forewing, in a posterior region (**Figure S3A and E**). *spalt* was expressed in four distinct domains in both hindwings and forewings (**Figure 3B, Figure S3H**), consistent with a previous study (Banerjee and Monteiro, 2020). In 24 hrs pupal wings, *Optix* was expressed in the upper and proximal margin of the wing, in a region that will produce silver coupling scales (Prakash et al., 2022), in distinct bands along the proximal-distal axis, corresponding to a marginal band and two bands of the central symmetry system, and in scale cells of the eyespot’s orange ring (**Figure 3D, Figure S4C, D, K, and L**). *spalt* was expressed in the future black scale cells of the eyespots (**Figure 4E**). Both genes formed a sharp boundary corresponding to the boundary observed between black and orange scale cells in adult individuals (**Figure 3F**). The exact same expression patterns were observed for the protein product of both genes using antibodies (**Figure S6**).

**Figure 3:**
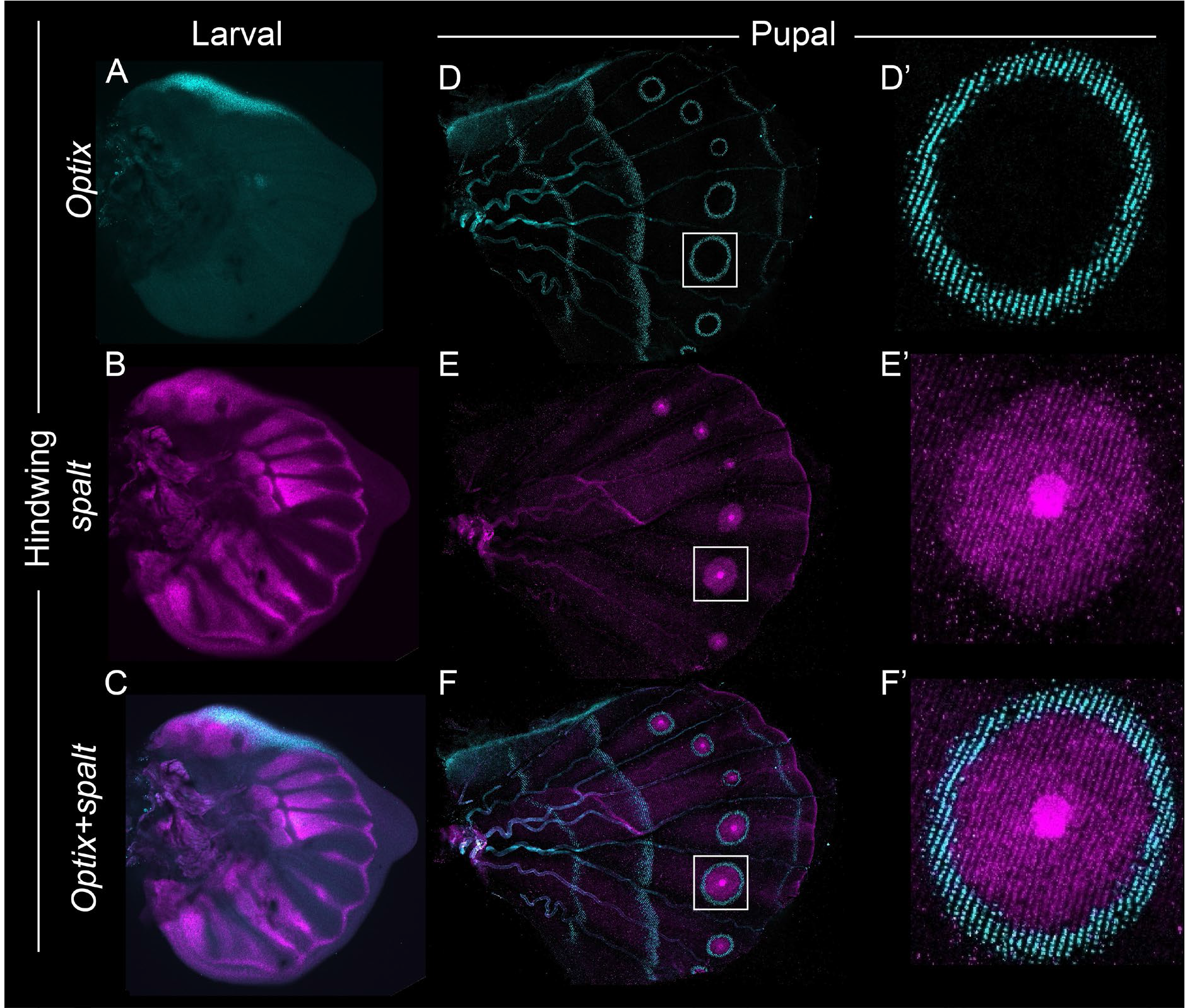
Expression of *Optix* and *spalt* in the larval and pupal wings of *B. anynana*. Expression of (**A**) *Optix* in the upper anterior compartment and (**B**) *spalt* in the four major domain of the larval wing disc. (**C**) Co-expression of *Optix* and *spalt*. Expression of (**D**) *Optix* in the eyespot orange ring cells, in the anterior and proximal margin of the wing, and in three bands transversing the proximal-distal axis, and (**E**) *spalt* in the future black scale cells of the eyespots, discal cell cross-vein, and along the wing margin. (**F**) Co-expression of *Optix* and *spalt*.

**Figure 4:**
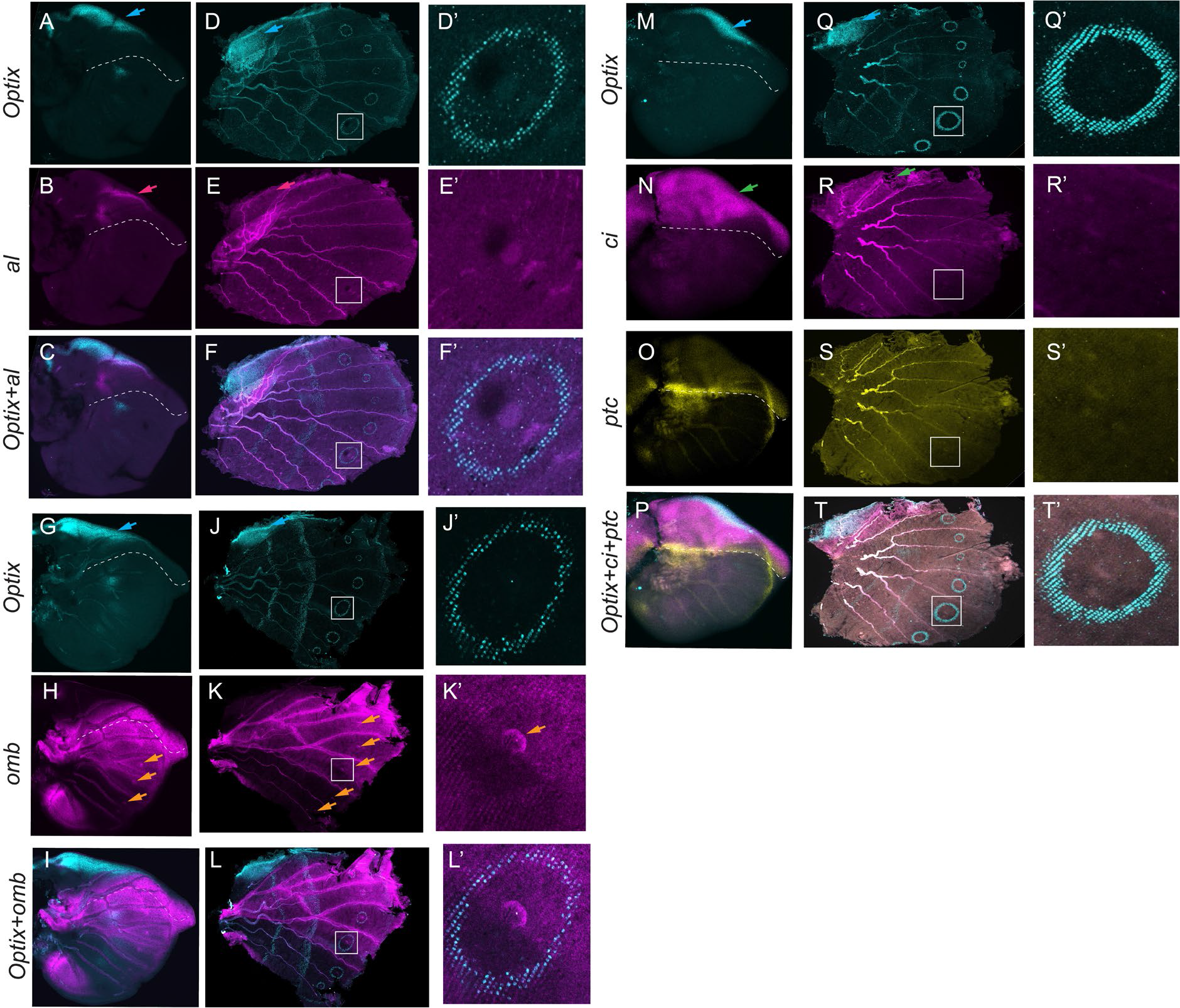
Expression of *Optix, aristaless (al), optomotor-blind (omb), cubitus interruptus (ci),* and *patched (ptc)* in the larval and 18-24 hrs pupal hindwings of *B. anynana*. *Optix* expression (**A, G, and M**) is used as a marker for the other genes. (**B**) *al* is expressed in the anterior compartment (pink arrow) of the larval wing. (**C**) Co-expression of *Optix* and *al*. (**D, J, and Q**) Expression of *Optix* in the eyespot ring and in the proximal-distal bands (blue arrow), and (**E**) *al* in the anterior compartment (pink arrow) in pupal hindwing. (**F**) Co-expression of *al* and *Optix*. (**H**) Expression of *omb* in a broad domain spanning AP boundary and in the lower posterior domain of the larval hindwing. Eyespot center specific expression was also observed during the larval stage (orange arrow). (**I**) Co-expression of *Optix* and *omb* in the larval hindwing. (**K**) In the pupal hindwing *omb* has the conserved AP expression domain along with faint expression in the orange ring and in the eyespot centers (orange arrows). (**N**) Expression of *ci* in the anterior compartment (green arrow) and of (**O**) *ptc* in the AP boundary and in the anterior compartment of larval hindwings. (**P**) Co-expression of *Optix, ci,* and *ptc* in the larval hindwing. (**R**) Expression of *ci* in the anterior compartment (green arrow) and of (**S**) *ptc* in the pupal hindwing. No specific domain of expression was observed for *ptc*. (**T**) Co-expression of *Optix, ci* and *ptc*. Dashed line in larval wings indicates the AP boundary.

### *optomotor-blind (omb)* is also expressed in larval wings and in eyespots

We next investigated the expression of other DE genes, also involved in the vein positional information mechanism in flies, including *al*, *omb*, *ci*, and *ptc* (Bier, 2000; Blair, 2007; De Celis, 2003). Both *al* and *omb* are known targets of Dpp signaling at the AP boundary, with *al* responding to Dpp at lower levels, further away from the boundary (Campbell et al., 1993; Martín et al., 2017). *al* showed upper anterior compartment expression in both larval and pupal wings (**Figure 4B and E**), consistent with a previous study in *B. anynana* (Banerjee and Monteiro, 2020), whereas *omb* was expressed in a large domain spanning the AP boundary in both larval and pupal wings (**Figure 4H and K, Figure S3F**). The small domain of *omb* expression observed in the lower posterior compartment in larval wings is likely activated by the lower posterior band of Dpp signaling that exists in butterfly wings (**Figure 2A, Figure S3J, Figure 4H, Figure S3F**). Faint eyespot center and orange ring expression was also observed for *omb* during the pupal stage (**Figure 4K**). Both *ci* and *ptc* showed anterior compartment specific expression in larval hindwings and forewings (**Figure 4N and O, Figure S3B and C**), *ptc* being strongly expressed along the AP boundary (**Figure 4O, Figure S3C**) where it likely transduces Hh signaling with the help of *ci* (Tanimoto et al., 2000). The anterior expression of *ci* was conserved in the pupal stage (**Figure 4R, Figure S4I and M**). No eyespot-specific expression was observed for *al, ci*, and *ptc* at either the larval or pupal stage (**Figure 4, Figure S4**). In summary, all these fly venation genes show conserved patterns of expression in early butterfly larval wings, indicating potential conservation of function, whereas only *omb* showed a novel expressed in eyespots.

To test if *omb*, *ci,* and *ptc* play a role in eyespot development, we again used CRISPR-Cas9. The single *ci* crispant individual had a defective anterior compartment but no eyespot defects (**Figure S9**), and *omb* and *ptc* didn’t showed any observable phenotype (**Table S9 and S10**). This suggests that these genes may play no role in eyespot development.

### *Optix* and *spalt* are required for the development of orange and black scales, respectively, and Spalt represses *Optix* expression

To test the function of *Optix* and *spalt* we injected CRISPR-Cas9 targeting each of the genes separately into early embryos. Individuals that emerged from these injections were mosaics where some cells were disrupted for *Optix* or *spalt* function and others were wildtype (**Figure 5A-F**). Some *Optix* crispants showed transformation of their orange-colored scales to a brown color (**Figure 5 A, B; Figure S7**) but did not show changes in the area of black scales, whereas *spalt* crispants developed orange scales in the black scale region of the eyespots (**Figure 5E and F; Figure S8**), as previously shown (Murugeshan et al., 2022). We confirmed that knocking out *Optix* and *spalt* led to deletion of nucleotides mostly near the targeted sites (**Figure 5S-U, Figure S7K and L, Figure S8K**). These results show that both *Optix* and *spalt* are required for the development of the orange and black scale cells, respectively. They also suggest that Spalt might be repressing the expression of *Optix* in the central black disc domain, because in *spalt* crispants, these central black scale cells become orange.

**Figure 5.**
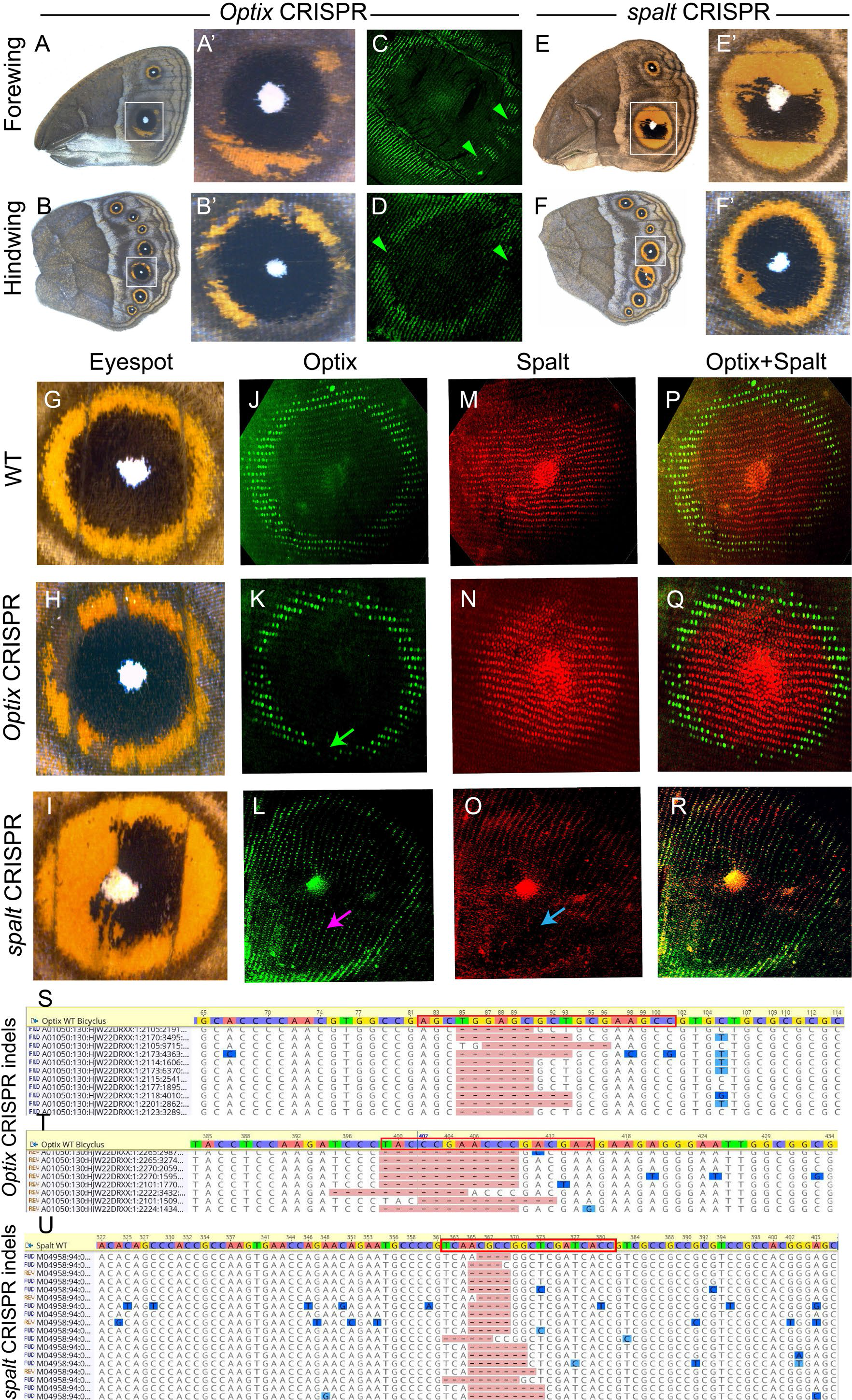
*Optix* and *spalt* are required for the development of orange and black scales, respectively, and Spalt prevents *Optix* from being expressed in the black central region of the eyespot. **(A and B)** CRISPR-Cas9 knockout of *Optix* produced defects in the development of the eyespot’s orange ring in both wings (S, T are amplified photos of the eyespots boxed in Q, R). **(C and D)** *Optix* CRISPR results in the loss of Optix protein in cells of the future orange ring area of the eyespots. **(E and F)** CRISPR-Cas9 knockout of *spalt* leads to orange scales developing inside the black scale domain. **(G)** WT eyespot, **(I)** *Optix* crispant, and **(J)** *spalt* crispant. **(J-L)** Eyespots stained with an antibody against Optix in WT, *Optix* crispant, and *spalt* crispant individuals (Note that different individuals are used for adult phenotyping and immunostains). **(L)** *spalt* crispant with Optix protein present in the eyespot central disc. **(M-O)** Eyespots stained with an antibody against Spalt in WT, *Optix* crispant and *spalt* crispant individuals. **(P-R)** Merged channels of Spalt and Optix. **(K)** Green arrow marks the region of missing Optix protein in the *Optix* crispant individual that does not affect Spalt-expressing cells. Pink arrow marks the ectopic presence of Optix, where Spalt proteins are missing (blue arrow). All wings were stained at 16-20 hrs pupal development. CRISPR deletions at the **(S and T)** *Optix* and **(U)** *spalt* target sites in three distinct individuals.

To test the regulatory interaction between *spalt* and *Optix* in eyespots, we performed knockouts of each gene in turn, followed by immunostainings targeting the protein products of both genes. *Optix* crispants showed loss of Optix proteins in cells of the orange ring **(Figure 5C, D, and K; Figure S7I),** but did not show changes in the presence of Spalt proteins in those cells (**Figure 5N and Q**). These data indicate that *Optix* is involved in giving the orange ring its color, but this gene is likely not involved in regulating the outer perimeter of the eyespot’s black disc nor *spalt* expression in the eyespots. *spalt* crispants showed loss of Spalt protein **(Figure 5O)**, and ectopic levels of Optix protein in the black scale disc region **(Figure 5 L; Figure S8D and E)**. These data indicate that *spalt* represses *Optix* expression in the central black disc region of the eyespot. In the eyespot centers (white pupils), Optix proteins are generally absent, but in some individuals, both Optix and Spalt proteins are found there (**Figure 5L, O, and R**), indicating that in these cells, the regulatory interaction between these genes is different.

## Discussion

Our study showed that several genes previously implicated in venation patterning in *Drosophila* wings are expressed in largely conserved, homologous domains in *B. anynana* butterfly wings, as well as in a novel trait, butterfly eyespots. In both species, at early stages of wing development, *dpp* is expressed along the AP boundary (**Figure 1A, Figure S3J and O**), and the downstream targets *spalt, omb, al,* and *Optix* are expressed in partially overlapping bands, spanning the central Dpp stripe, or some distance away from it (Banerjee and Monteiro, 2020; Blair, 2007; Martín et al., 2017) (**Figure 3C, Figure 4A-C and G-I**). The activation of all these genes is likely carried out by Mad6, a Dpp signal transducer (Matsuda and Affolter, 2017) expressed in a broad domain spanning the AP boundary (**Figure 1C, Figure S3K**). At later stages of development, in the pupal wing, a subset of these genes is also expressed in and around the rings of butterfly eyespots, suggestive of network co-option. In particular, *dpp* is found in the center of the eyespots, and *spalt*, *omb*, and *Optix*, are found in concentric circles of expression at variable distances from the center.

In addition to these suggestive expression patterns, we showed that the genetic circuit involving the repression of *Optix* by Spalt proteins, that sets-up vein L2 in *Drosophila* (Martín et al., 2017), is also conserved in *B. anynana* to set up the sharp boundary between the black and orange color rings in eyespots. In both species, the expression of *Optix* abuts the expression of *spalt* in the anterior compartment of larval wings (**Figure 3A-C**). In *Drosophila* wing discs, down-regulation of *spalt* with RNAi results in the expansion of the Optix domain in the anterior compartment (Martín et al., 2017). However, downregulation of *Optix* with RNAi does not affect the spatial domain of Spalt (Martín et al., 2017). Similarly, in the eyespots, Spalt also represses *Optix’s* expression from its own domain (**Figure 5G-R**). We showed that *Optix* is required for orange and *spalt* for black scale differentiation, and disruptions to *Optix* lead to the loss of Optix proteins and orange scales but do not affect the expression domain of Spalt (**Figure 5H, K, N, and O**), whereas disruptions of *spalt* result in the ectopic expression of Optix, and ectopic appearance of orange scales (**Figure 5I, L, O, and R**).

Venation patterning in *Drosophila* also involves other key members from the Dpp and Hh signaling pathways (Blair, 2007) that were DE across anterior and posterior compartments in *B. anynana* larval wings, and further visualized with HCR (**Figure S1, S3, and S4**). Hh, expressed in the posterior compartment in both species (Blair, 2007; Saenko et al., 2011; Tabata et al., 1992) (**Figure S1**), is known to activate the expression of *dpp* at the AP boundary in fly wings by binding to the receptor Patched (Ptc), and transducing the signal via Cubitus interruptus (Ci) (Banerjee and Monteiro, 2020; Bier, 2000; Tanimoto et al., 2000). Correspondingly, both *ptc* and *ci* were expressed strongly in the anterior compartment of *B. anynana* larval wings (**Figure 4N-P**) (Keys et al., 1999; Saenko et al., 2011). Perhaps due to the timing of the stainings, we were unable to localize the transcripts of *ci* and *ptc* in the eyespots during larval and pupal stages with HCR (**Figure 4N, R, O, S; Figure S4I, J, M, and N**). Previous researchers, however, did visualize Ci protein in *B. anynana* eyespot centers (Keys et al. 1999) and *ptc* transcripts in *Junonia coenia* eyespot centers in larval wings. While we were also unable to visualize the Dpp target gene *al* in larval and pupal eyespots (**Figure 4B and E**), *omb* showed faint expression in the center of the eyespot and in the orange ring cells (**Figure 4H and K**). Knocking-out *ci, ptc,* and *omb* using CRISPR-Cas9 didn’t result in any eyespot-specific phenotype (**Figure S9, Table S9, Table S10**). These results suggest that *al, omb*, and Hh signaling members *ci* and *ptc* may not play a role in eyespot development. Thus, the gene-regulatory network (GRN) patterning the eyespot rings of *B. anynana* likely involves only a subset of the venation GRN with *spalt* and *Optix* being important regulators of the sharp and prominent orange and black rings of color (**Figure 6**).

**Figure 6.**
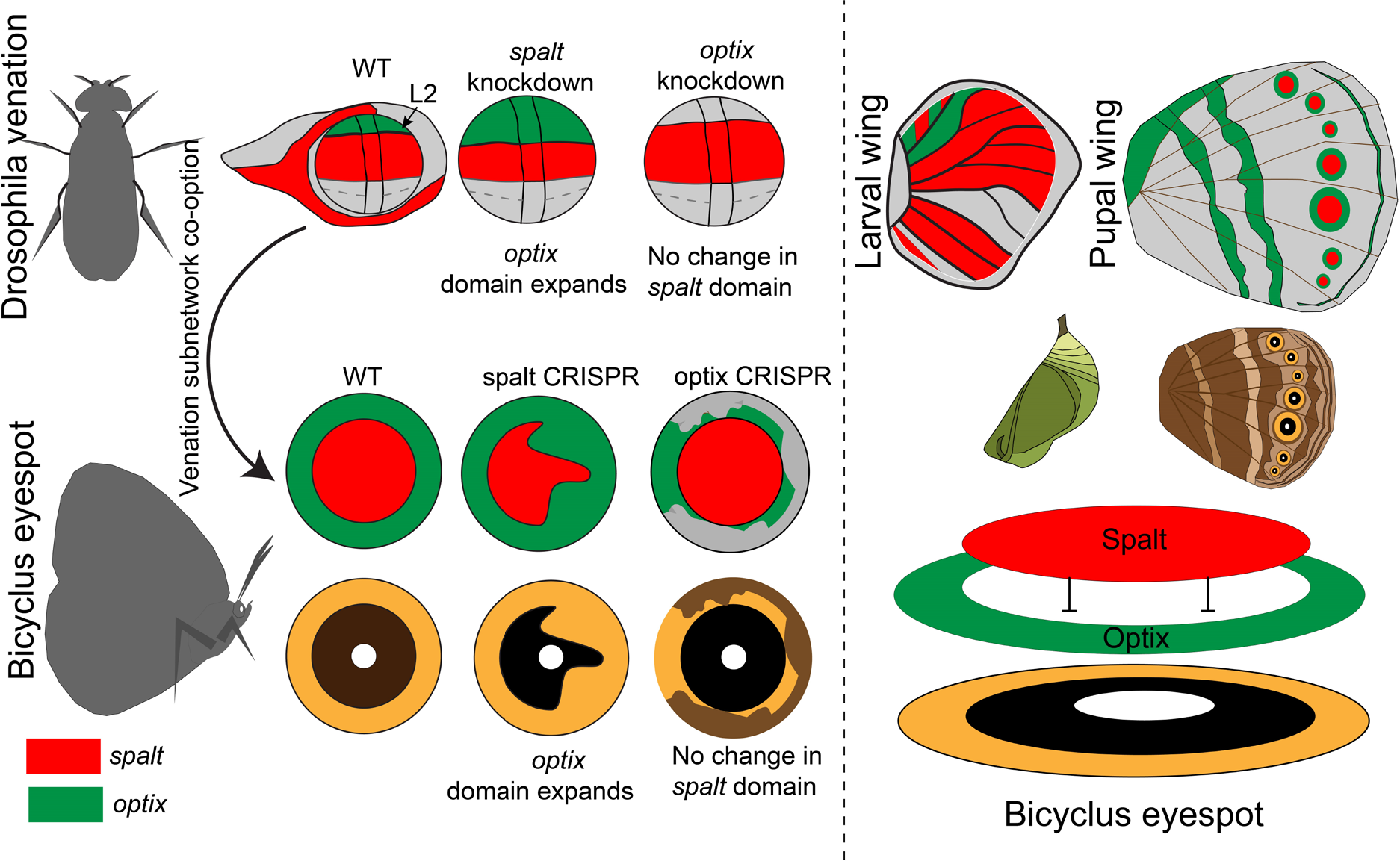
Co-option of a venation subnetwork involving *spalt* and *Optix* in patterning the eyespot rings of *B. anynana*. In *Drosophila*, larval wing venation patterning involves the negative regulation of *Optix* by Spalt in the anterior compartment, at the location of vein L2, where Optix does not cross-regulate the *spalt* expression domain. The same exact regulation is observed in *B. anynana* eyespots where the loss of Spalt proteins results in the ectopic expression of Optix proteins and orange scales, while the loss of Optix only affects the orange scales without affecting the expression of Spalt proteins or the development of black scales. Spalt thus, represses *Optix* in the black scale region and creates a sharp color boundary in the eyespot rings.

The phenotypes we observed with embryonic *dpp* knockouts suggest that *dpp* has an early role is establishing hindwing identity, in addition to a possible later role in eyespot ring formation. Knockouts of *dpp* during the embryonic stages proved quite challenging due to *dpp’s* role in embryonic development (Ashe et al., 2000)). The single crispant individual we obtained had fewer hindwing eyespots but also had other eyespots enlarged, making that hindwing resemble a forewing. The loss of the M1 (or M2) eyespot was likely due to the complete loss of Dpp along one vein (either the M1or M2 vein) (**Figure S5**) (Banerjee and Monteiro, 2020). However, the hindwing acquiring forewing features, such as enlarged M1/M2 and Cu1 eyespots, and the development of silver scales in the posterior compartment (**Figure 2M and N, and Figure S5**), are not consistent with a mere eyespot developmental function for *dpp*. We propose that these features could be due to an interaction between *Ubx* and *dpp,* as a previous *Ubx* crispant in *B. anynana* showed a similar phenotype (Matsuoka and Monteiro, 2021). In *Drosophila, Ubx*, a Hox gene responsible for hindwing identity, controls both the transcription and mobility of Dpp in the haltere (Crickmore and Mann, 2006). We propose that Ubx is playing a similar role in *Bicyclus* hindwings during the larval stage. At this stage, *dpp* expression surrounds the expression of Armadillo (a Wnt transducer) and Distal-less proteins in the eyespot centers (Connahs et al., 2019). The increase in size of M1 and Cu1 eyespots with *dpp* knockouts (**Figure 2M and N**) could be due to the proposed role of Dpp proteins in restricting the expression of these two other genes to the eyespot centers via a reaction-diffusion mechanism (Connahs et al., 2019). If Dpp proteins are removed, the centers might be able to expand, leading to larger eyespots. Also, if Ubx is regulating both *dpp* expression and mobility, *dpp* knockouts might mimic *Ubx* knockouts. Disentangling the early and late roles of *dpp* in eyespot development will only be possible once we are able to knock-out *dpp* in late wing development with somatic CRISPR.

Two roles were also observed for *spalt:* in eyespot center differentiation during the larval stage, and in the differentiation of the black disc during the pupal stage. *spalt* is likely playing a role in a proposed reaction-diffusion mechanisms proposed for focus differentiation during the larval stages (Connahs et al., 2019), because *spalt* knockouts result in loss of eyespots (Murugeshan et al., 2022) or in split eyespot centers (**Figure S8J**). During the pupal stage, however, disruptions of *spalt* lead to transformation of black to orange scales, via the *spalt-Optix* regulatory mechanism identified here.

Several knowledge gaps still remain surrounding eyespot ring differentiation. Support for the vein network GRN co-option proposed in this study would benefit from testing whether *Optix* and *spalt* are responding to Dpp signals produced at the center of the eyespots transduced through the transcription factor Mad6. It is also unclear whether Dpp and Wingless (Wnt1) are jointly required for eyespot ring differentiation. Wnt1 is another proposed morphogen for eyespot development (Monteiro et al., 2006; Özsu et al., 2017) and its function should be newly investigated using knockouts. In the present work we found evidence for the presence of many Wnt signaling members such as *Wnt1, armadillo, pangolin,* and *dishevelled* highly upregulated in the pupal eyespot tissue (**Figure 1E**). In addition, the role of *engrailed* (*en*) in eyespot ring differentiation, a gene co-expressed in the same orange cells as *Optix*, is still unclear (Brunetti et al., 2001) (**Figure S10**). Multiple paralogs are expressed in eyespots (Banerjee et al., 2020), and disruptions of all paralogs simultaneously may need to be performed to overcome functional redundancy.

## Methods

### Rearing Bicyclus anynana

*Bicyclus* butterflies were raised in the lab at 27°C, 60% humidity and 12-12 hrs day-night cycle. Larvae were fed young corn leaves and adults were fed mashed bananas.

### CRISPR-Cas9

*Optix*, *spalt, ci, knirps, ptc, dpp,* and *omb* CRISPR-Cas9 experiments were carried out based on a protocol previously described (Banerjee and Monteiro, 2018). Guides were designed to target the coding sequence of the genes (see the supplementary file for sequences and the regions targeted). A solution containing Cas9 protein (IDT, Cat No. 1081058) and guide RNA, each at a concentration of 300ng/ul, diluted in Cas9 buffer, and a small amount of food dye was injected into 1509 embryos for *Optix*, 2474 embryos for *spalt,* 2012 embryos for dpp, 283 embryos for *ci,* 306 embryos for *ptc*, 1259 embryos for *knirps,* and 734 embryos for *omb* (**Table S4-S10**). A few of the *spalt* and *optix* injected individuals were dissected at 24 hrs after pupation to perform immunostainings, while the rest were allowed to grow till adulthood. After eclosing, the adults were frozen at −20° C and imaged under a Leica DMS1000 microscope.

Dpp synthetic guide based CRISPR-Cas9 experiment were carried out at embryonic developmental stages. The experiment was conducted in the same way as mentioned above except a synthetic guide from Integrated DNA Technologies (IDT) was used. Briefly a 20 bp region was selected using the IDT custom guide tool (AAAGAGCCCAAACATAATCA). For the preparation of the synthetic guide-Cas9 mixture, the *dpp* guide (crRNA) was mixed with equimolar amount of tracrRNA (IDT), heated at 95°C for 5 mins and cooled to room temperature. Afterward Cas9 enzyme was added and left for 20 mins at room temperature. Non-toxic food dye was added to the mix and injected into the embryos. After eclosion, the adults were frozen and imaged under a Leica DMS1000 microscope.

For identification of the indels, DNA from the mosaic CRISPants wings were isolated using Omega Tissue DNA extraction kit (Cat No: D3396-01). Afterwards PCR was performed to amplify the region of interest. PCR products were purified and sent to the next generation sequencing facility at Genewiz. After sequencing, the reads were aligned using Geneious 10.0 to the reference WT gene sequences.

### Immunostainings

Pupation time was recorded using an Olympus tough tg-6 camera, and pupal wings were dissected at different timepoints after pupation under a Zeiss Stemi 305 microscope in 1x PBS at room temperature based on a protocol previously described (Banerjee and Monteiro, 2020b). Wings were fixed using 4% formaldehyde in fix buffer (see **Table S11** for details), followed by four washes in 1x PBS, five mins each. After the washes, the wings were incubated in block buffer (see **Table S11** for details) at 4°C overnight. The next day primary antibodies against Optix (1:3000, rat, a gift from Robert D. Reed) and Spalt (1:20000, guinea pig GP66.1), were added in wash buffer and incubated at 4°C for 24 hrs. The next day anti-rat AF488 (Invitrogen, #A-11006) and anti-guinea pig AF555 (Invitrogen, # A-21435) secondary antibodies at the concentration of 1:500 in wash buffer were added followed by four washes in wash buffer, 20 mins each. Wings were then mounted on an inhouse mounting media (see **Table S11** for details) and imaged under an Olympus fv3000 confocal microscope.

### Enzyme based *In-situ* hybridization

For the visualization of *dpp* expression in pupal wings of *Bicyclus*, pupation time was recorded, and the wings were dissected from 18-24 hrs old pupae in 1x PBS under a Zeiss stemi 305 microscope based on a previously described protocol (Banerjee and Monteiro, 2020b) and transferred to 1xPBST with 4% formaldehyde. After fixation for 30 min the wings were washed three times in 1x PBST for 5 mins each. The wings were then treated with proteinase K and glycine and washed again three times using 1x PBST. After, the wings were gradually transferred into pre-hybridization buffer (see **Table S12** for composition) and heated at 65°C for 1 hr. Hybridization buffer with a *dpp* probe (see **Table S12** for composition) was added to the wings and they were incubated for 16 hrs at 65°C. Wings were then washed five times for 30 mins each in pre-hybridization buffer. After washing, wings were moved to room temperature and gradually transferred to 1x PBST and washed in 1x PBST. Afterwards, the wings were incubated in block buffer (see **Table S12** for composition) for 1 hr, followed by addition of anti-digoxygenin (Sigma-Aldrich, Cat No. 11093274910) diluted 1/3000 times in block buffer. After 1 hr of incubation the wings were washed five time, five mins each in block buffer. Finally, wings were transferred to alkaline-phosphatase buffer (see supp Table S4 for composition) supplemented with NBT-BCIP (Promega, Cat No. S3771) and incubated in the dark till the development of color. The wings were imaged under a Leica DMS1000 microscope.

### Hybridization chain reaction (HCR3.0)

HCR3.0 was carried out based on the protocol described in Choi et al., 2018 with the modifications for butterfly wing tissue described below. Larval and pupal wings were dissected in 1x PBS followed by fixation in 4% formaldehyde in 1x PBST. Afterwards wings were washed in 1x PBST permeabilized in a detergent solution (Bruce et al., 2021) and washed with 5x SSCT. Wings were incubated for 1 hrs in 30% probe hybridization buffer followed by overnight incubation in 30% probe hybridization buffer supplemented with HCR probes (see supp section for sequences) at 37°C. After incubation wings were washed five times with 30% probe wash buffer at 37°C. Afterwards wings were brought back to room temperature and washed twice with 5x SSCT followed by incubation in amplification buffer and overnight incubation in amplification buffer supplemented with fluorescent hairpin probes (Molecular instruments) in dark for 16-20 hrs. The next day wings were washed four times with 5x SSCT mounted in an inhouse mounting media and imaged under an Olympus fv3000 microscope.

### Laser micro-dissection

Laser microdissections of the anterior and posterior compartments of larval wings and eyespot and adjacent control tissue from pupal wings were carried out based on the protocol described in (Banerjee et al., 2022). Briefly, wings were dissected in 1x PBS and mounted in PEN membrane slides (Cat No.: LCM0522). Wings were washed with cold 50%, 75% and 100% ethanol followed by staining using Histogene staining solution (Cat. No.: KIT0415) for 20-30 seconds. Wings were washed twice with cold 100% ethanol and cold acetone. The slides were either kept at −80°C or immediately dissected using Zeiss PALM microbeam-Laser microdissection microscope using the parameters described in (Banerjee et al., 2022). 12 hindwings in the larval stage and 4 hindwings in the pupal stage were used for each biological replicate, respectively. A total of three biological replicates were used for downstream analysis.

### RNA-isolation from the laser dissected samples

Microsections of the larval and pupal wings were picked up from the membrane slide using a pair of fine forceps (Dumont, Cat. No. 11254-20) and transferred to 200 µl tubes with 50 µl distilled water. Afterwards, 100 µl of LCM lysis buffer (Banerjee et al., 2022) was added and the tubes were incubated at 55°C in a water bath for 6 mins with vortexing in between. The samples were then homogenized with 0.1mm zirconium beads (BioSpec, Cat. No. 11079101z) in a homogenizer (Next Advance Bullet Blender) for 3 mins followed by RNA isolation using a Qiagen RNA isolation kit (Cat. No. 74136). RNA samples were lyophilized in Genewiz RNA stabilization tubes and sent for sequencing (Next Generation Sequencing, Azenta US, Inc).

### RNA-seq analysis

Raw reads were analyzed using fastqc and cleaned using bbduk (Bushnell, 2014)with the parameters ktrim= r, k = 23, mink = 11, hdist = 1, tpe, tbo. Cleaned reads were aligned using *B. anynana* ncbi reference genome assembly (txid: 110368) using Hisat2 (Kim et al., 2019) and mapped to reference *B. anynana* gtf file obtained from ncbi (txid: 110368) using stringtie (Pertea et al., 2015). A gene count matrix table was created from the stringtie files using pydr.py (Pertea et al., 2015) package using python2.7 followed by analysis using DESeq2 in R (Love et al., 2014). Volcano plots were created using EnhancedVolcano package in R.

## Data availability

The sequencing data generated in the paper has been submitted to NCBI under the bio-project PRJNA948390.

## Authors contribution

TDB and AM designed the experiments, analysed the results, and wrote the manuscript. TDB performed the experiments. Both the authors read and agreed to the final version of the manuscript.

## Acknowledgements

We would like to thank Robert Reed for the anti-Optix antibody. Suriya Narayanan Murugesan and Shen Tian for helpful discussion on the manuscript. We would like to thank DBS-CBIS confocal facility and Ms. Tong Yan for access to the Olympus fv3000 confocal microscope. The authors declare no conflict of interest.

## Supplementary materials

**Figure S1:**
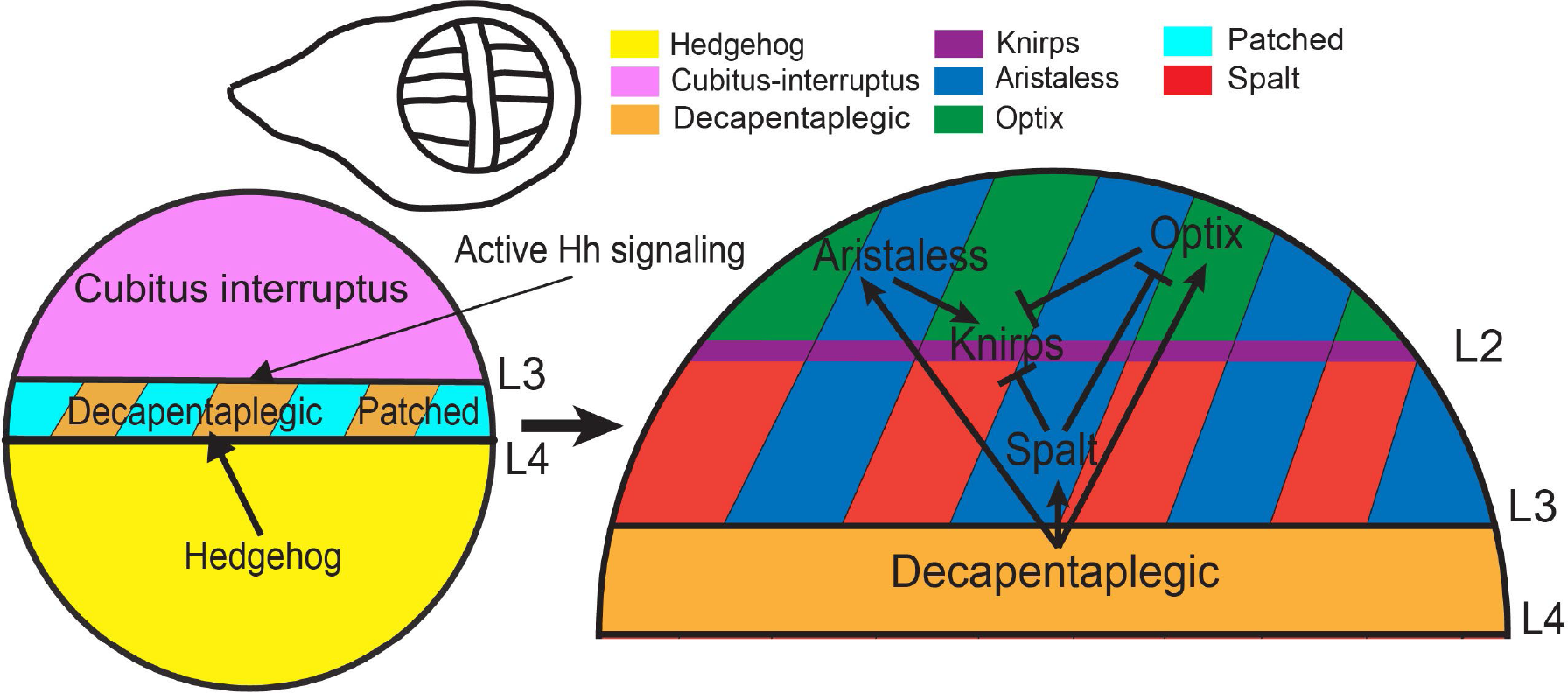
Venation patterning in the anterior compartment of *Drosophila melanogaster* (redrawn from Banerjee and Monteiro, 2020). Hedgehog (Hh), a short-range morphogen expressed in the posterior compartment of the larval wing disc in the presence of its receptor Patched and Cubitus interruptus activates Dpp signaling in a thin stripe of tissue in the anterior compartment. Dpp is capable of diffusing to the most anterior part of the larval wing disc where it activates key genes such as *aristaless* and *Optix* at low, and *spalt* at high concentration thresholds. Spalt protein represses the expression of *Optix* in the posterior part of the anterior compartment. Aristaless is responsible for the activation of another transcription factor *knirps* while both Spalt and Optix repress the activation of *knirps*. As a result, *knirps* is only activated in a thin stripe which forms the L2 vein of the *Drosophila* wing.

**Figure S2:**
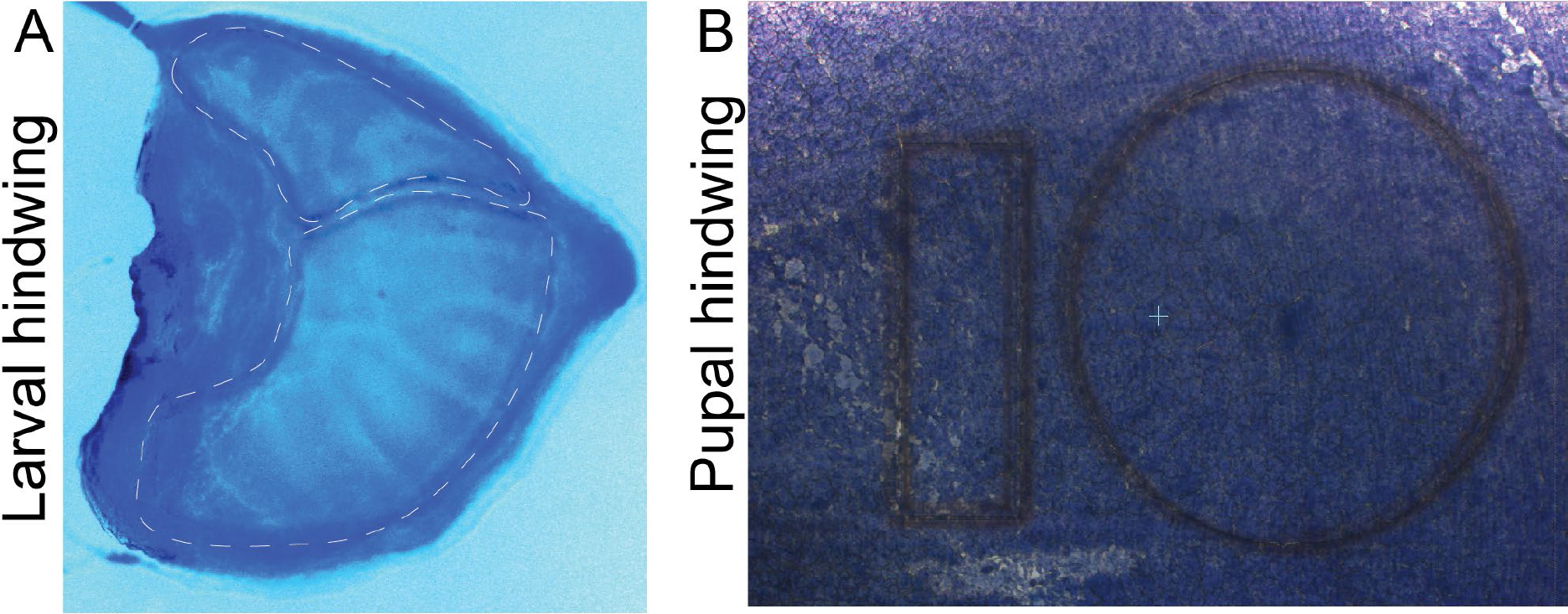
Areas dissected using laser microdissection in (A) Larval wing and (B) pupal wing of *B. anynana*.

**Figure S3:**
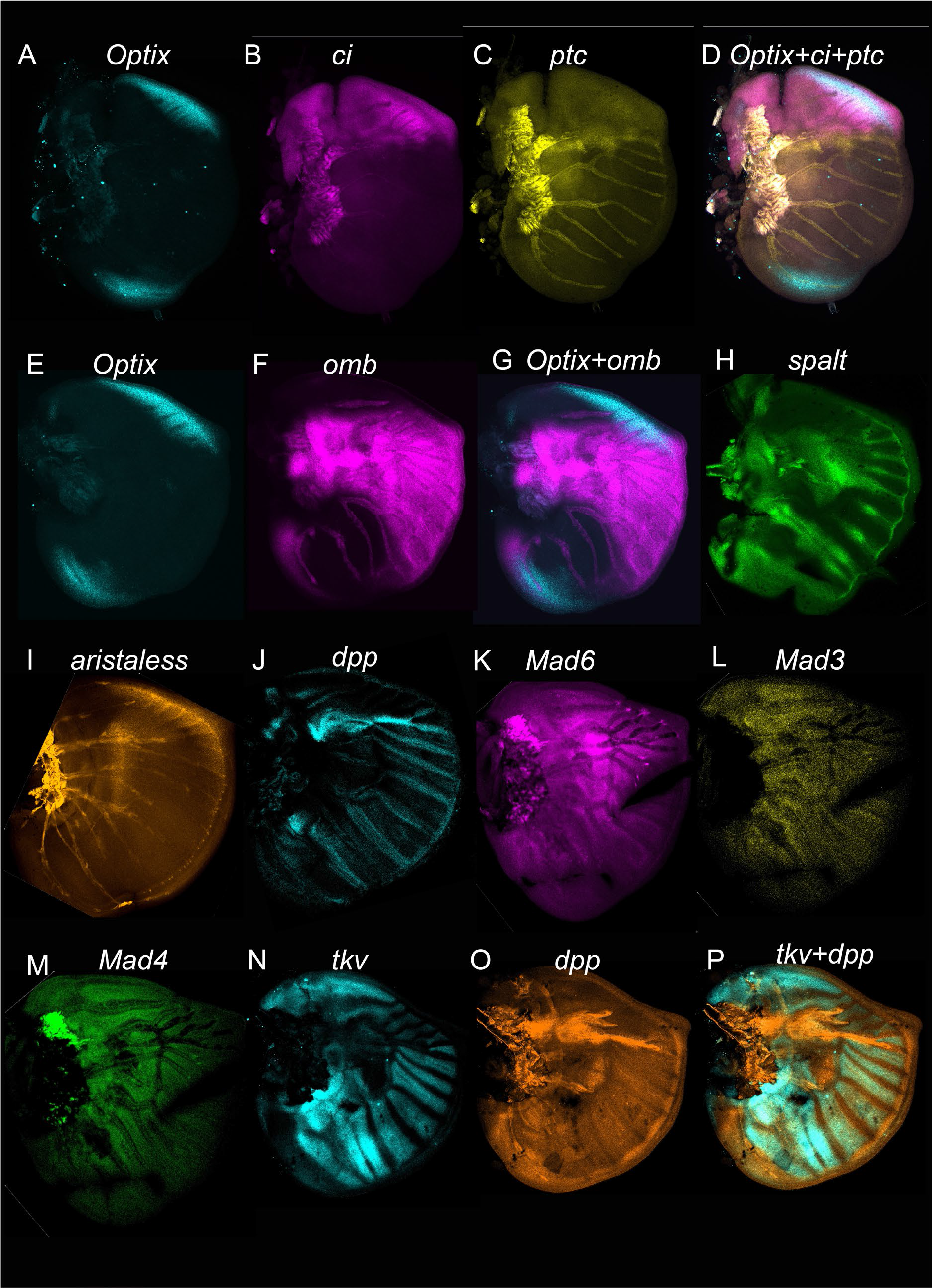
Expression of *Optix, ci, ptc, omb, spalt, aristaless, dpp, tkv, Mad6, Mad3,* and *Mad4* in the larval forewing of *B. anynana*. Expression of (**A**) *Optix* in the upper anterior and the lower posterior compartment, (**B**) *ci* in the anterior compartment, (**C**) *ptc* in the AP stripe and the anterior compartment, (**D**) Co-expression of *Optix*, *ci,* and *ptc*. Expression of (**E**) *Optix* in upper anterior and lower posterior domain and (**F**) *omb* in a broad domain spanning the AP boundary and the lower posterior compartment. (**G**) Co-expression of *Optix* and *omb*. Expression of (**H**) *spalt* in four main domains, (**I**) *al* in the anterior compartment, (**J**) *dpp* in the AP boundary stripe and in the lower posterior compartment, (**K**) *thickveins* in the upper anterior and in a domain spanning two sectors in the posterior compartment, (**L**) *Mad6* in the domain spanning the AP boundary. No expression of (**M**) *Mad3* and (**N**) *Mad4* was observed.

**Figure S4.**
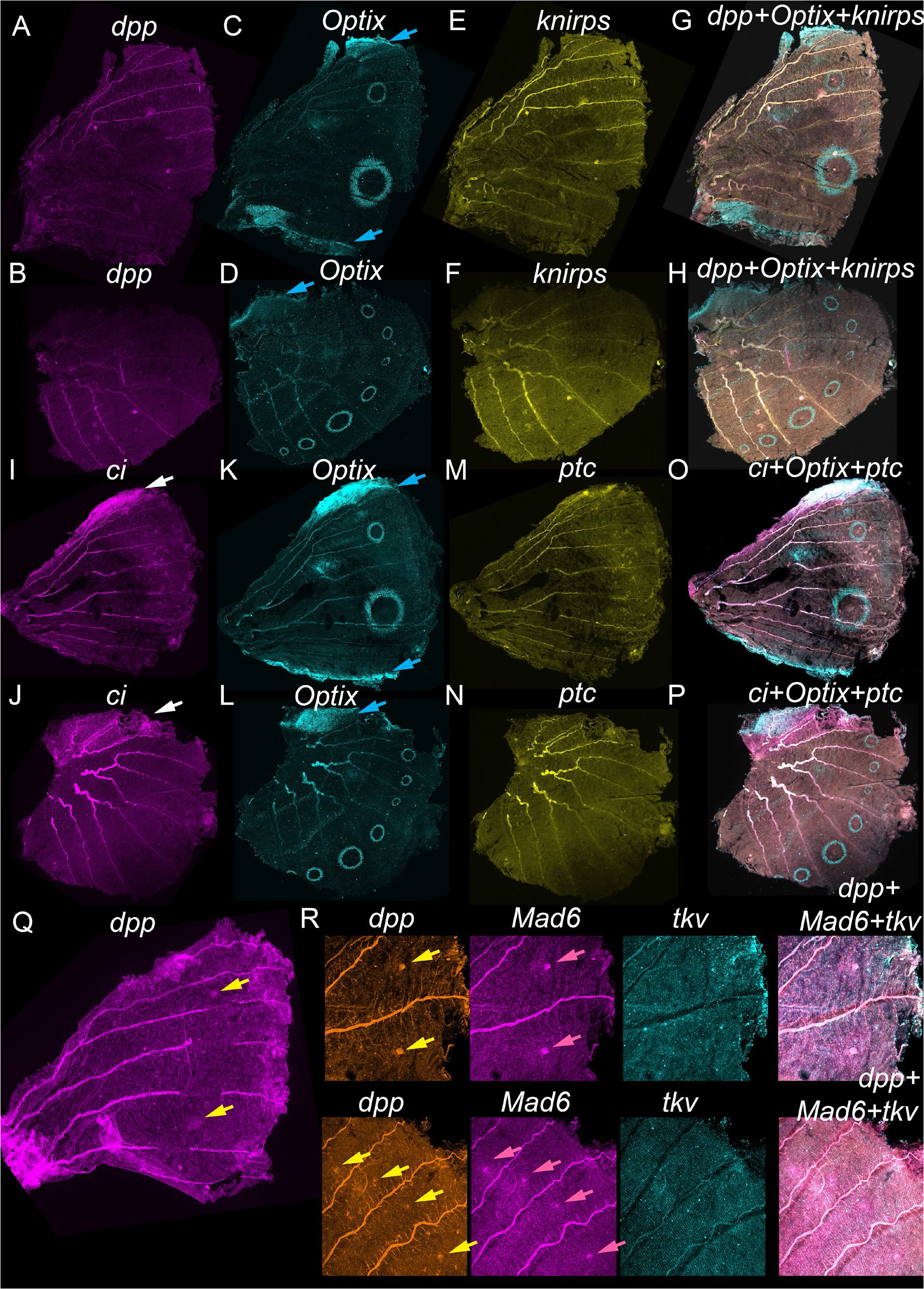
Expression of *dpp, knirps, Optix, ci, ptc*, *Mad6*, and *tkv* in 16-24 hrs pupal forewings and hindwings of *B. anynana*. Expression of *dpp* in **(A)** forewings and (**B**) hindwings showing no eyespot specific expression. Expression of *Optix* in (**C and K**) forewings showing expression in the upper anterior and posterior compartment and in the eyespot orange ring, and in (**D and L**) hindwings, showing upper anterior compartment specific expression, along three bands crossing the proximal-distal axis of the wing, and in the eyespot orange ring. No specific expression domain of *knirps* was observed in the (**E**) forewing, and (**F**) hindwing. Co-expression of *dpp, Optix*, and *knirps* in the (**G**) forewing and (**H**) hindwing. Expression of *ci* in the (**I**) forewing and (**J**) hindwing showing expression in the upper anterior compartment. No specific expression domain was observed for *ptc* (**M**) in forewings and (**N**) hindwings. Co-expression of *ci, Optix*, and *ptc* in the (**O**) forewing and (**P**) hindwing. (**Q**) Expression of *dpp* in a pupal forewing showing eyespot specific expression (yellow arrow). (**R**) Expression of *dpp, Mad6*, and *tkv* in pupal hindwing. *dpp* and *Mad6* shows expression in the eyespot center. While *tkv* doesn’t show any specific expression domain.

**Figure S5:**
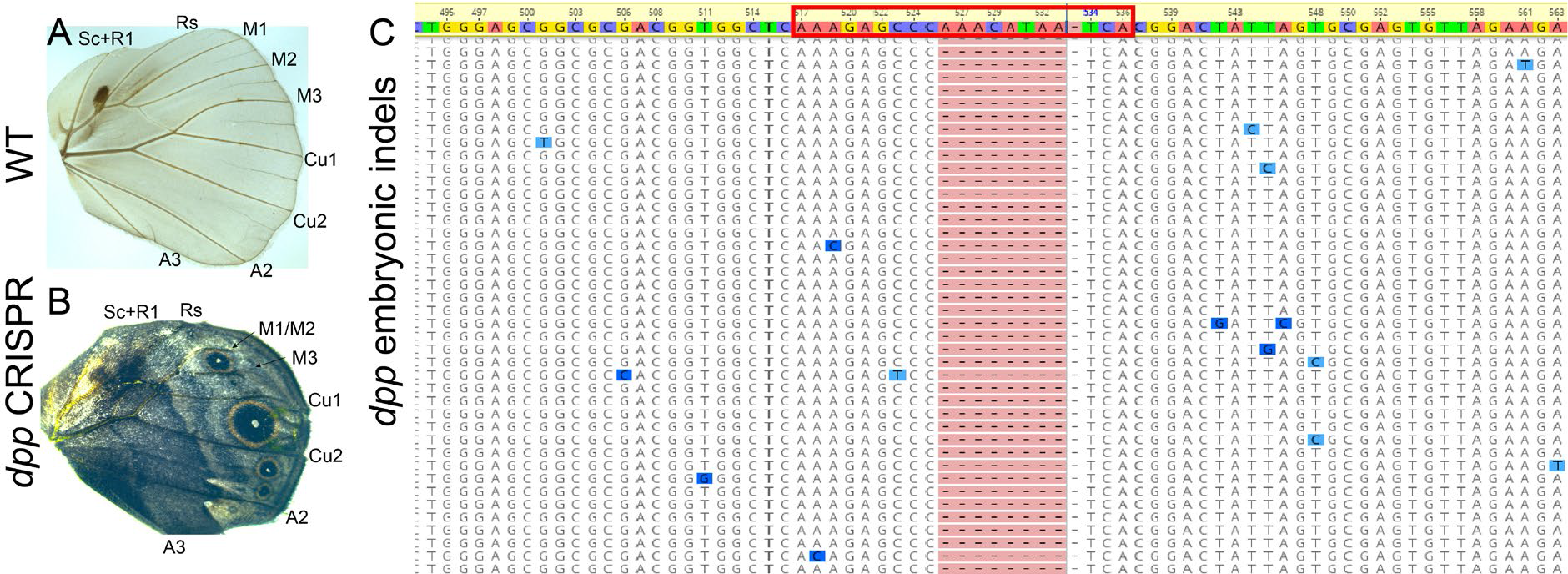
Effect of *dpp* CRISPR on venation and eyespot development. (**A**) WT adult hindwing venation. (**B**) Effect of disruptions to *dpp* on hindwing venation. We observed loss of M1 or M2 veins and incomplete formation of the M1/M2 and M3 veins. Loss of an eyespot, enlargement of eyespots, and ectopic silver scale development was observed in the hindwing of the *dpp* crispant. (**C**) Indels at the site of *dpp* CRISPR from the wing in panel **B**.

**Figure S6.**
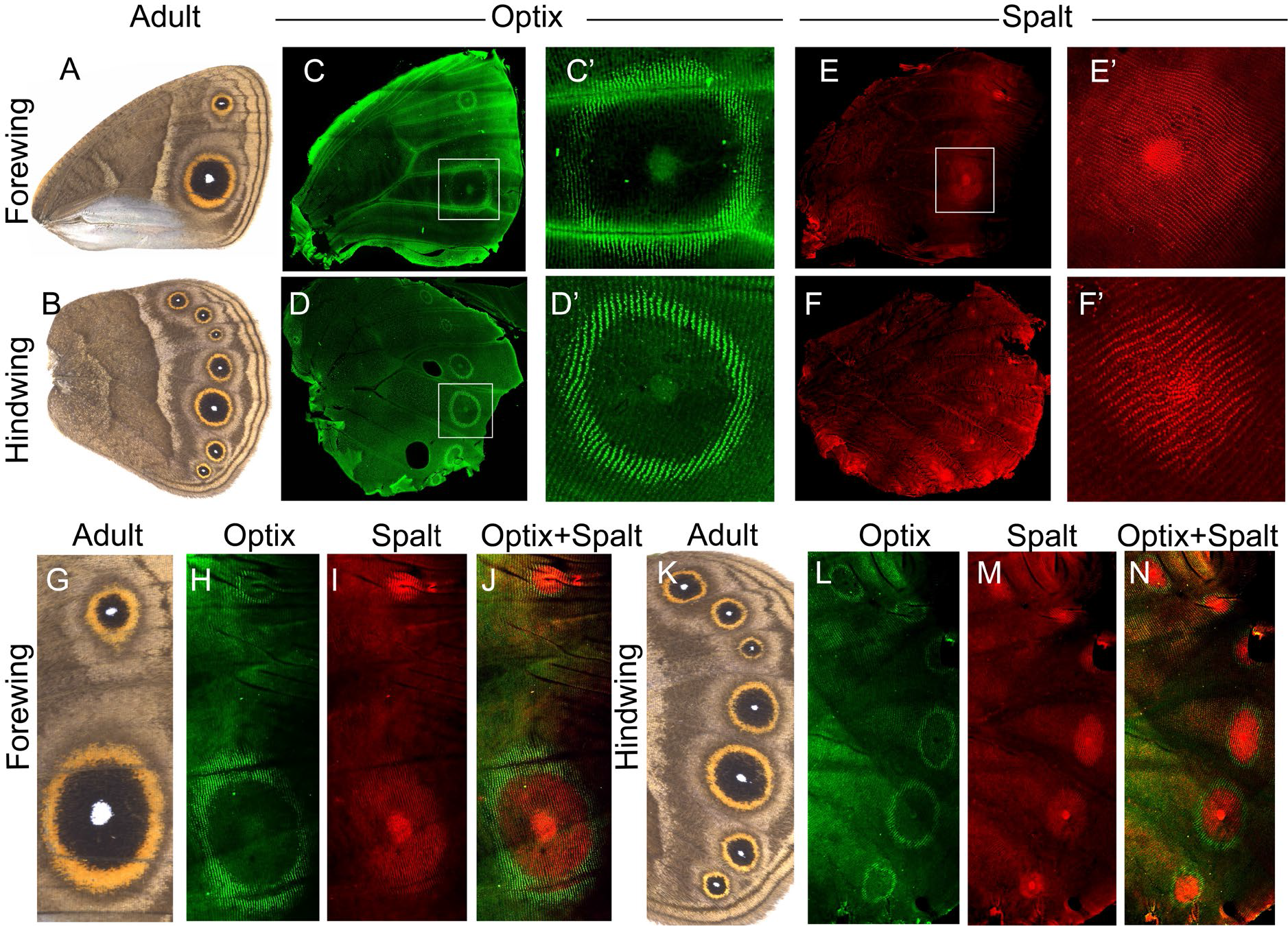
*B. anynana* adult wings and the localization of Optix and Spalt proteins in 16-44 hrs old pupal wings. **(A)** WT forewing, **(B)** WT hindwing. **(C)** The presence of Optix proteins in the forewing and **(D)** in the hindwing. **(E and F)** Presence of Optix in the orange ring of Cu1 eyespots (boxed in C and D). (**G**) Presence of Spalt protein in the forewing and **(H)** in the hindwing. **(I and J)** Presence of Spalt protein in the black disc of Cu1 eyespots. (**G**) Adult forewing. (**H**) Optix localization in the forewing. (**I**) Spalt localization in the forewing. (**J**) Merged channels of Optix and Spalt. (**K**) Adult hindwing. (**L**) Optix localization in the hindwing. (**M**) Spalt localization in the hindwing. (**N**) Merged channels of Optix and Spalt.

**Figure S7.**
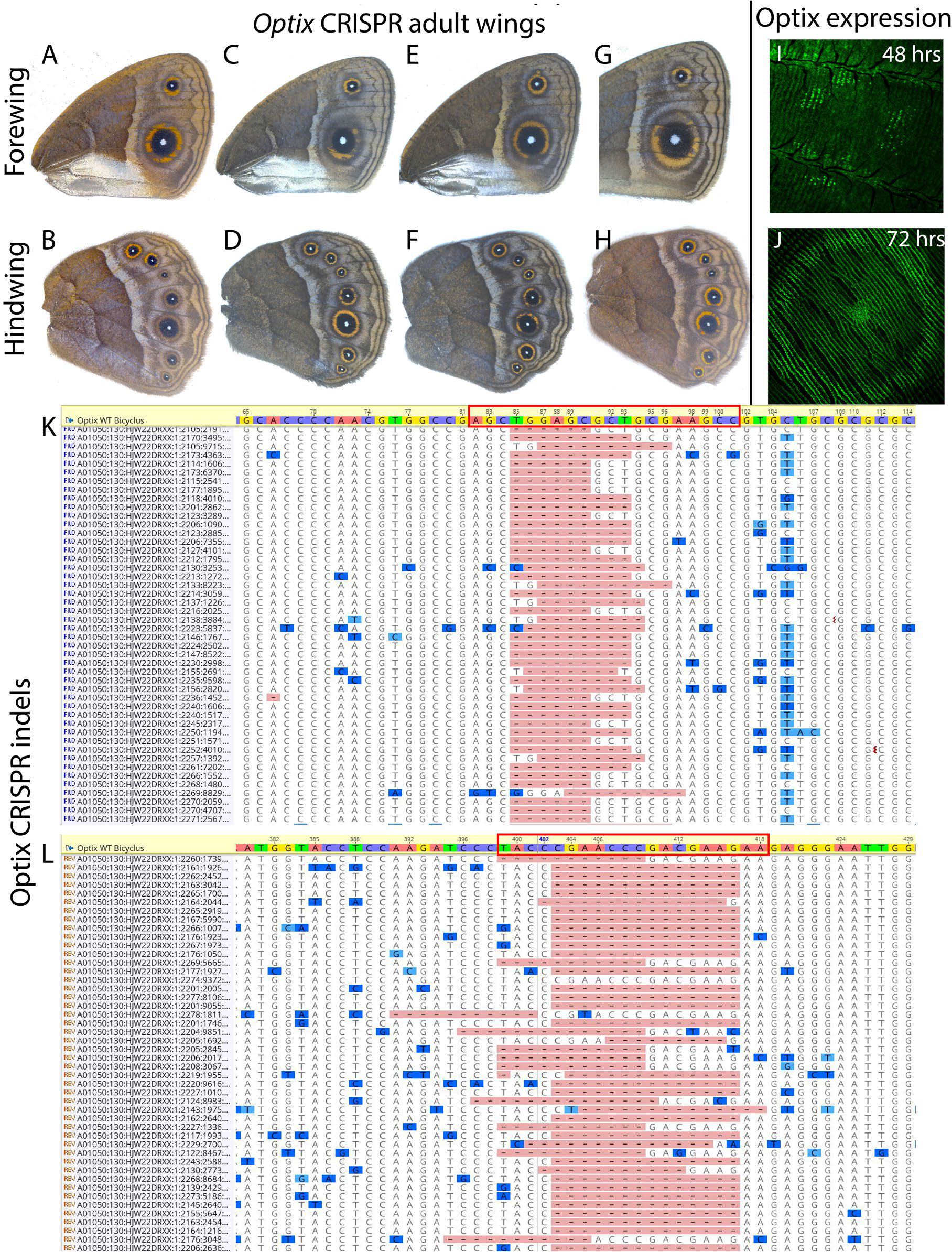
Function of *Optix* in *Bicyclus anynana* butterflies and localization of Optix protein in pupal wings. (A-H) *Optix* CRISPR adult wings. *Optix* CRISPR results in the conversion of orange scales into brown scales in the eyespots. **(I and J)** Antibody staining of Optix proteins in an *Optix* CRISPR individual at 48 hrs and a WT 72 hrs old pupal wing. **(K and L)** Deletions at the two sites targeted for *Optix* CRISPR.

**Figure S8:**
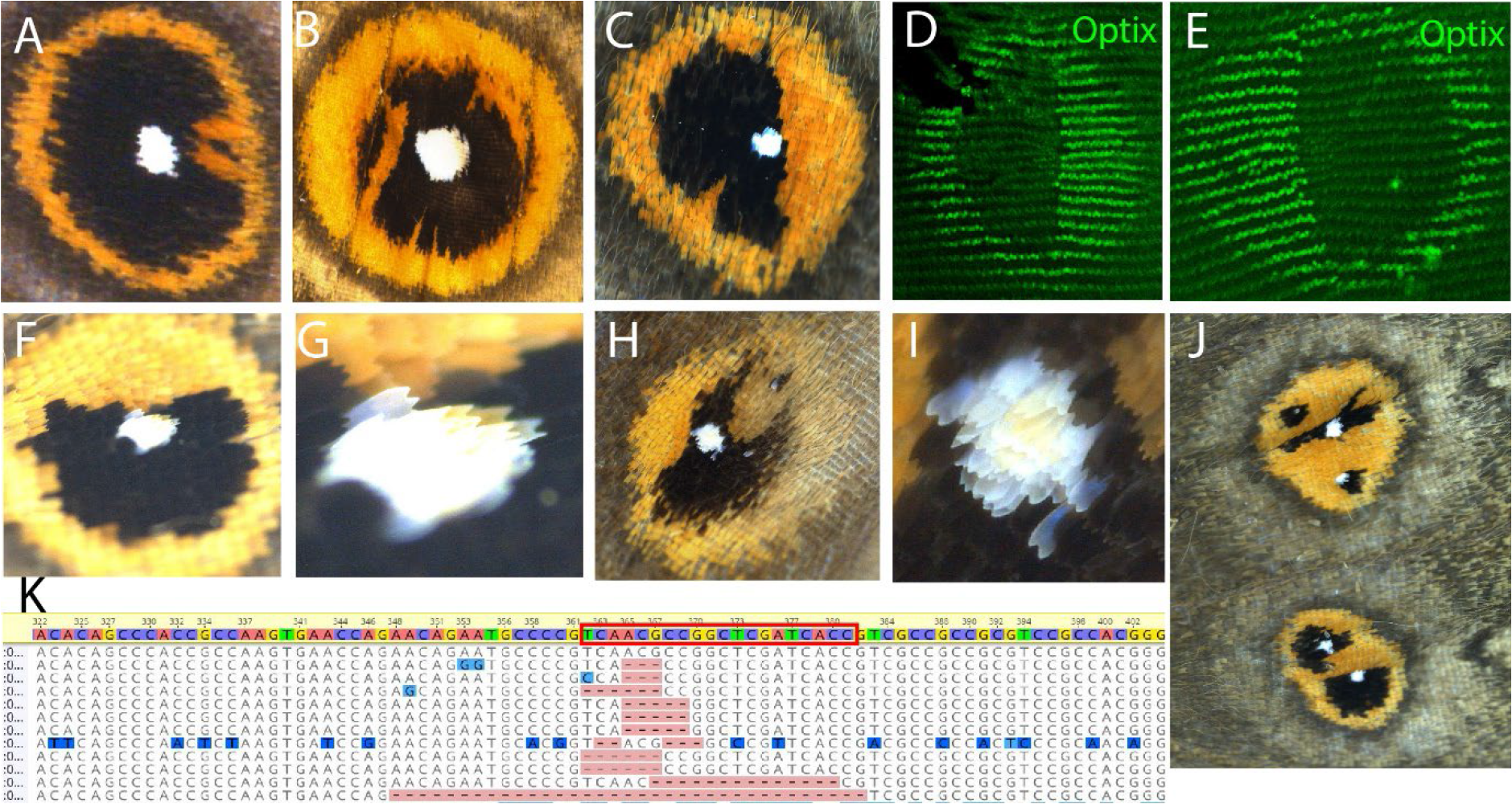
Effect of *spalt* CRISPR on the eyespots. (A-C) Loss of Spalt results in the development of orange scales in the black scale region due to the presence of (**D and E**) Optix in that region. (**F-I**) Loss of *spalt* results in the development of yellow scales in the white center scale region of the eyespot. (**J**) Loss of *spalt* results in the split of eyespot foci with distinct domains of the white, black, and orange scales. (**K**) Deletions at the site of *spalt* CRISPR (red box).

**Figure S9.**
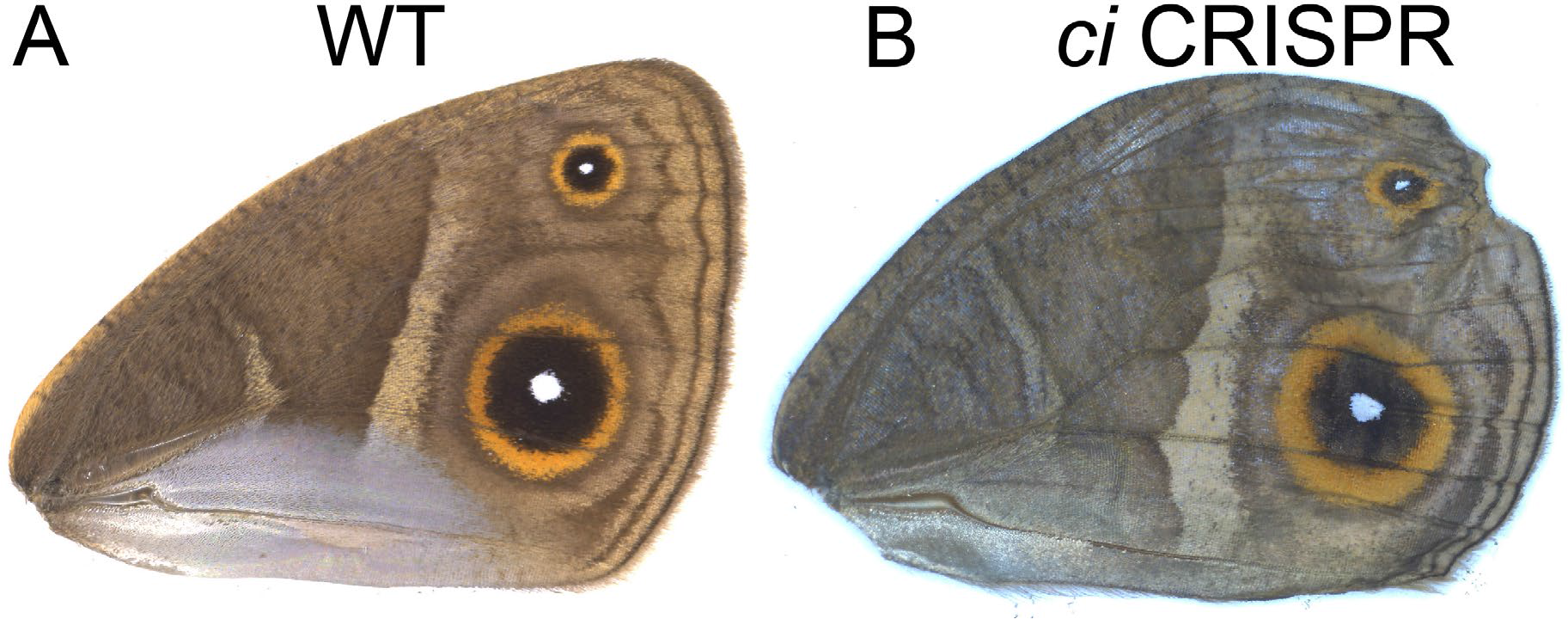
Effect of *ci* CRISPR in the adult forewing. **(A)** Adult WT forewing. (B) *ci* CRISPR forewing showing defect in the anterior compartment. No defect was observed in the eyespots.

**Figure S10.**
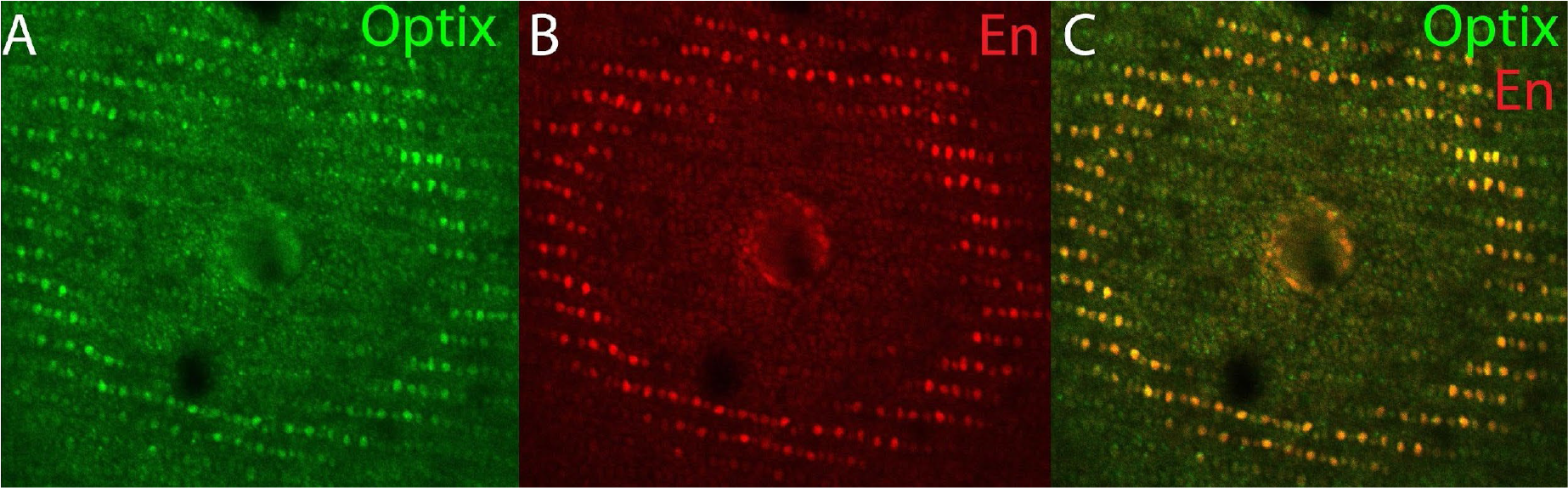
Co-localization of Optix and Engrailed (En) in the orange ring of the eyespots. Both (**A**) Optix and (**B**) En proteins are present in the cells that will form the orange scales of the eyespot. (**C**) Co-expression of Optix and En.

**Table S1:**
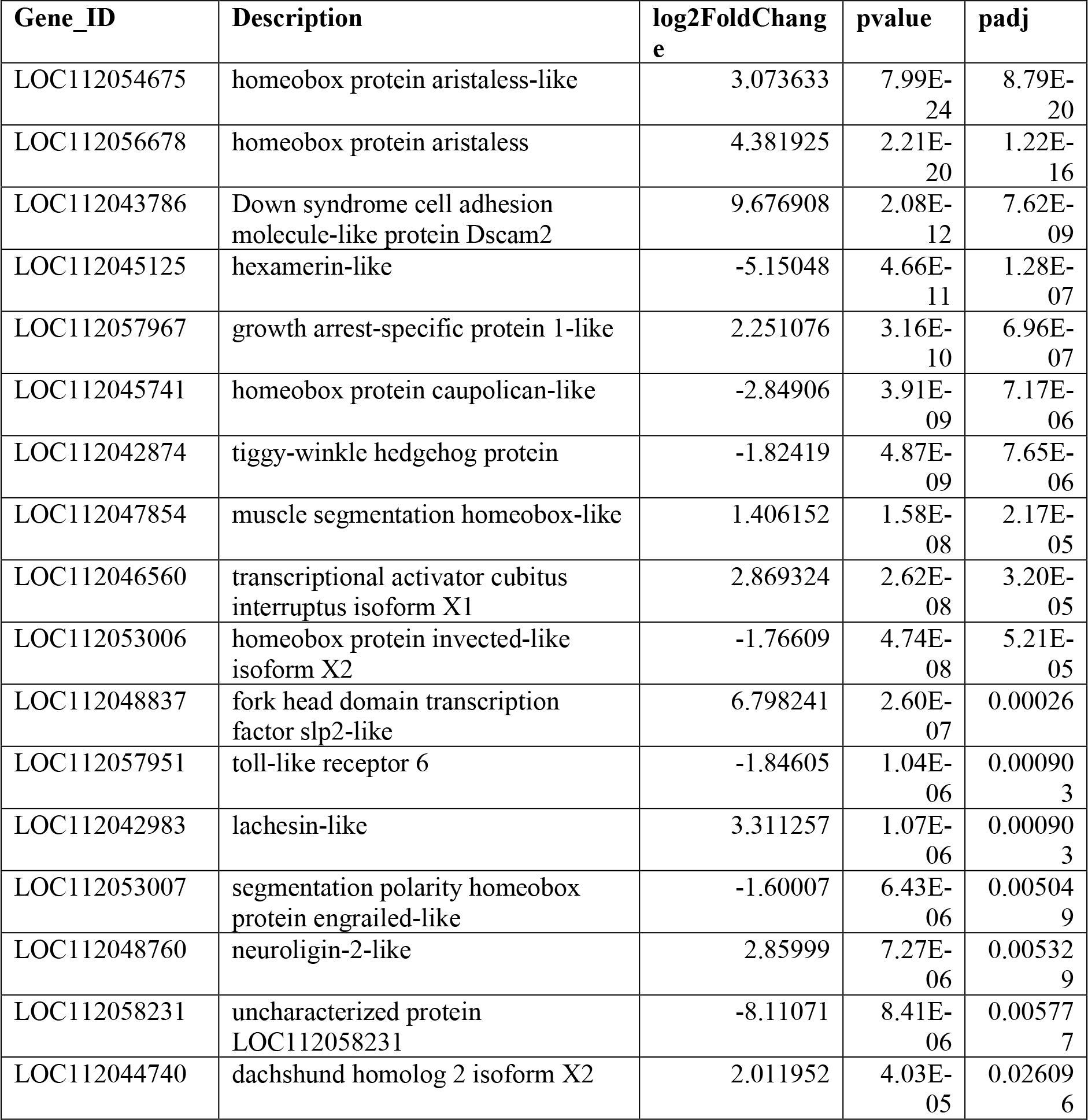
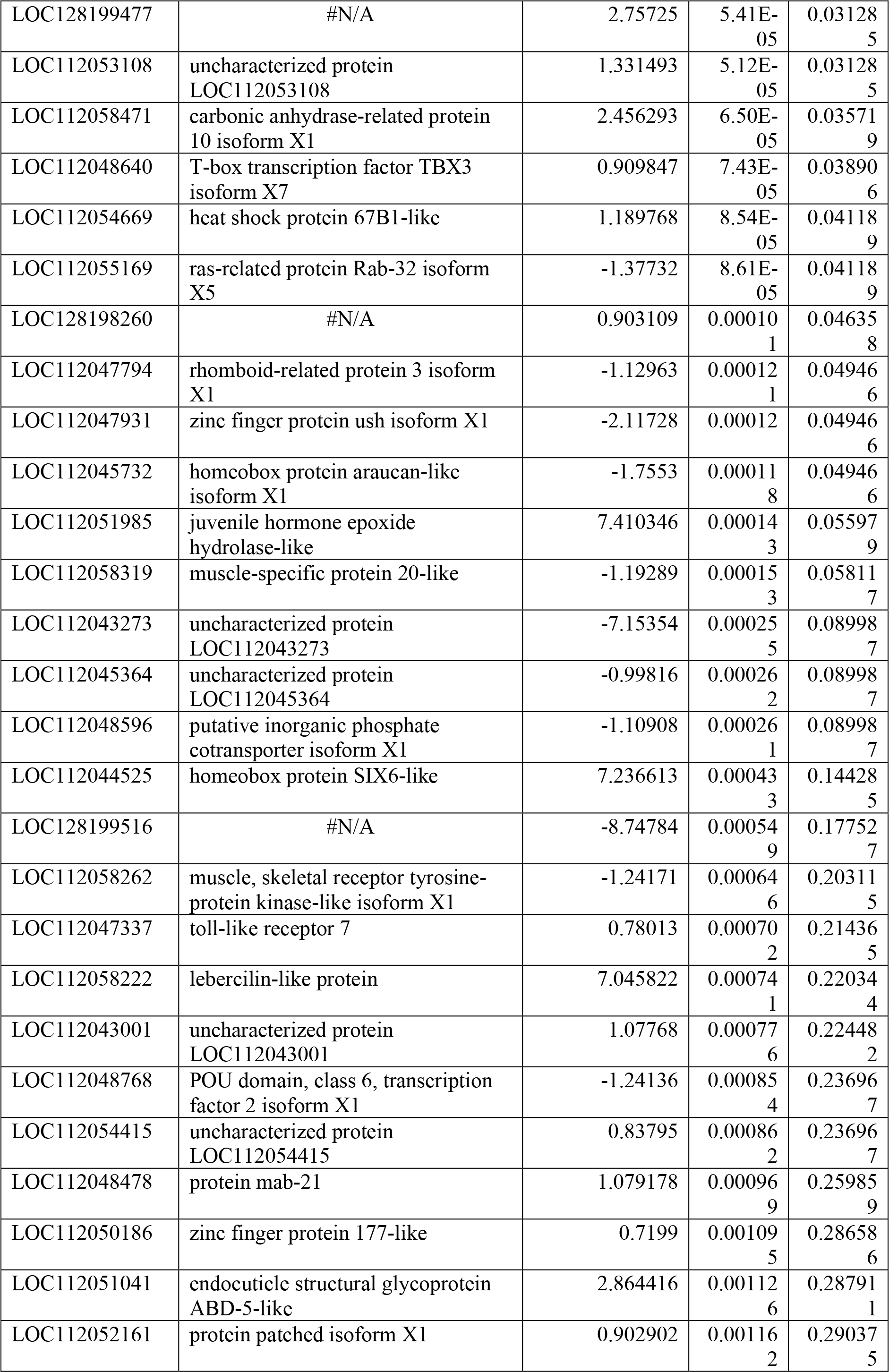
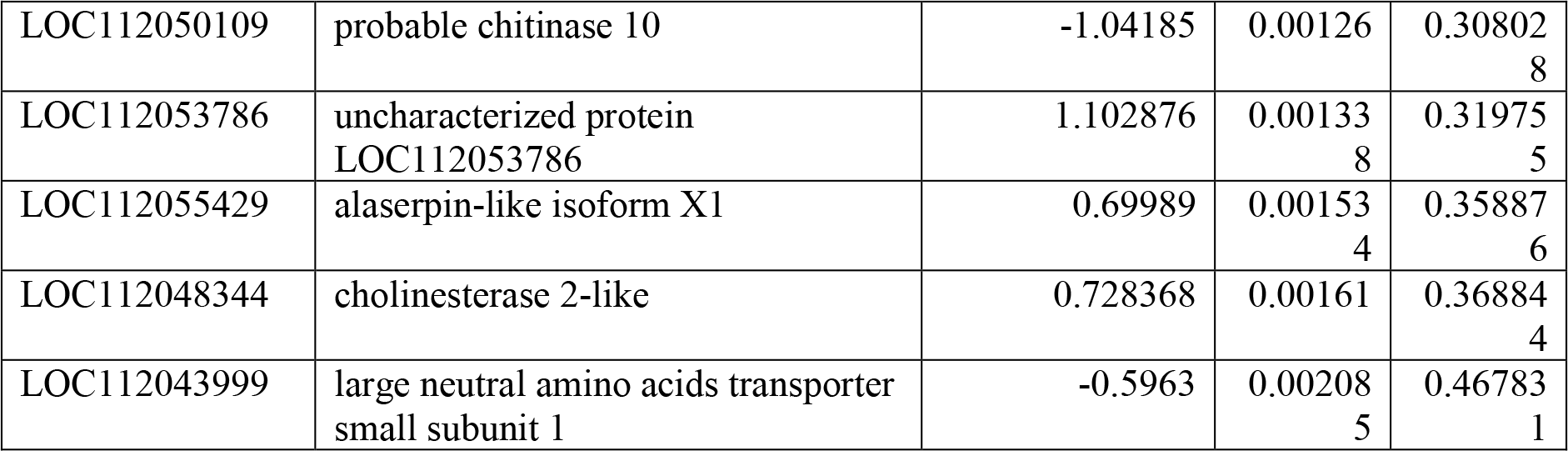
Top differentially expressed genes in the anterior and posterior compartment of larval *B. anynana* wings.

**Table S2:**
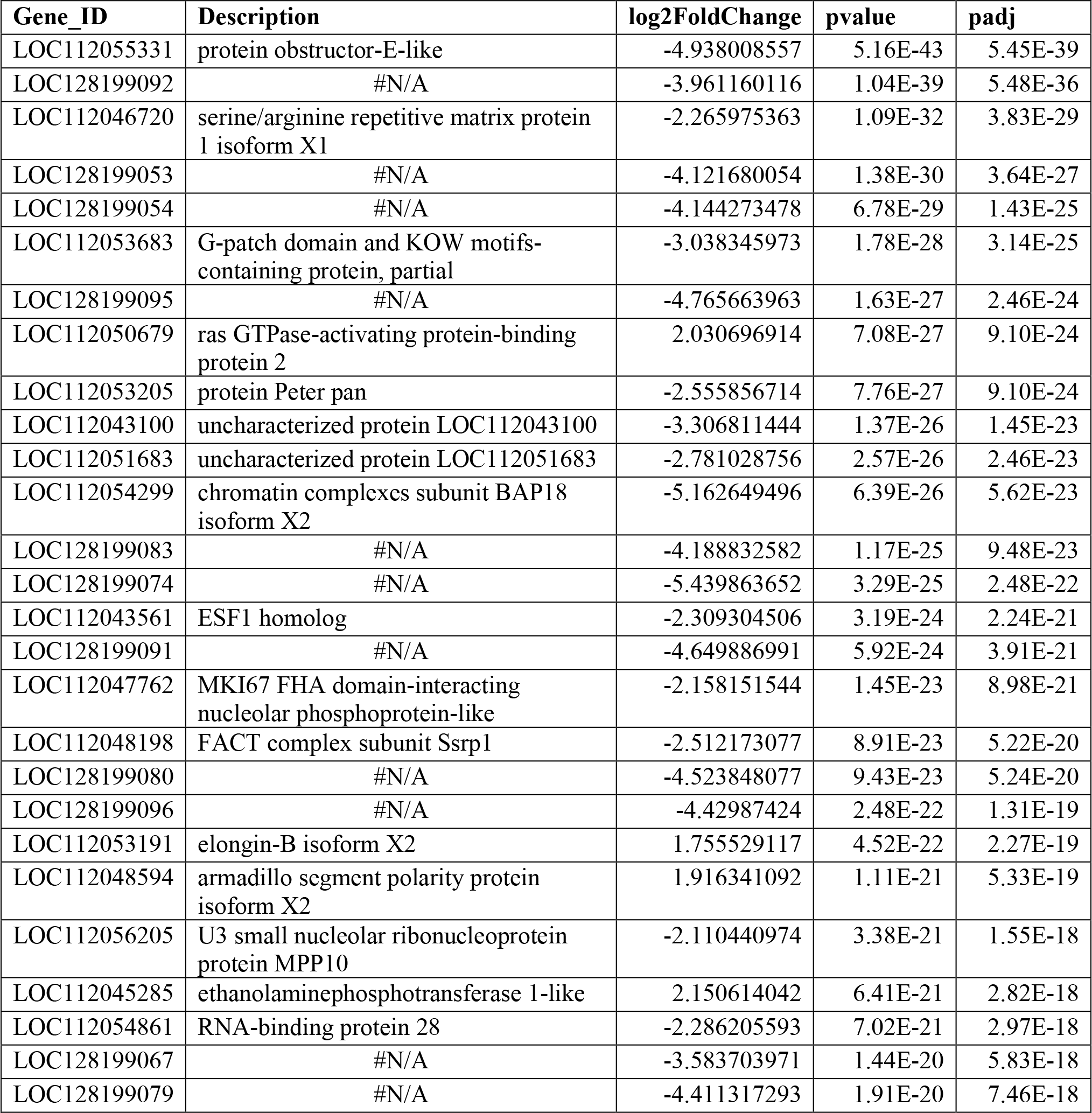
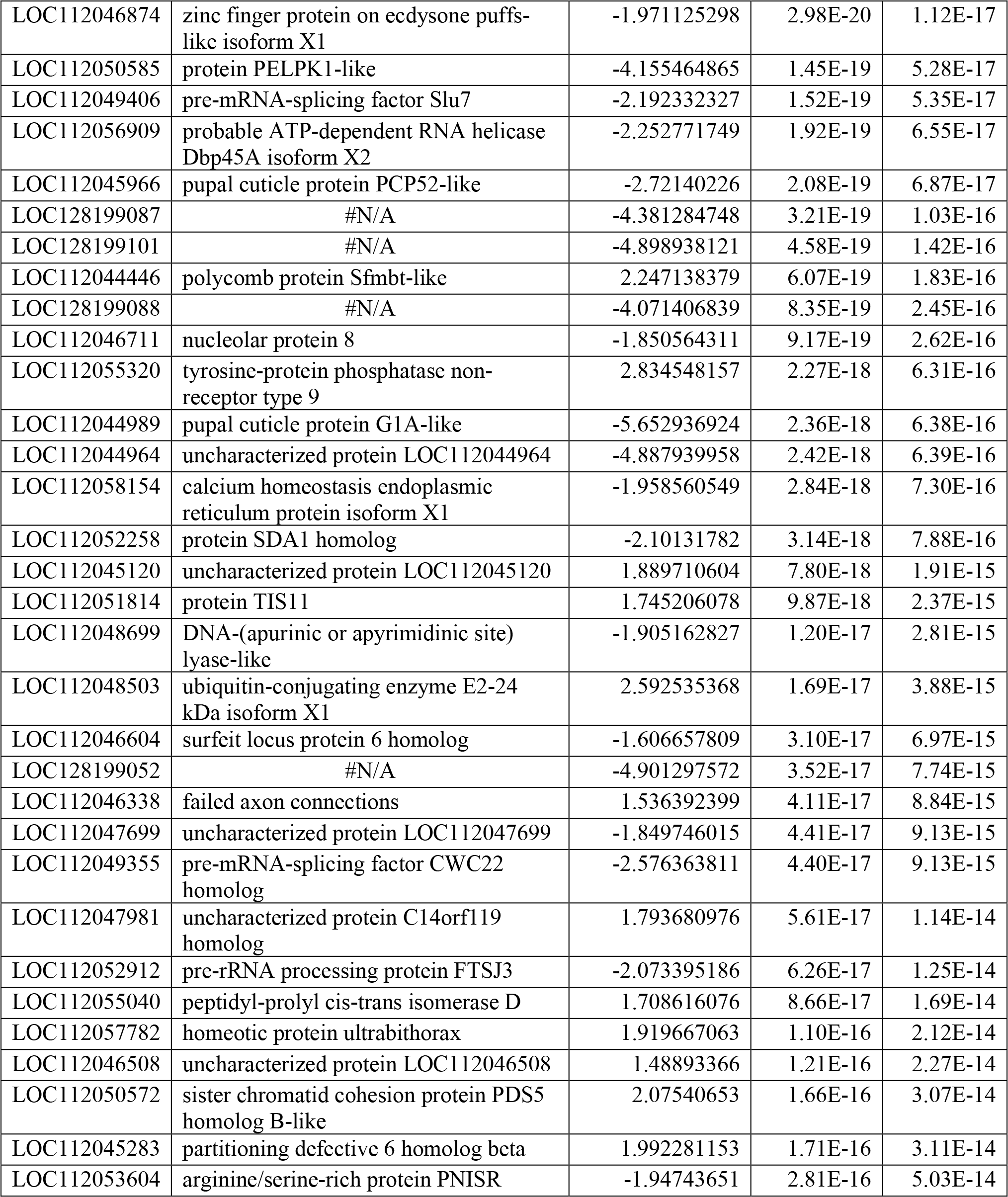
Top differentially expressed genes in the eyespot and control tissue of *B. anynana* pupal wings.

**Table S3:**
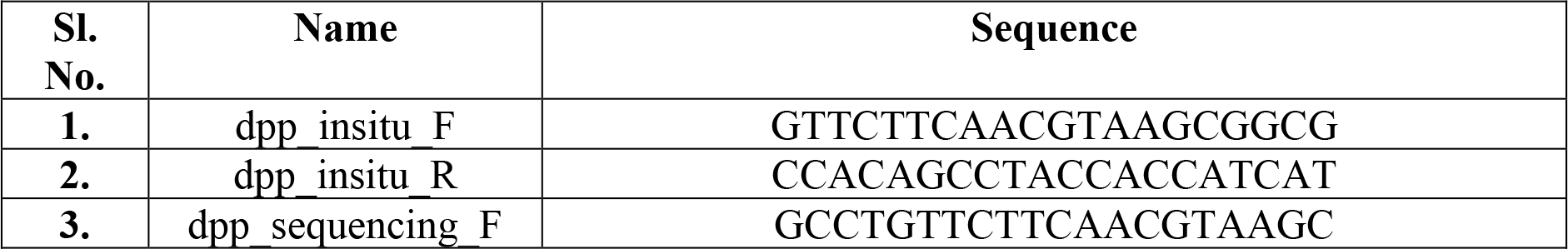
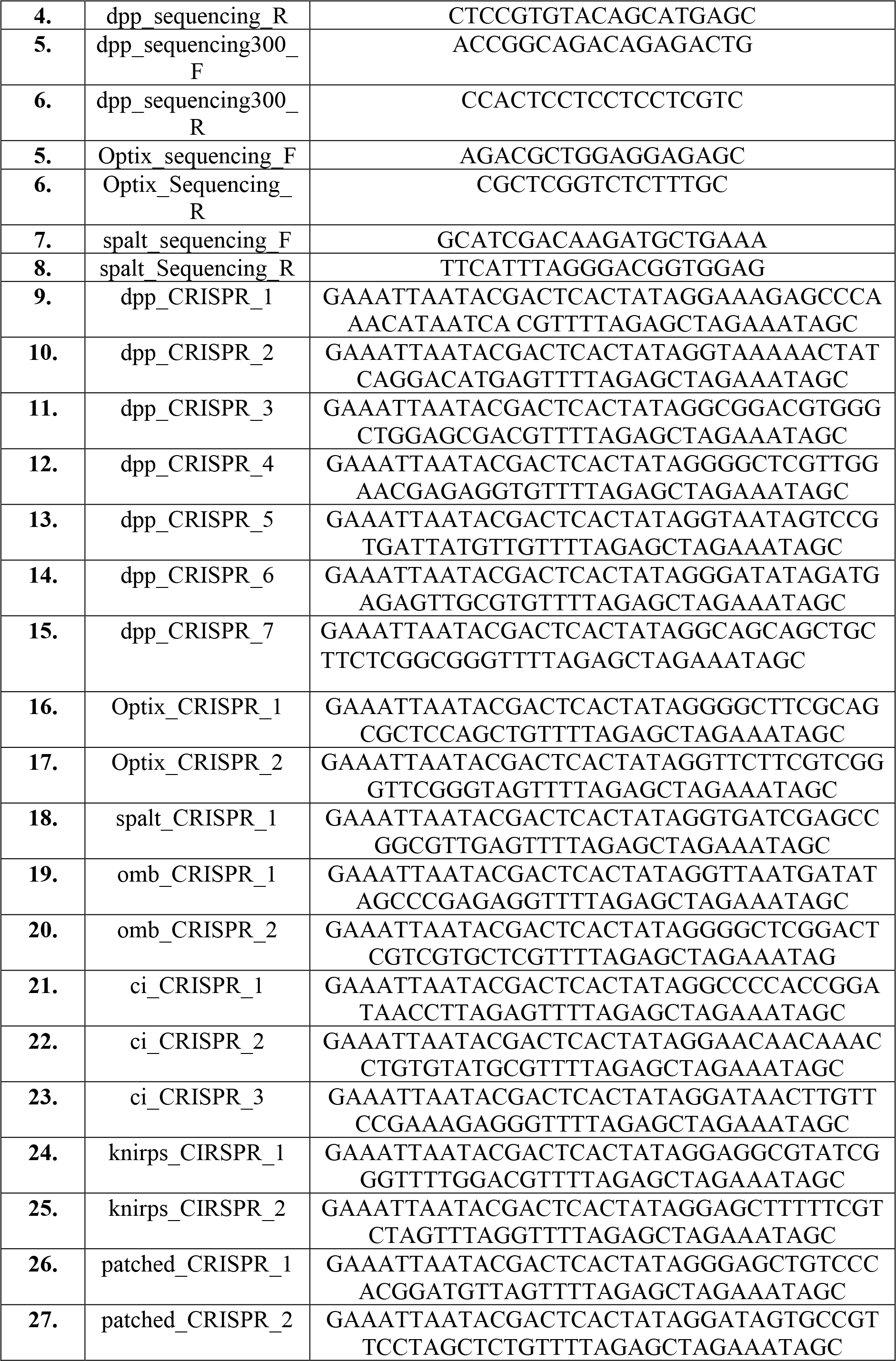
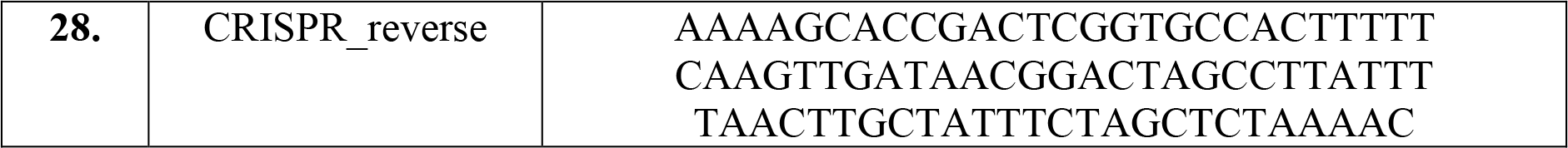
Primer table.

**Table S4.** Optix CRISPR-Cas9 injection table.

| **Sl. No.** | **Concentration** | **Date** | **Eggs Injected** | **Hatchlings** | **% Hatchlings** |
| --- | --- | --- | --- | --- | --- |
| **1.** | 300 ng/μl | 6<sup>th</sup> June 2020 | 326 | 65 | 19.9 |
| **2.** | 300 ng/μl | 9<sup>th</sup> July 2020 | 923 | 126 | 13.6 |
| **3.** | 300 ng/μl | 27<sup>th</sup> Aug 2020 | 389 | 88 | 22.6 |

**Table S5.**
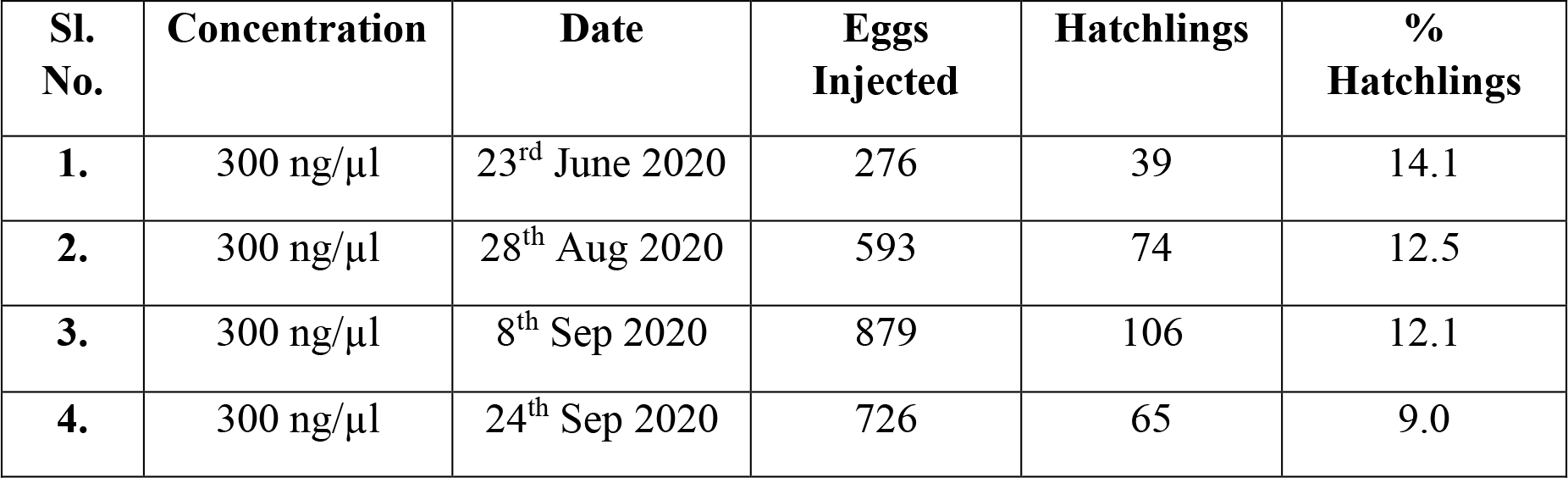
spalt CRISPR-Cas9 injection table.

| **Sl. No.** | **Concentration** | **Date** | **Eggs Injected** | **Hatchlings** | **% Hatchlings** |
| --- | --- | --- | --- | --- | --- |
| **1.** | 300 ng/μl | 23<sup>rd</sup> June 2020 | 276 | 39 | 14.1 |
| **2.** | 300 ng/μl | 28<sup>th</sup> Aug 2020 | 593 | 74 | 12.5 |
| **3.** | 300 ng/μl | 8<sup>th</sup> Sep 2020 | 879 | 106 | 12.1 |
| **4.** | 300 ng/μl | 24<sup>th</sup> Sep 2020 | 726 | 65 | 9.0 |

**Table S6.** dpp CRISPR-Cas9 injection table.

| **Sl. No.** | **Concentration** | **Date** | **Eggs Injected** | **Hatchlings** | **% Hatchlings** |
| --- | --- | --- | --- | --- | --- |
| **1.** | 300 ng/μl | 23<sup>rd</sup> Jan 2020 | 683 | 89 | 13.0 |
| **2.** | 300 ng/μl | 3<sup>rd</sup> March 2020 | 923 | 117 | 12.7 |
| **3.** | 300 ng/μl | 11<sup>th</sup> Feb 2021 | 282 | 51 | 18.1 |
| **4.** | 300 ng/μl | 6<sup>th</sup> July 2021 | 124 | 15 | 12.1 |
| **5.** | 300 ng/μl | 23<sup>rd</sup> July 2021 | 472 | 29 | 6.14 |
| **6.** | 300 ng/μl | 27<sup>th</sup> July 2021 | 326 | 43 | 13.2 |
| **7.** | 50ng/μl | 6<sup>th</sup> Sept 2022 | 472 | 65 | 13.7 |
| **8.** | 100 ng/μl | 27<sup>th</sup> Sept 2022 | 326 | 57 | 17.5 |
| **9.** | 100ng/μl | 9<sup>th</sup> Feb 2022 | 186 | 24 | 12.9 |

**Table S7.** ci CRISPR-Cas9 injection table.

| **Sl. No.** | **Concentration** | **Date** | **Eggs Injected** | **Hatchlings** | **% Hatchlings** |
| --- | --- | --- | --- | --- | --- |
| **1.** | 300 ng/μl | 8<sup>th</sup> July 2021 | 283 | 35 | 12.0 |

**Table S8.** knirps CRISPR-Cas9 injection table.

| **Sl. No.** | **Concentration** | **Date** | **Eggs Injected** | **Hatchlings** | **% Hatchlings** |
| --- | --- | --- | --- | --- | --- |
| **1.** | 300 ng/μl | 15<sup>th</sup> Feb 2020 | 725 | 128 | 17.6 |
| **2.** | 300 ng/μl | 12<sup>th</sup> Feb 2020 | 534 | 97 | 18.2 |

**Table S9.** omb CRISPR-Cas9 injection table.

| **Sl. No.** | **Concentration** | **Date** | **Eggs Injected** | **Hatchlings** | **% Hatchlings** |
| --- | --- | --- | --- | --- | --- |
| **1.** | 300 ng/μl | 9<sup>th</sup> Feb 2020 | 734 | 85 | 11.6 |

**Table S10.** patched CRISPR-Cas9 injection table.

| **Sl. No.** | **Concentration** | **Date** | **Eggs Injected** | **Hatchlings** | **% Hatchlings** |
| --- | --- | --- | --- | --- | --- |
| **1.** | 300 ng/μl | 23<sup>rd</sup> Nov 2020 | 306 | 78 | 25.5 |

**Table S11.** Immunofluorescence Buffers.

| **Buffers** | **Chemicals** | **Amount** |
| --- | --- | --- |
| Fix buffer (30 ml) | 0.1M PIPES pH 6.9 (500 mM) | 6 ml |
| | 1 mM EGTA pH 6.9 (500mM) | 60 μl |
|  | 1% Triton x-100 (20 %) | 1.5 ml |
| | 2 mM MgSO<sub>4</sub> (1M) | 60 μl |
| | 37% Formaldehyde | 55 μl per 500 μl of buffer |
|  | dH<sub>2</sub>O | 22.4 ml |
| Block buffer (40 ml) | 50 mM Tris pH 6.8 (1 M) | 2 ml |
|  | 150 mM NaCl (5 M) | 1.2 ml |
|  | 0.5% IGEPAL (NP40) (20%) | 1 ml |
|  | 5 mg/ml BSA | 0.2 gr |
|  | H<sub>2</sub>O | 35.8 ml |
| Wash buffer (200 ml) | 50mM Tris pH 6.8 (1 M) | 10 ml |
|  | 150 mM NaCl (5 M) | 6 ml |
|  | 0.5% IGEPAL (20 %) | 5 ml |
|  | 1 mg/ml BSA | 0.2 gr |
|  | dH<sub>2</sub>O | 179 ml |
| Mounting media | Tris-HCl (pH 9.2) | 20 mM |
|  | N-propyl gallate | 0.5% |
|  | Glycerol | 60% |

**Table S12.** In-situ hybridization Buffers.

| Buffers | Chemicals | Amount |
| --- | --- | --- |
| 10X PBS (500 ml)<br>* Sterilize by autoclaving. | K<sub>2</sub>HPO<sub>4</sub> | 5.34 g |
|  | KH<sub>2</sub>PO<sub>4</sub> | 2.64 g |
|  | NaCl | 40.9 g |
|  | DEPC treated H<sub>2</sub>O | To 500 ml |
| 1X PBST (50 ml) | 1X PBS | 50 ml |
|  | Tween® 20 | 50 µl |
| 20X SSC (1000 ml)<br>*Adjust the pH to 7.0 with 1M HCl and sterilize by autoclaving. | NaCl | 175.3 g |
|  | Trisodium citrate | 88.2 g |
|  | DEPC treated H<sub>2</sub>O | Till 1000 ml |
| Pre-hybridization buffer (40 ml) | Formamide | 20 ml |
|  | 20X SSC | 10 ml |
|  | DEPC treated water | 10 ml |
|  | TWEEN20 | 40 µl |
| Hybridization buffer (40 ml) | Formamide | 20 ml |
|  | 20X SSC | 10 ml |
|  | DEPC treated water | 10 ml |
|  | TWEEN20 | 40 µl |
|  | Spaltmon sperm | 40 µl |
|  | Glycine (100mg/ml) | 40 µl |
| Block buffer (50 ml) | 1X PBS | 50 ml |
|  | TWEEN20 | 50 µl |
|  | BSA | 0.1 gm |
| Alkaline phosphatase buffer (20 ml) | Tris-HCl (pH 8.0) | 2 ml |
|  | NaCl (5M) | 400 µl |
|  | MgCl<sub>2</sub> (200mM) | 250 µl |
|  | DEPC treated water | Till 20 ml |
|  | TWEEN20 | 20 µl |

## Sequence of *decapentaplegic* used for enzyme based *in-situ* hybridization

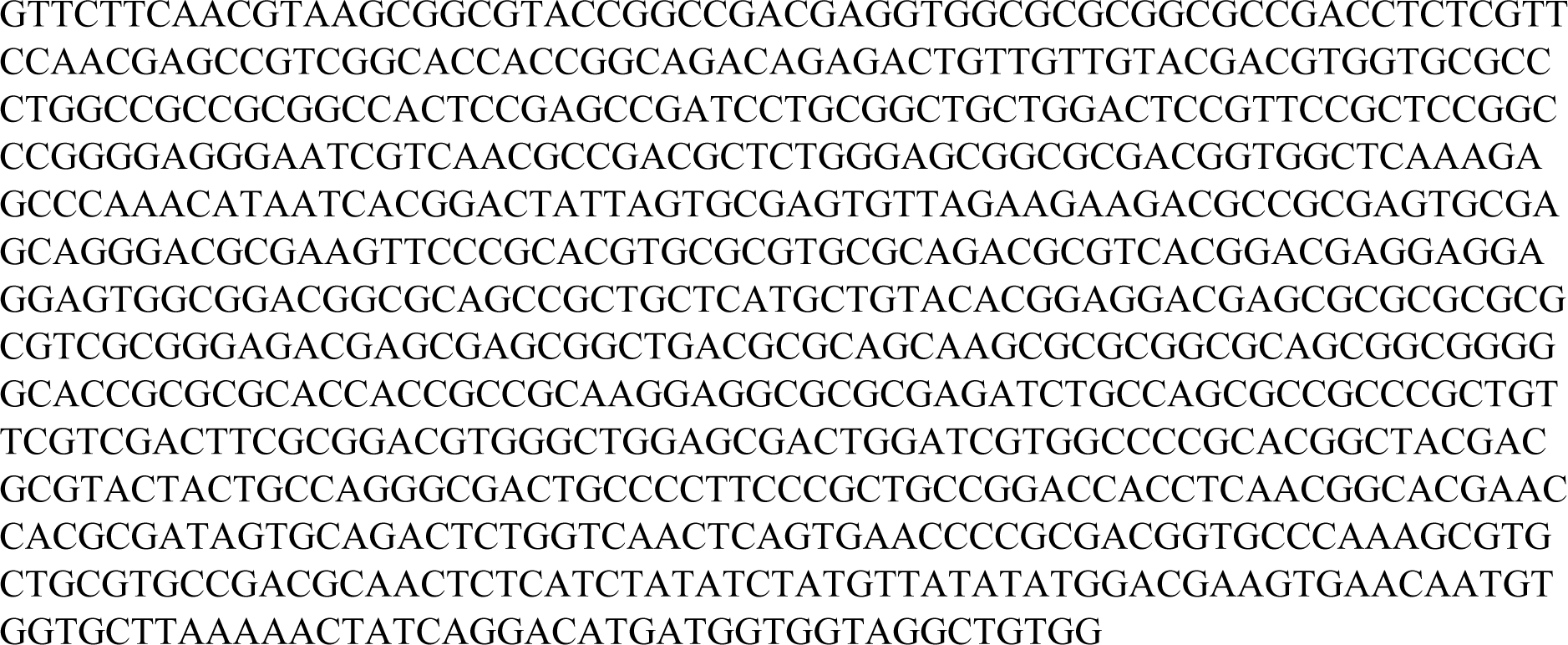

## Region of *Optix* used for CRISPR-Cas9 (Highlighted in red)

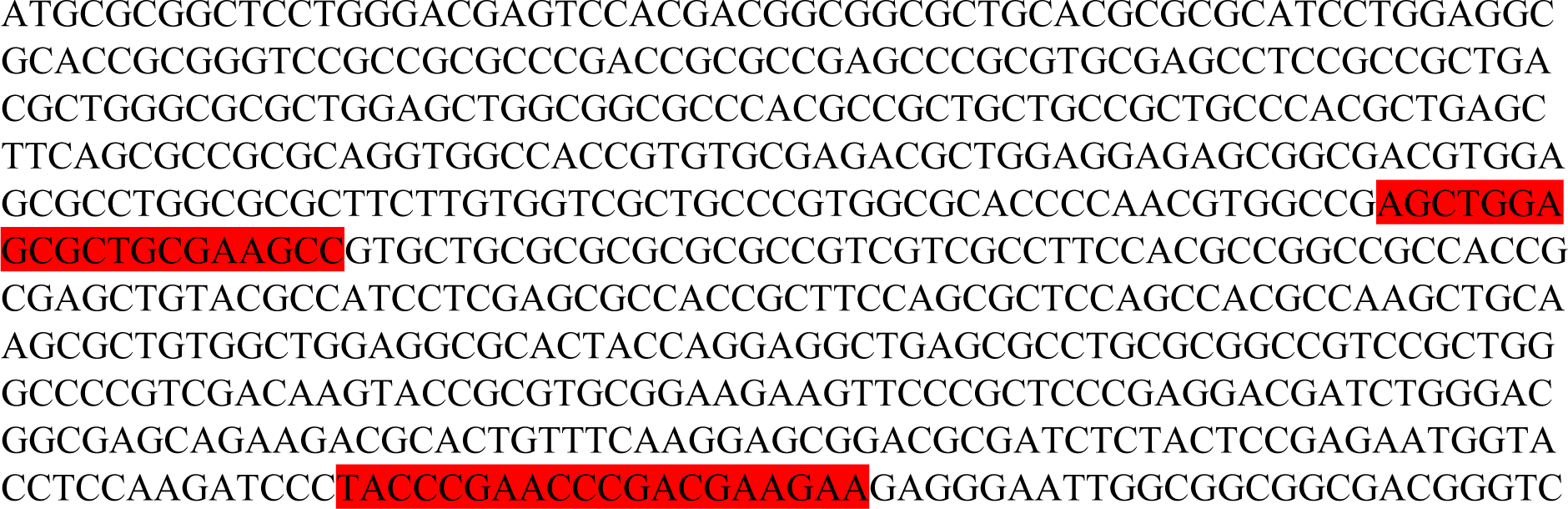
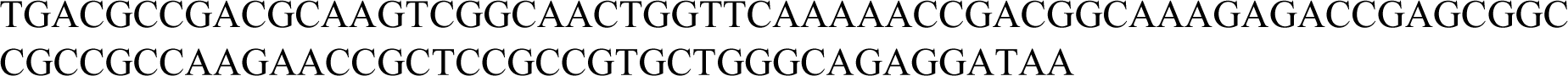

## Region of *spalt* targeted by CRISPR-Cas9 (location of guide RNA highlighted in red)

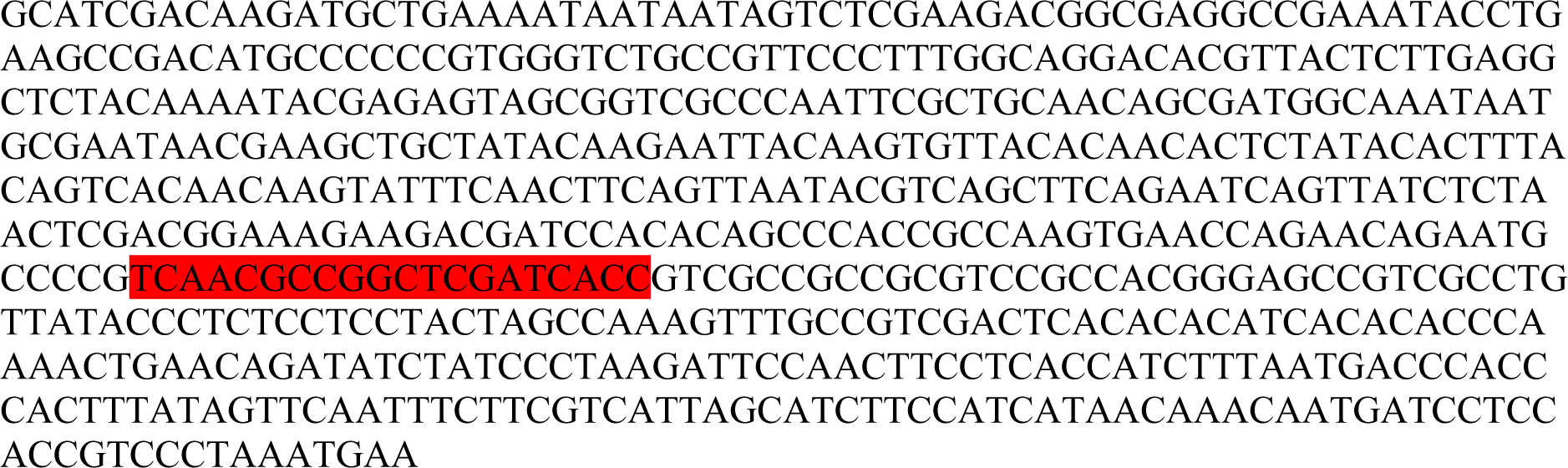

## Region of *dpp* used for CRISPR-Cas9 (Highlighted in red)

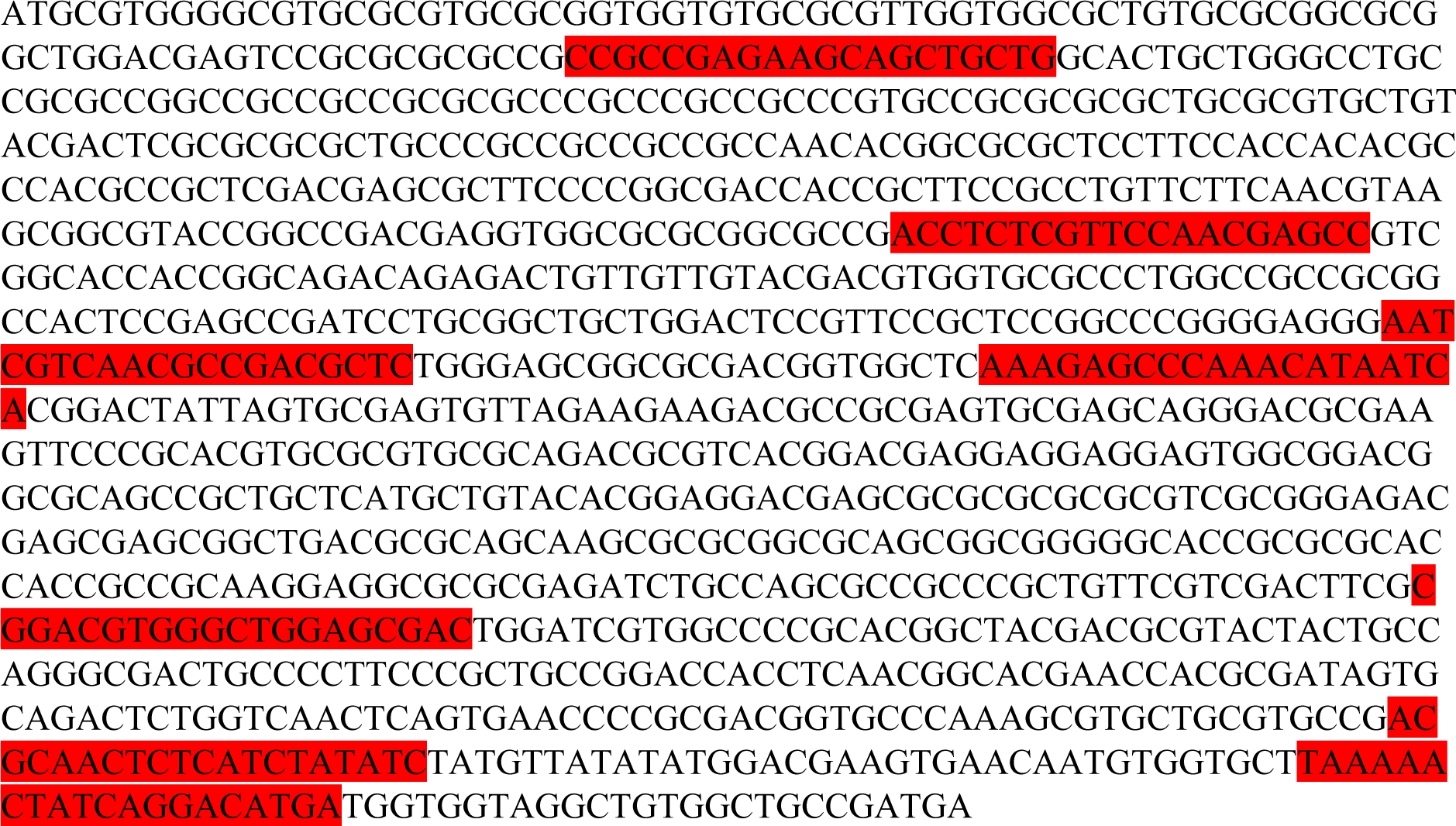

## Region of *omb* used for CRISPR-Cas9 (Highlighted in red)

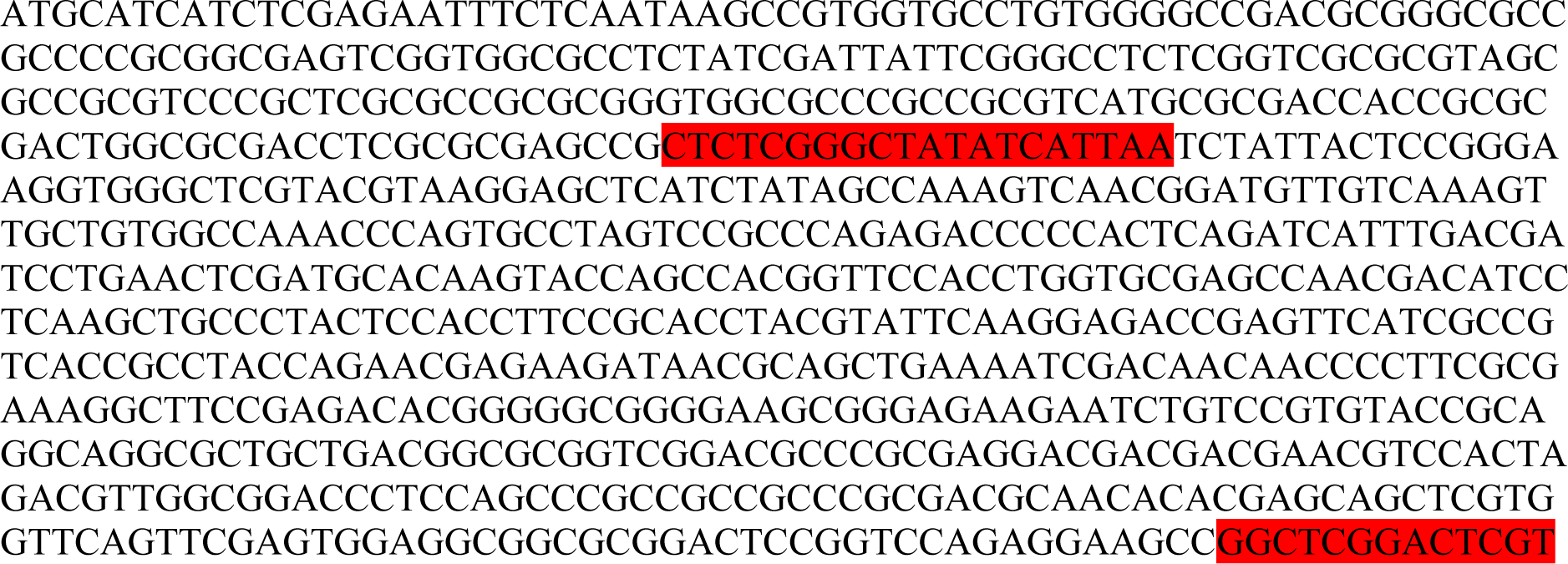
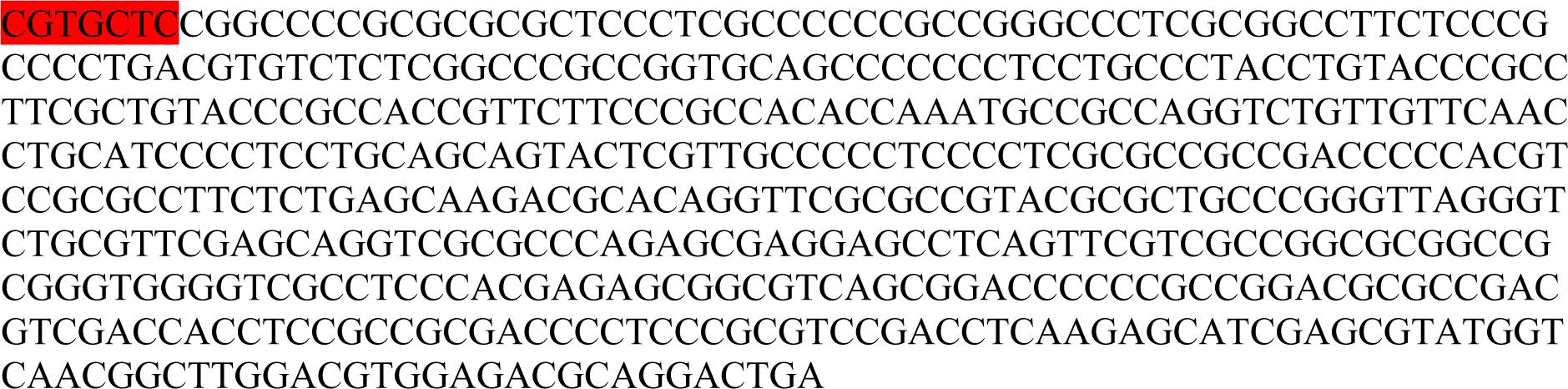

## Region of *ci* used for CRISPR-Cas9 (Highlighted in red)

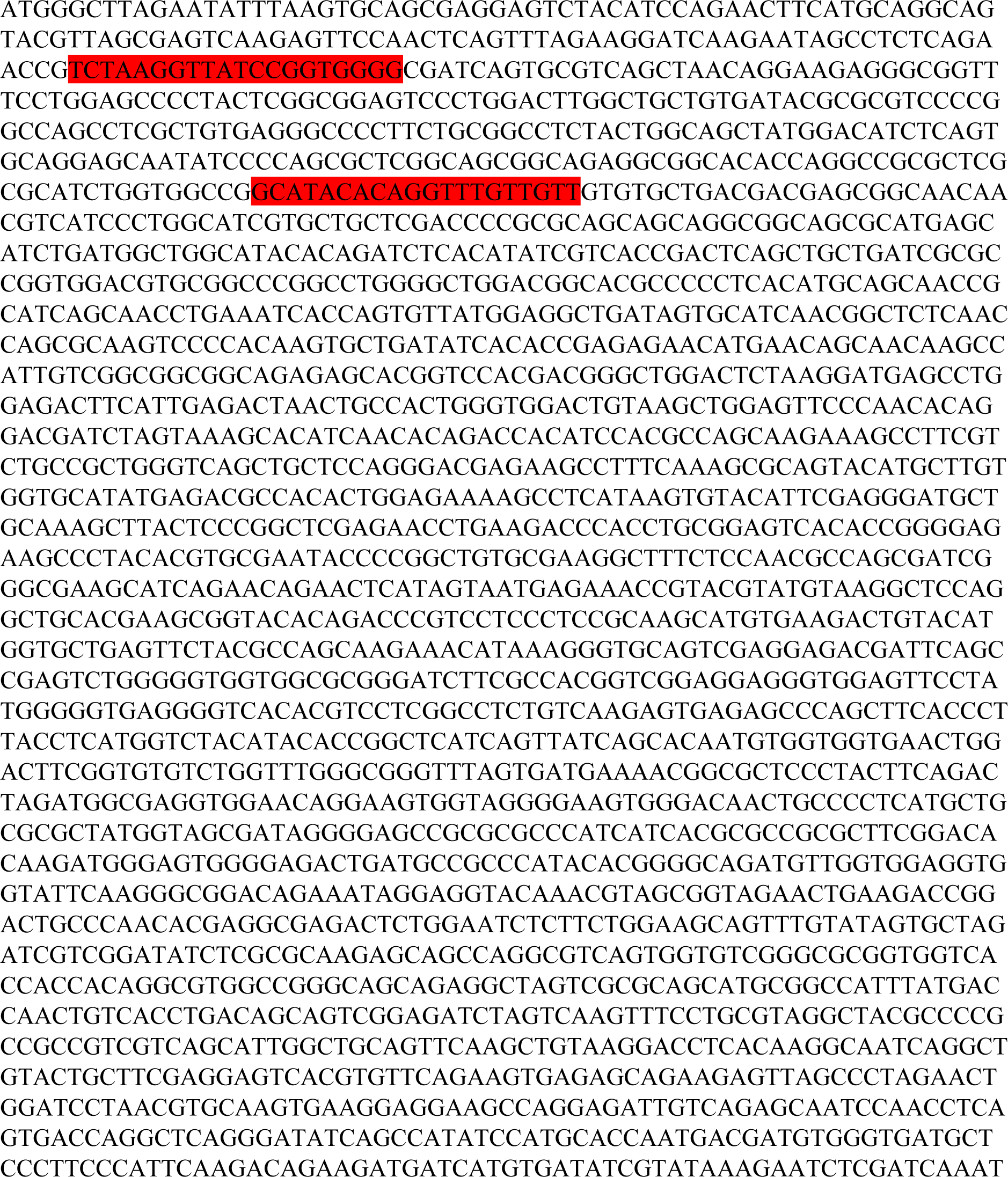
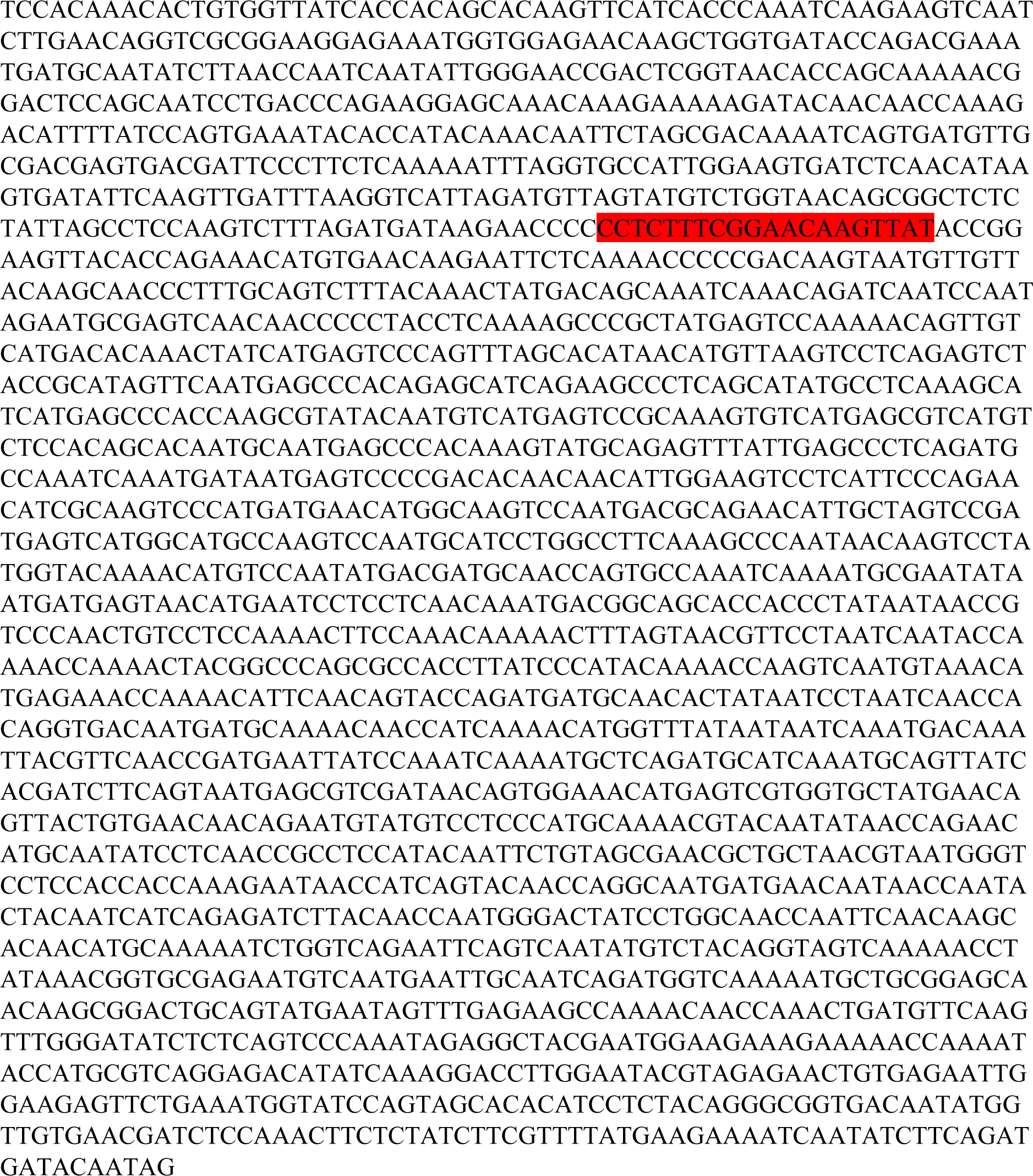

## Region of *knirps* used for CRISPR-Cas9 (Highlighted in red)

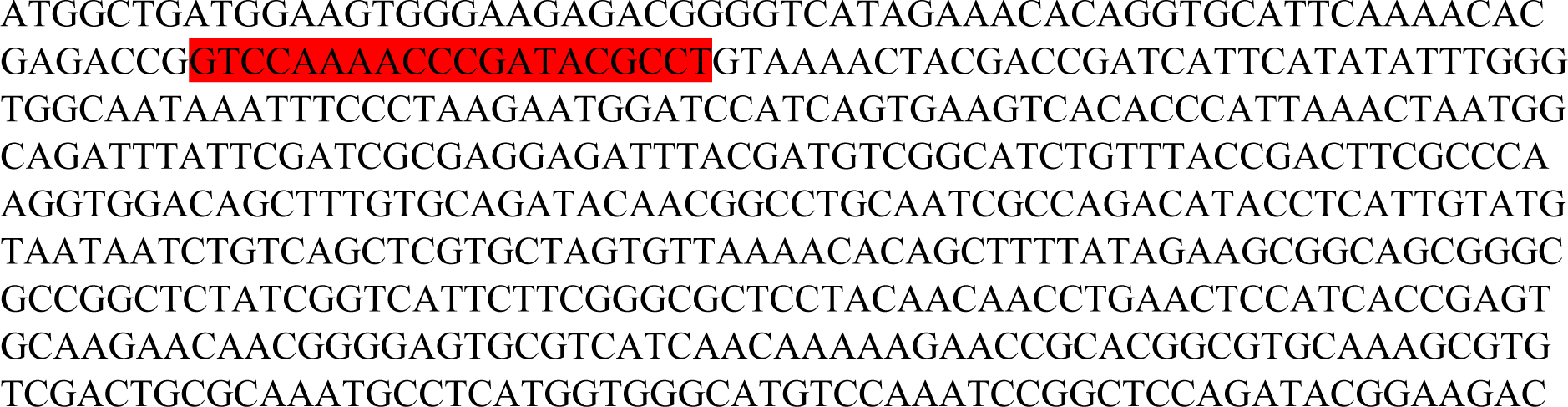
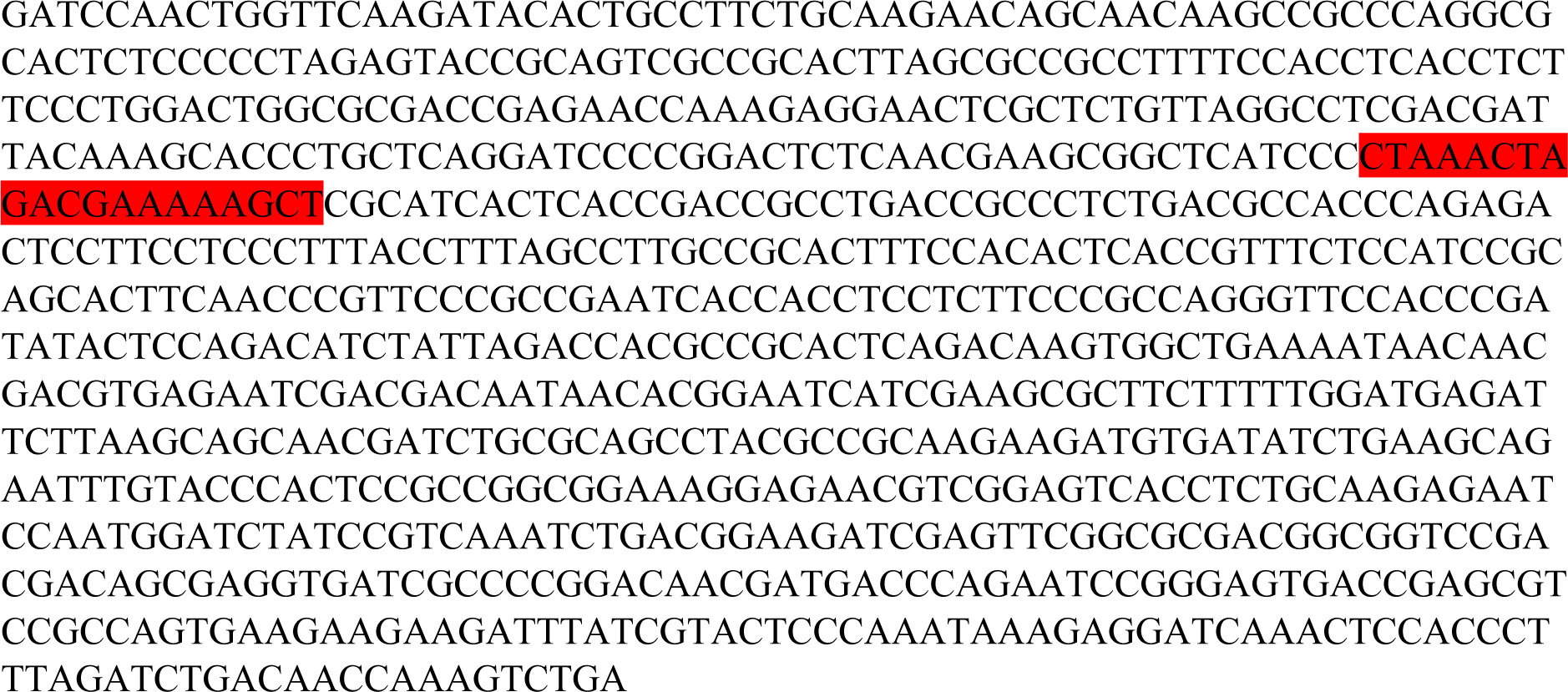

## Region of *ptc* used for CRISPR-Cas9 (Highlighted in red)

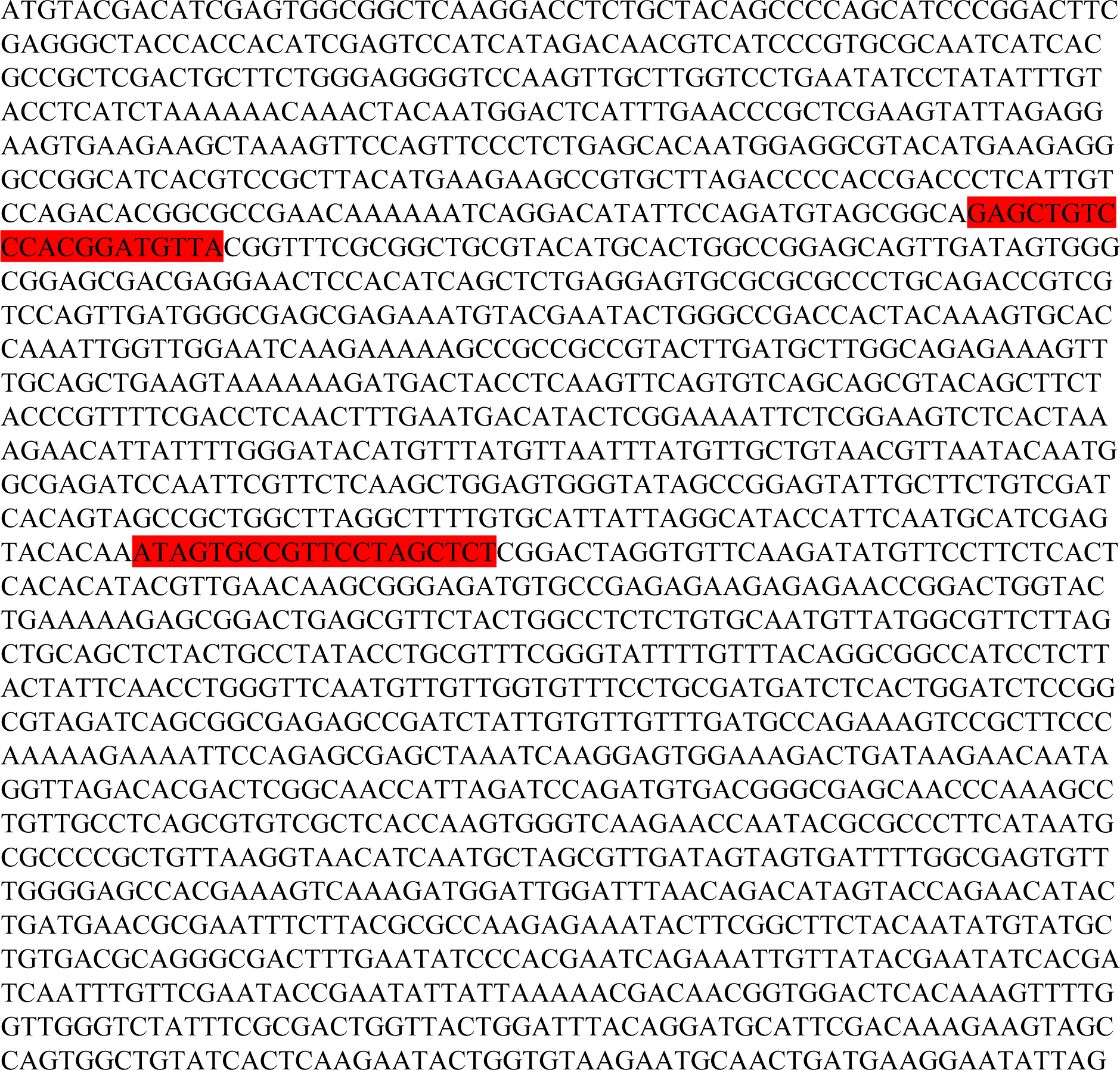
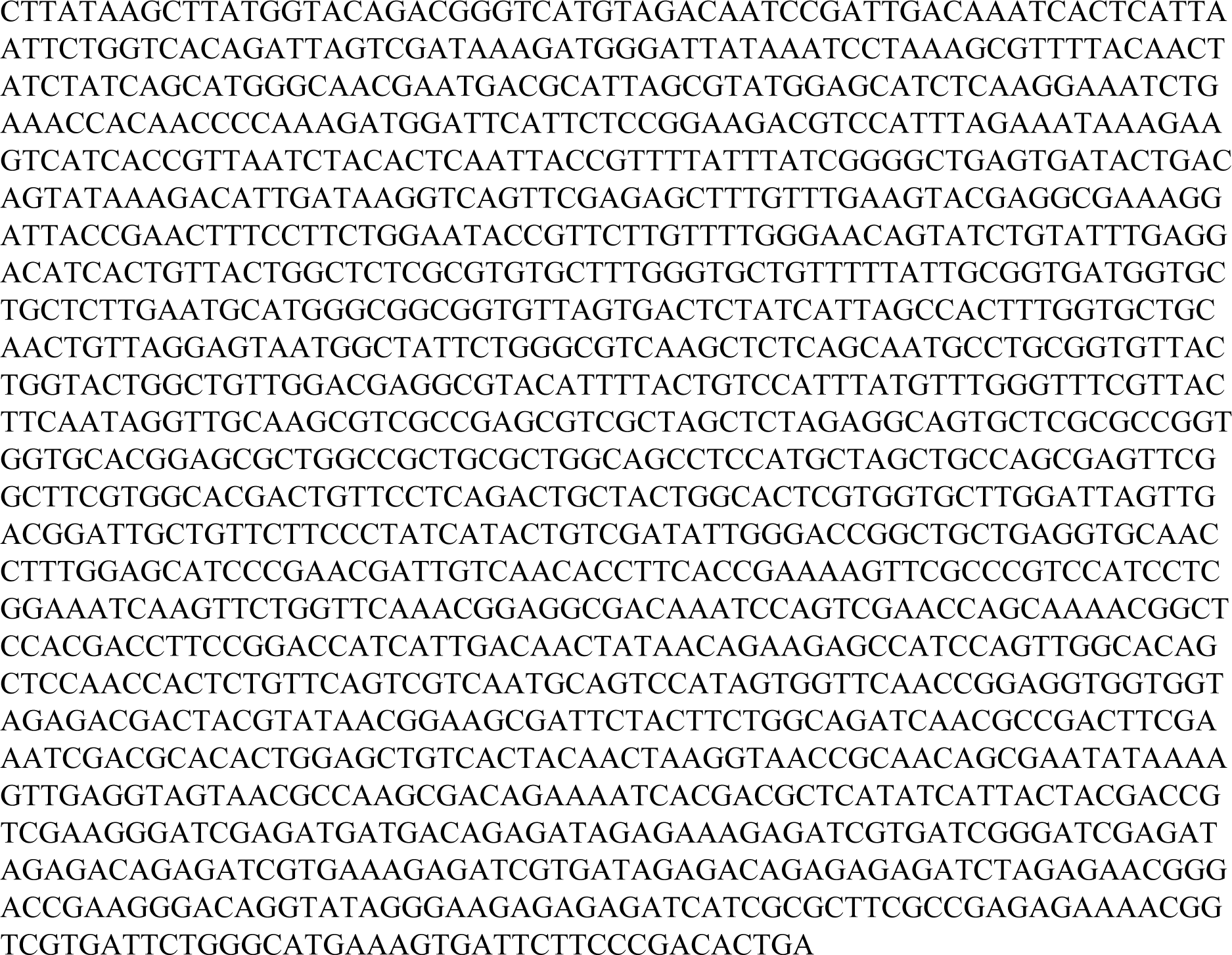

## Sequence of the genes and the oligos used for HCR3.0

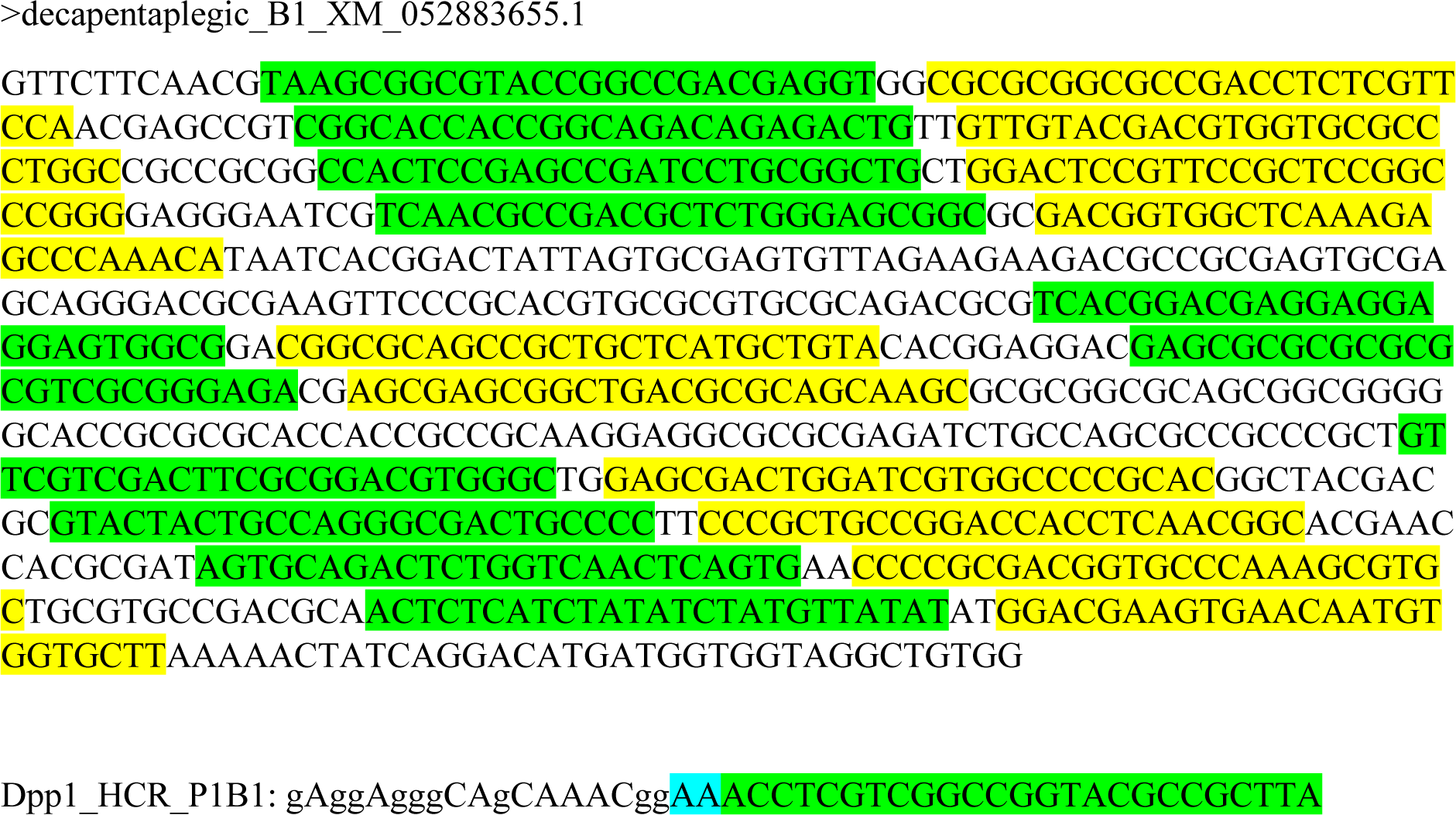
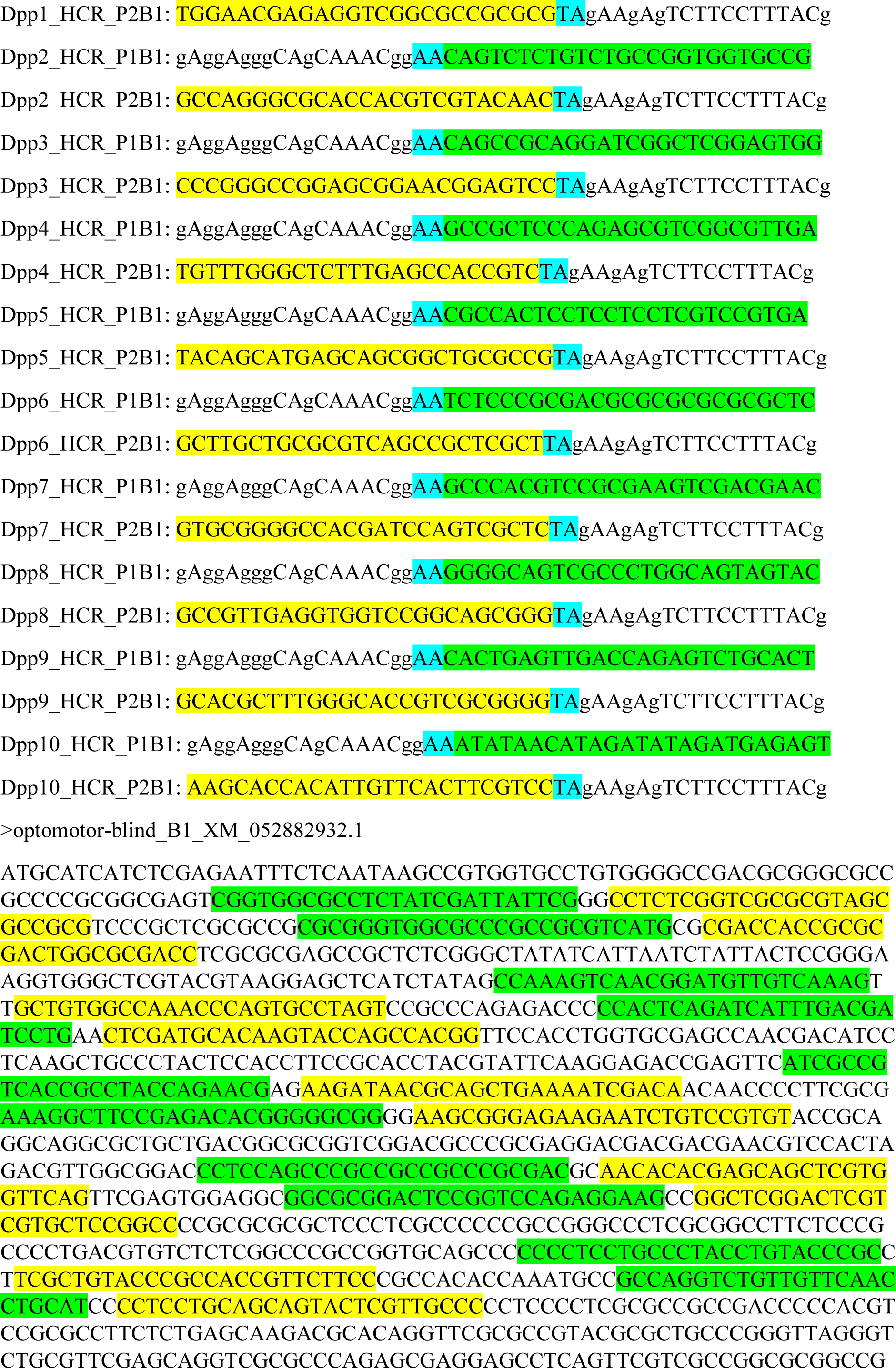
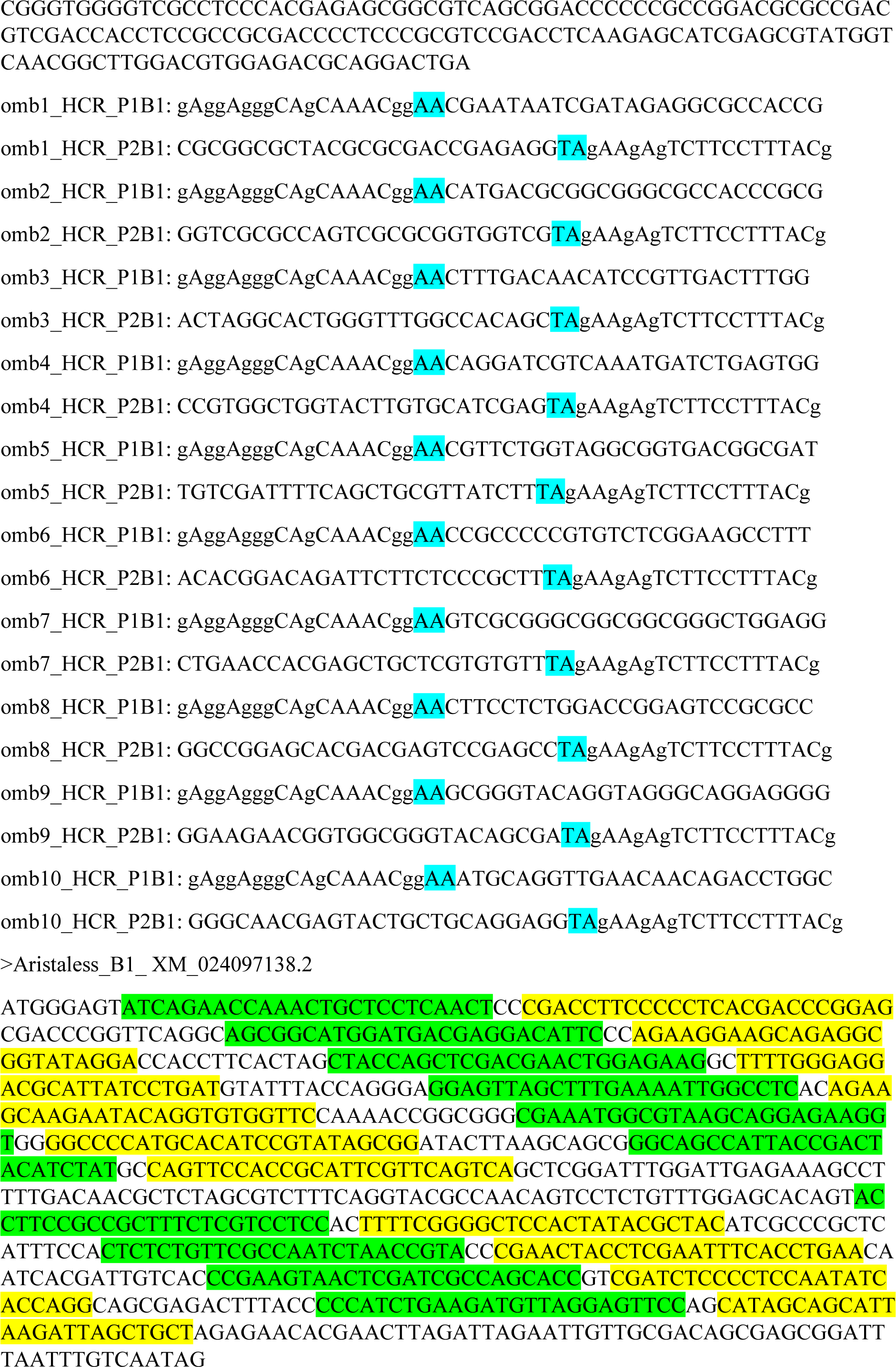
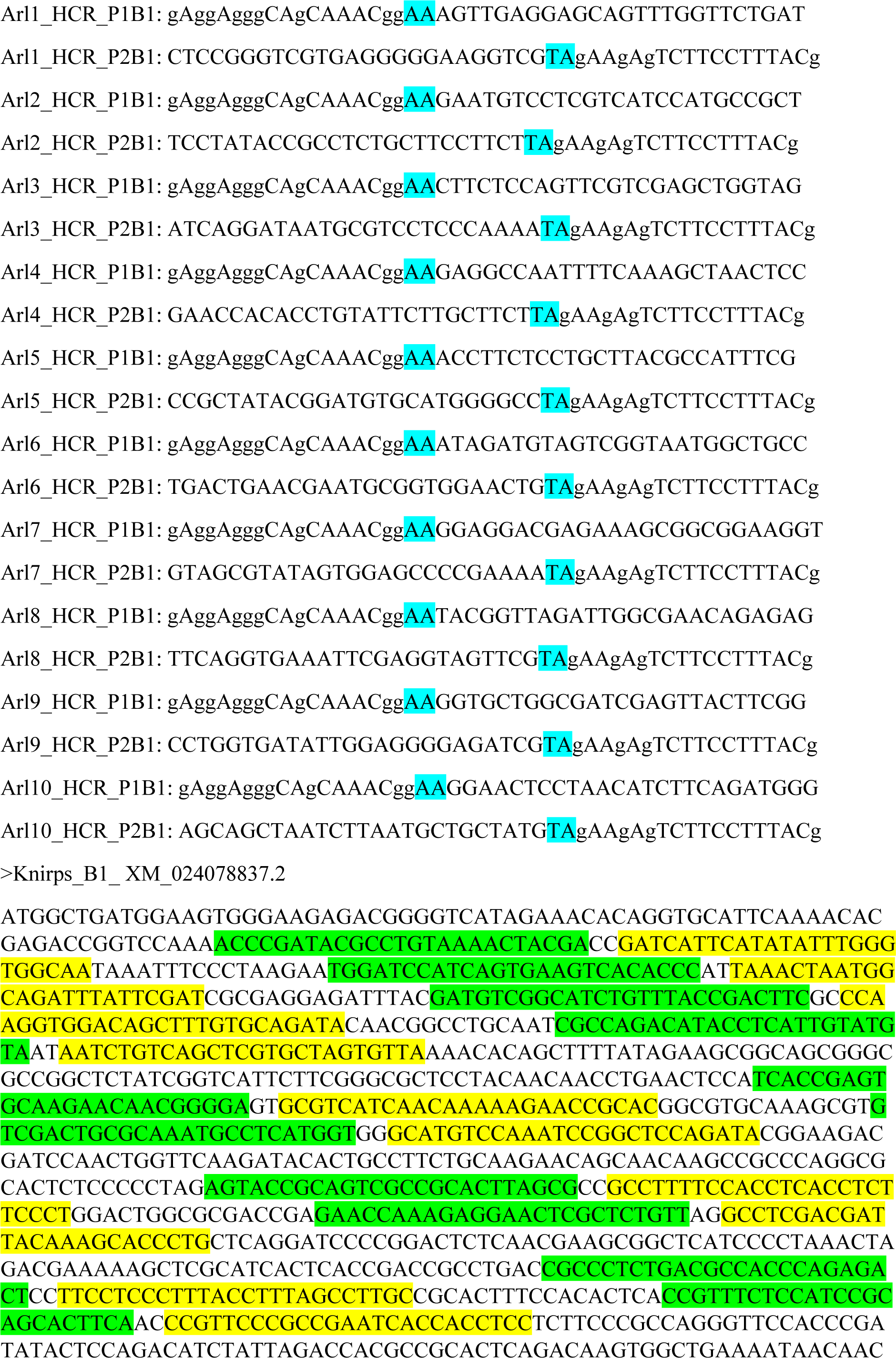
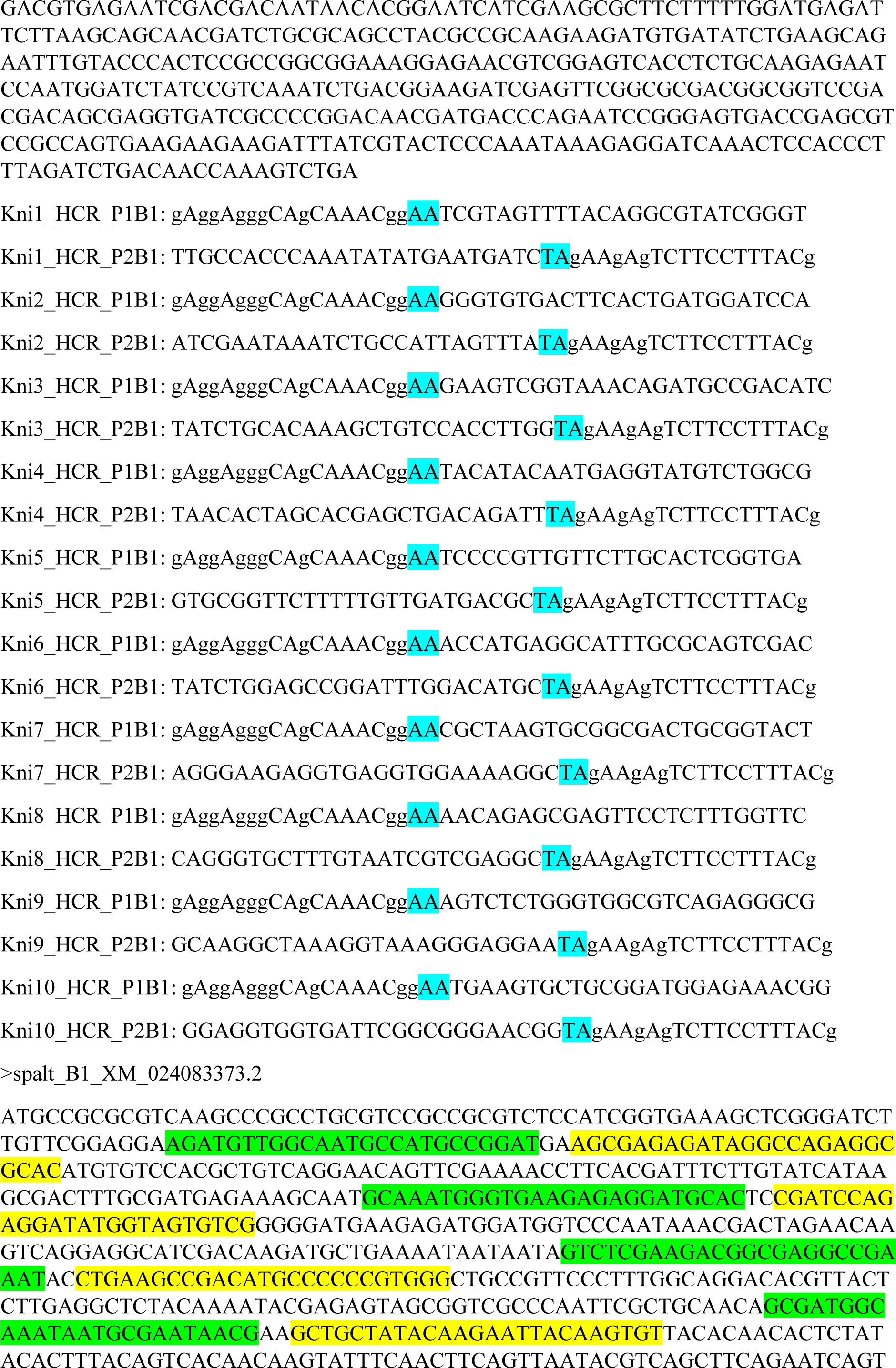
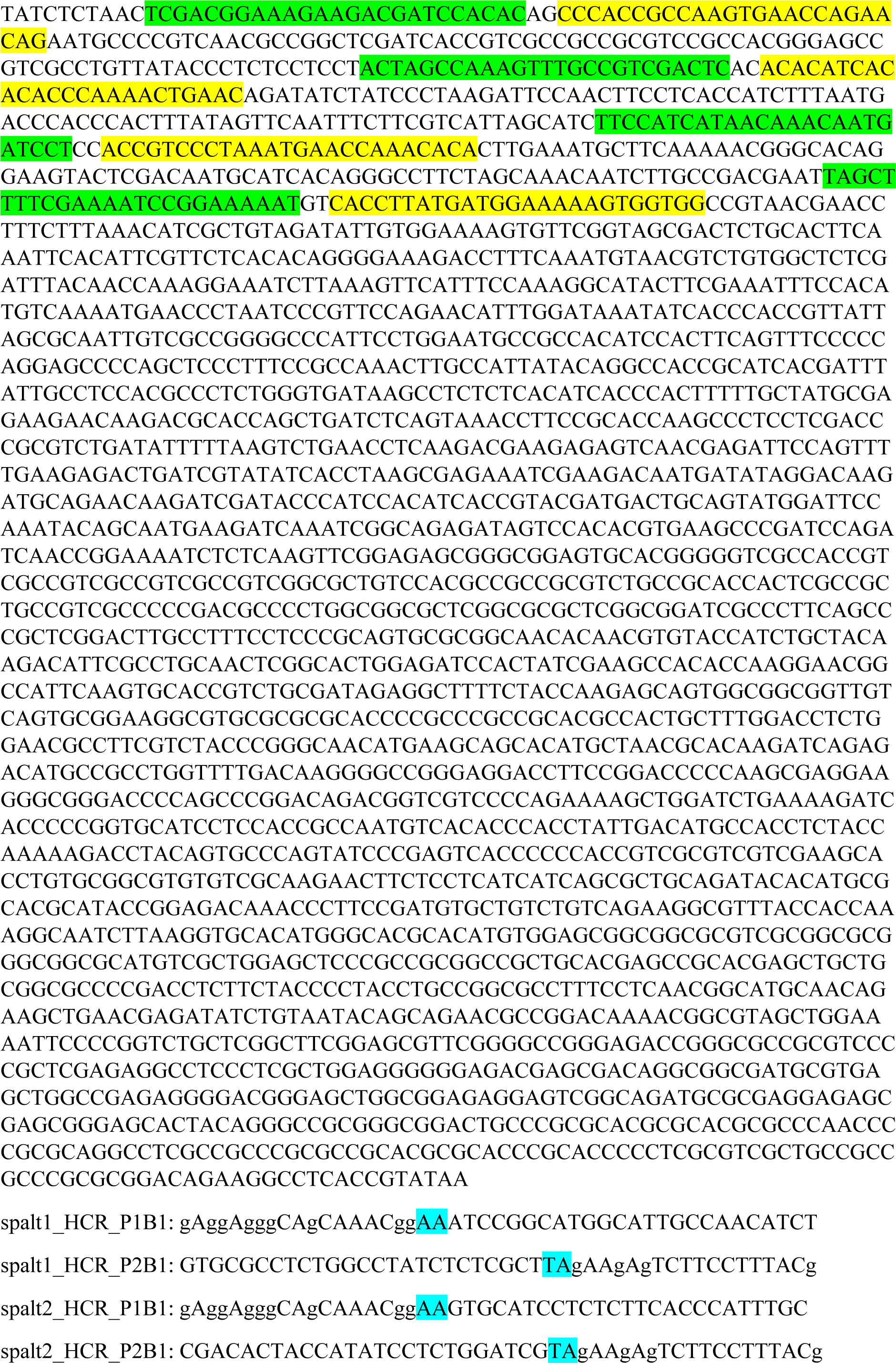
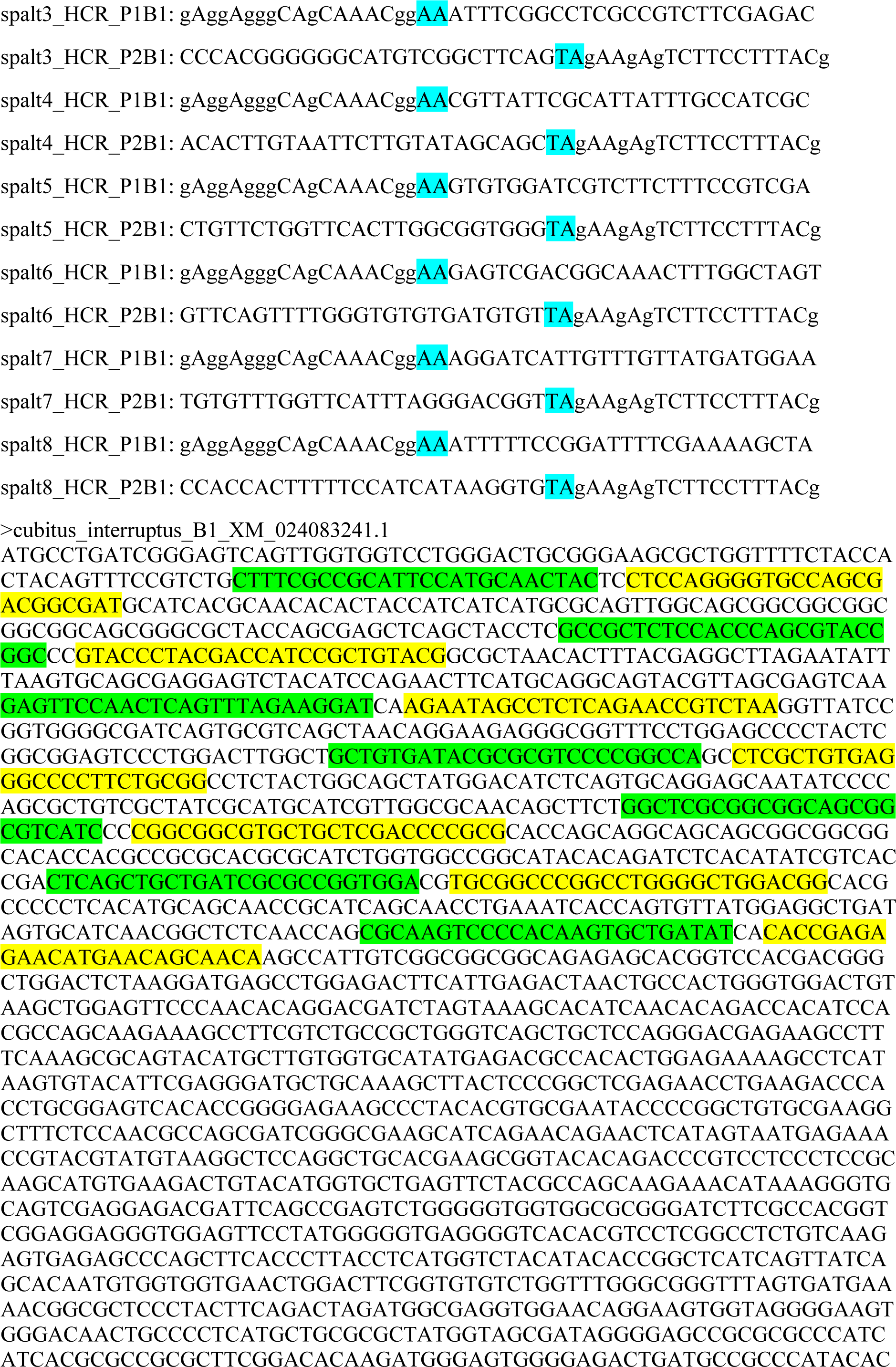
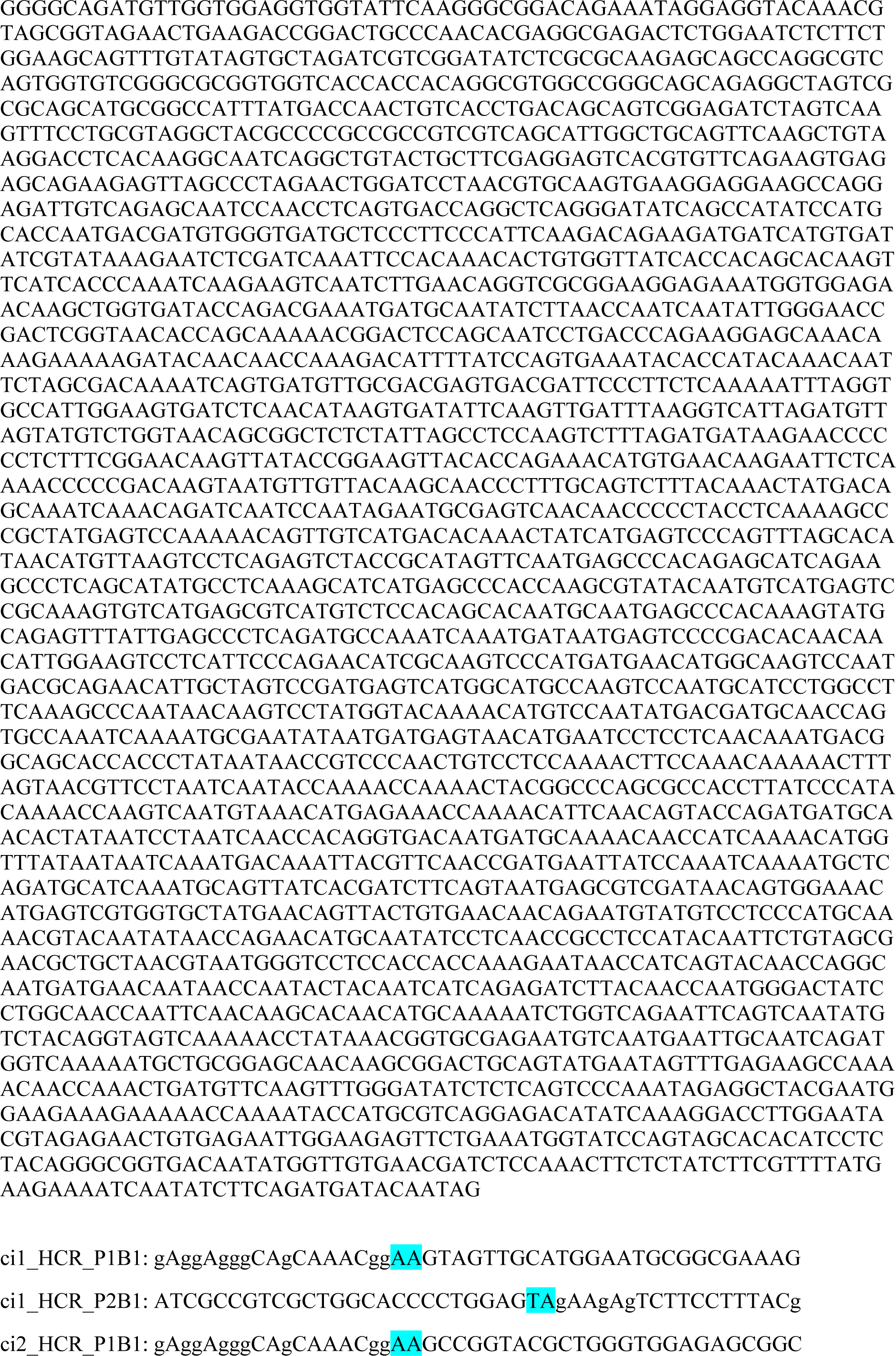
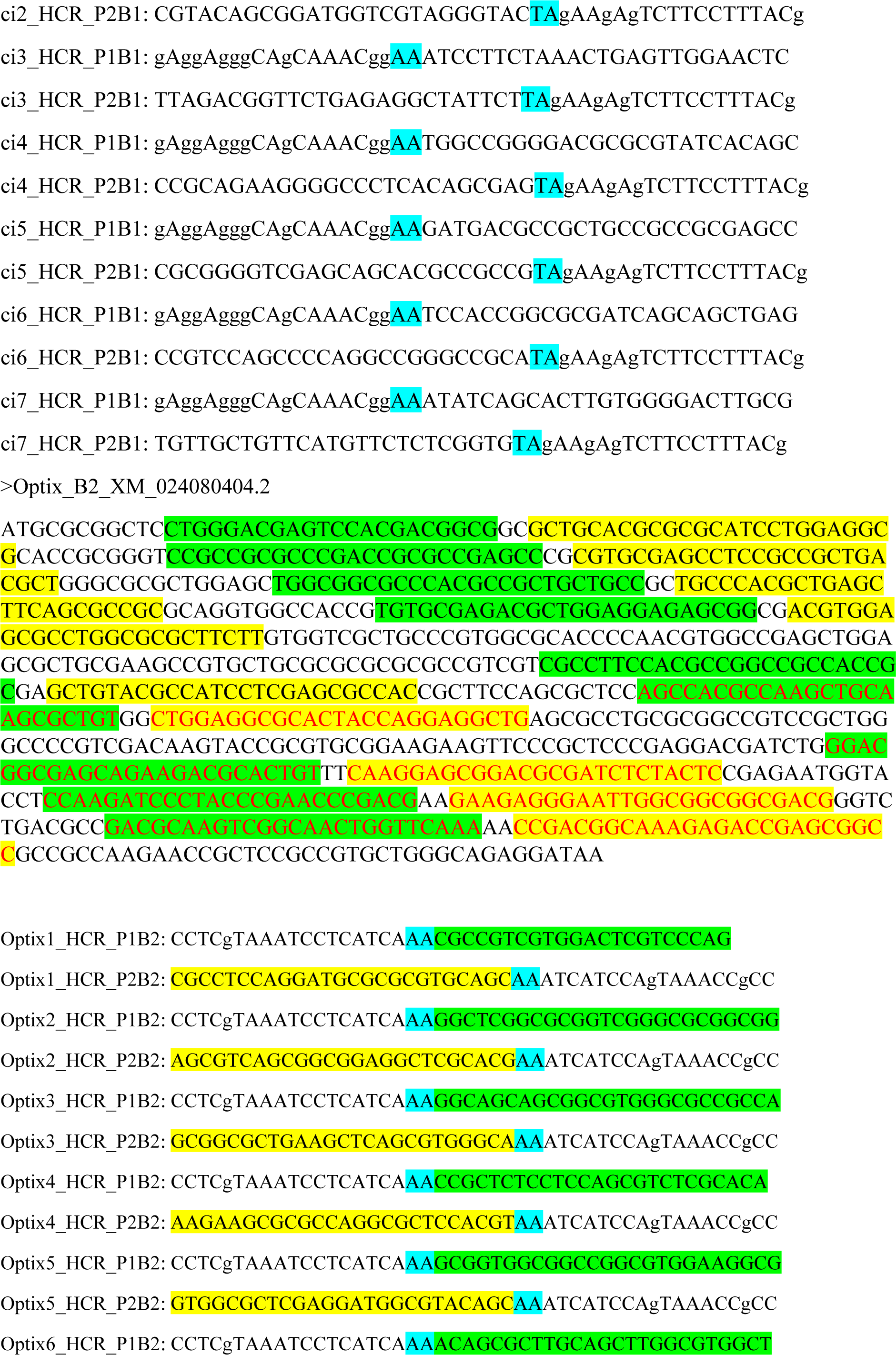
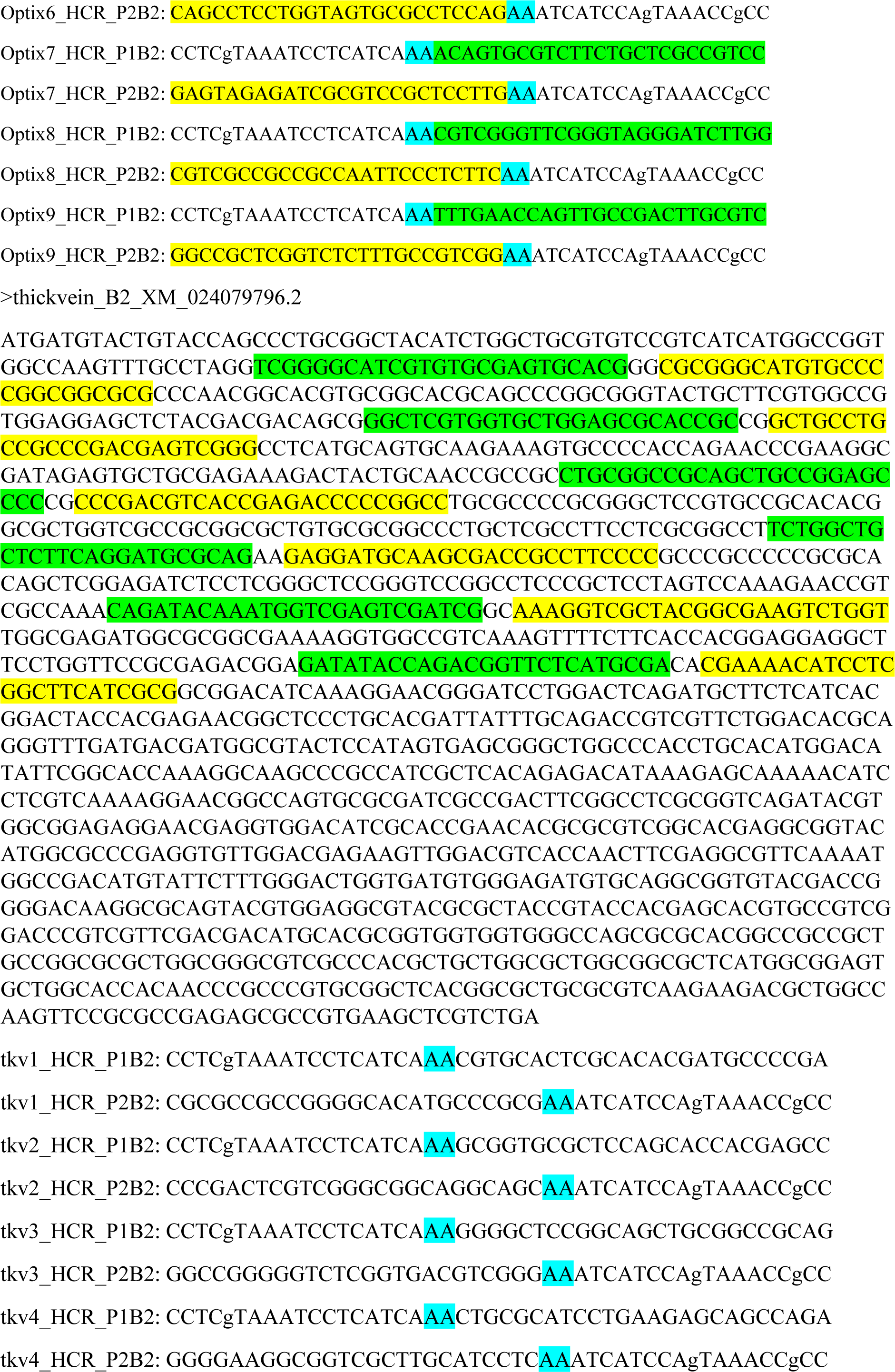
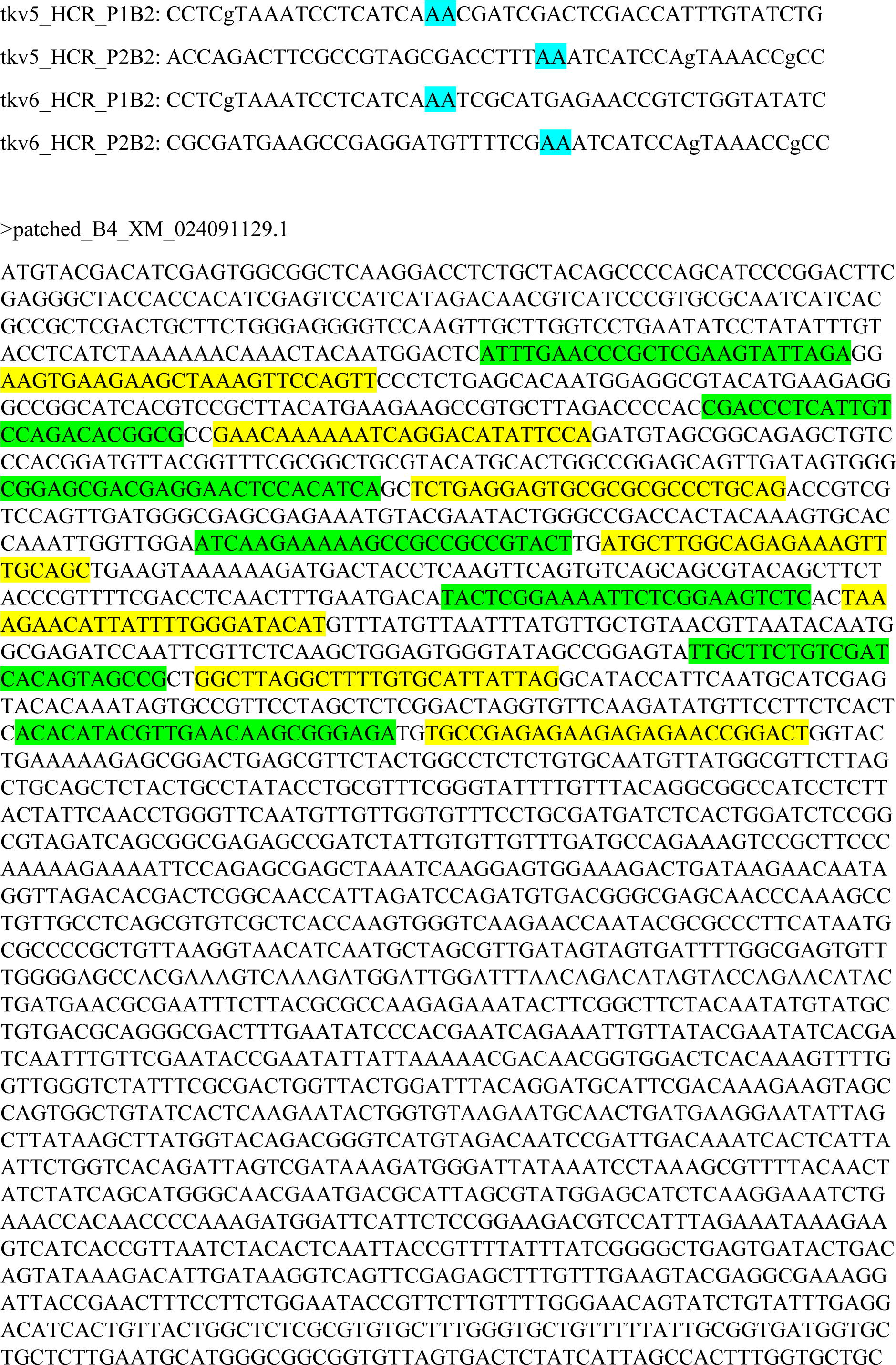
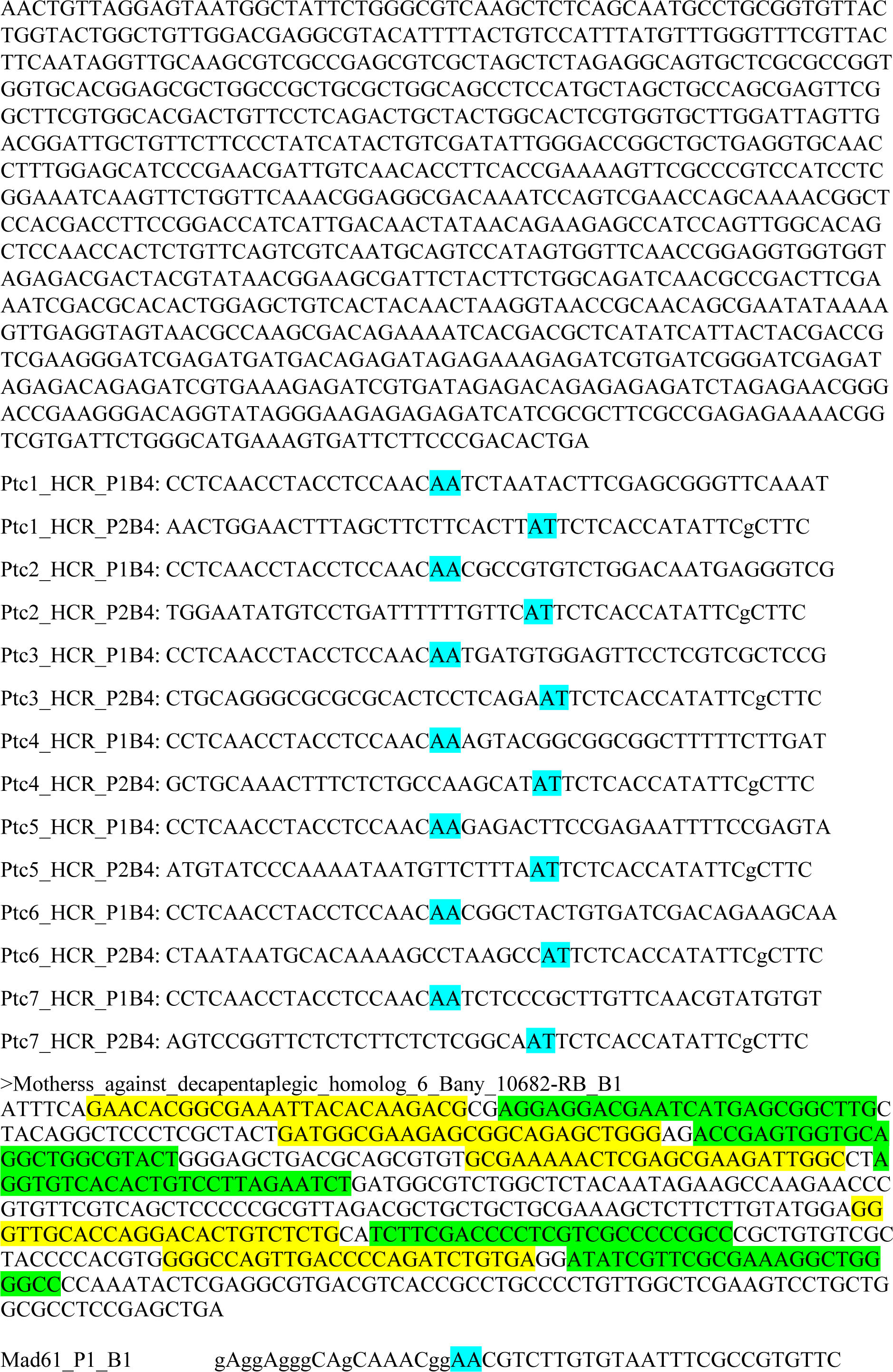
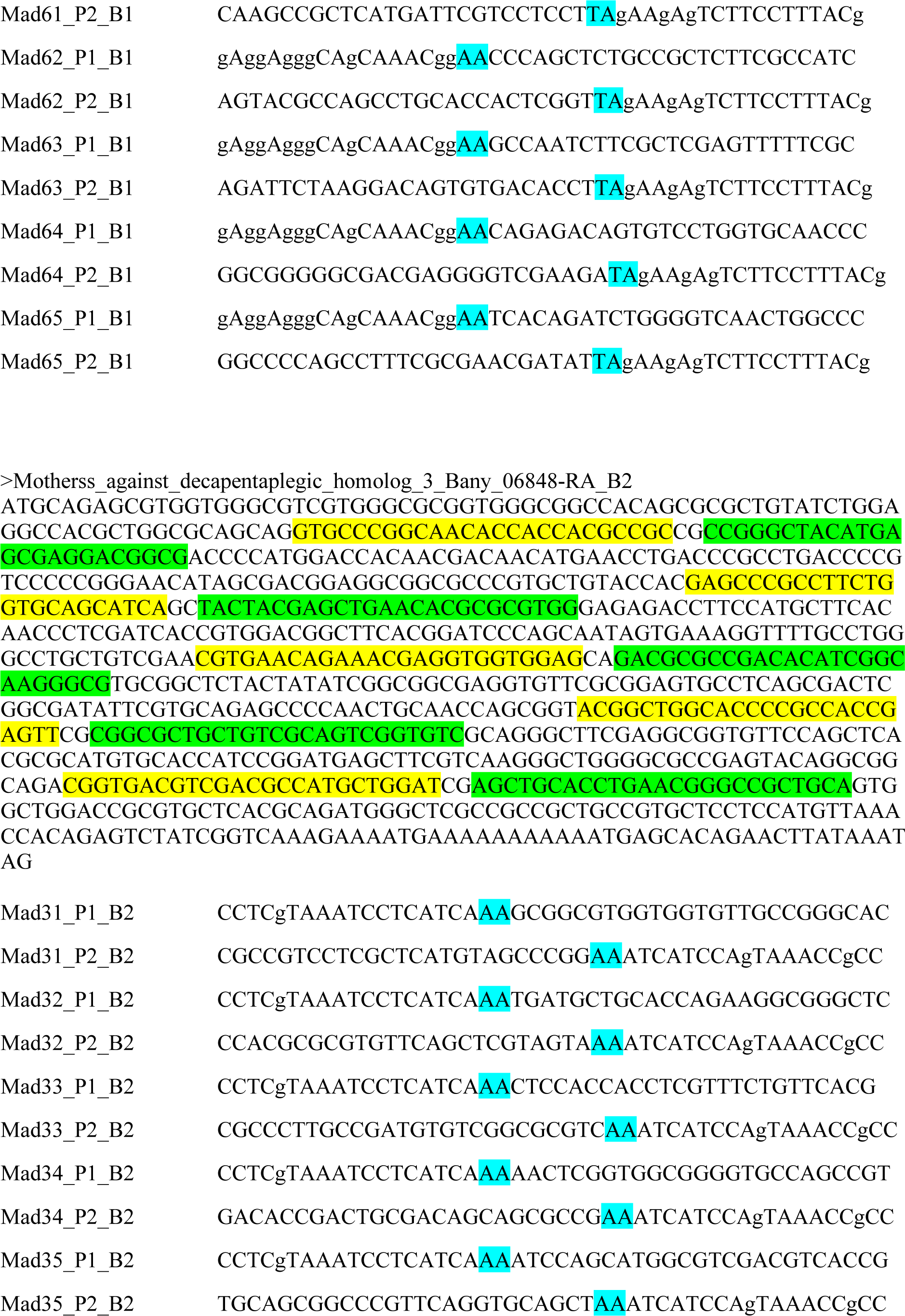
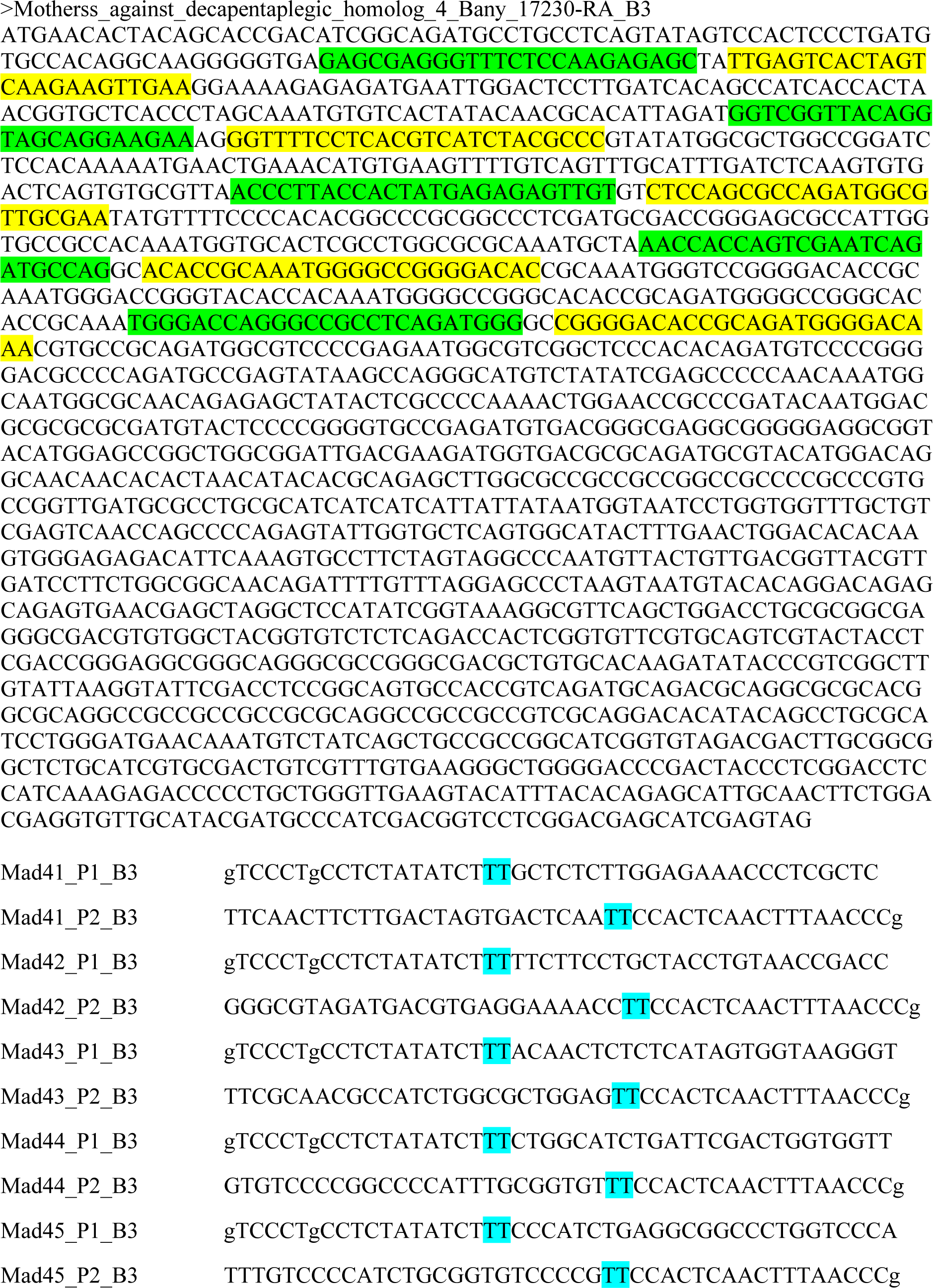

